# The process of Lewy body formation, rather than simply alpha-synuclein fibrillization, is the major driver of neurodegeneration in synucleinopathies

**DOI:** 10.1101/751891

**Authors:** Anne-Laure Mahul-Mellier, Johannes Burtscher, Niran Maharjan, Laura Weerens, Marie Croisier, Fabien Kuttler, Marion Leleu, Graham Knott, Hilal A. Lashuel

## Abstract

Parkinson’s disease (PD) is characterized by the accumulation of misfolded alpha-synuclein (α-syn) into intraneuronal inclusions named Lewy bodies (LB). Although it is widely believed that α-syn plays a central role in the pathogenesis of PD and synucleinopathies, the processes that govern α-syn fibrillization and LB formation in the brain remain poorly understood. In this work, we sought to reverse engineer LBs and dissect the spatiotemporal events involved in their biogenesis at the genetic, molecular, biochemical, structural, and cellular levels. Toward this goal, we took advantage of a seeding-based model of α-syn fibril formation in primary neurons and further developed this model to generate the first neuronal model that reproduces the key events leading to LB formation; including seeding, fibrillization, and the formation of LB-like inclusions that recapitulate many of the biochemical, structural, and organizational features of LBs found in *post-mortem* human PD brain tissues. Next, we applied an integrative approach combining confocal and correlative light-electron microscopy (CLEM) imaging methods with biochemical profiling of α-syn species and temporal proteomic and transcriptomic analyses to dissect the molecular events associated with LB formation and maturation and to elucidate their contributions to neuronal dysfunctions and neurodegeneration in PD and synucleinopathies. The results from these studies demonstrate that LB formation involves a complex interplay between α-syn fibrillization, post-translational modifications, and interactions between α-syn aggregates and membranous organelles, including mitochondria and the autophagosome and endolysosome. Furthermore, we demonstrate that the process of LB formation and maturation, rather than simply fibril formation, is the major driver of neurodegeneration through disruption of cellular functions and inducing mitochondria damage and deficits, as well as synaptic dysfunctions. Having a neuronal model that allows for unlinking of the key processes involved in LB formation is crucial for elucidating the molecular and cellular determinants of each process and their contributions to neuronal dysfunction and degeneration in PD and synucleinopathies. Such a model is essential to efforts to identify and investigate the mode of action and toxicity of drug candidates targeting α-syn aggregation and LB formation.

## Introduction

The intracellular accumulation of aggregated forms of alpha-synuclein (α-syn) in neurons and glial cells represents one of the main pathological hallmarks of Parkinson’s disease (PD) and related synucleinopathies, including dementia with Lewy bodies (DLB) and multiple system atrophy (MSA)^1^. PD and DLB are characterized by the presence of Lewy bodies (LB) and Lewy neurites (LN) in subcortical and cortical neurons, while in MSA, α-syn inclusions are mainly detected in glial cells and are referred to as glial cytoplasmic inclusions (GCIs)^2, 3^. Although it is widely believed that α-syn plays a central role in the pathogenesis of synucleinopathies, the molecular and cellular processes that trigger and govern the misfolding, fibrillization, LB formation, and spread of α-syn in the brain remain poorly understood. Furthermore, whether the processes of α-syn fibrillization and inclusion formation are protective or neurotoxic remains a subject of active investigation and debate. Addressing this knowledge gap is crucial to 1) advancing our understanding of the molecular mechanisms underpinning the pathogenesis of PD and other synucleinopathies; 2) guiding the development of preclinical models that reproduce the full-spectrum or well-defined features of the human pathology; and 3) developing more effective diagnostic and therapeutic strategies for the management and treatment of these devastating diseases.

In-depth *post-mortem* neuropathological examinations of human brains from patients with LB disorders and other synucleinopathies have revealed the existence of different sub-types of pathological inclusions that are enriched in aggregated forms of α-syn, including fibrils^3–8^. This has led to the hypothesis that aggregation of α-syn has a primary role in the formation of LBs and other α-syn pathological aggregates. However, the absence of experimental models that reproduce all the stages of LB formation and maturation has limited our ability to decipher the different processes involved in LB formation and the contribution of these processes to the pathogenesis of PD and synucleinopathies. To address this knowledge gap, it is crucial to develop cellular and/or animal models that not only produce α-syn aggregates but also recapitulate the process of LB formation at the biochemical, structural, and organizational levels. One advantage of cellular models is that they offer unique opportunities to apply advanced imaging and omics approaches to elucidate in great detail the key events associated with α-syn aggregation and LB formation and maturation and to correlate these events with alteration in cellular pathways and functions/dysfunctions.

Until recently, the great majority of cellular models of synucleinopathies were based on overexpression of WT or mutant α-syn alone, with other proteins, or coupled to treatment with other stress inducers (e.g., toxins, proteasome inhibitors)^9–19^. In many cases, these conditions were sufficient to induce α-syn accumulation and aggregation. However, very few studies investigated the extent to which these aggregates reproduce the cardinal biochemical, structural, and organizational features of LBs and other pathological states of α-syn in PD brains. The formation of LB-like structures is usually assessed using a limited number of methods to indirectly assess the aggregation properties of α-syn (e.g., proteinase K resistant^9, 16^ and binding to the amyloid-specific dye, thioflavin S^13^) and immunoreactivity for selected LB markers^10, 12–15, 17, 19^, such as p62, ubiquitin (Ub), and phosphorylated α-syn (pS129), to define their LB-like properties. When electron microscopy (EM) was used, it clearly demonstrated that the α-syn inclusions do not share the compositional complexity and morphological features of the LBs, despite their immunoreactivity to the standard LB markers^13, 20, 21^. These findings suggest that the great majority of α-syn cellular models reproduce some aspects of α-syn aggregation or fibril formation, but do not recapitulate all of the key events leading to LB formation and maturation.

Recently, it was shown that exogenously adding pre-formed fibrils of α-syn at nanomolar concentration can act as a seed to initiate the misfolding and aggregation of endogenous α-synuclein in both cellular^22, 23^ and animal models^24^. This neuronal seeding model is the first model that shows the formation and accumulation over time of long filamentous aggregates. Although this seeding model enables the induction of α-syn fibrillization in neurons, in the absence of α-syn overexpression, the transition from fibrils to LB-like structures has not been reported, even though these aggregates are immunoreactive for the standard LB makers (e.g., pS129 α-synuclein, p62, and ubiquitin). These observations suggest that the neuronal seeding model, as initially developed, is a suitable model for investigating the processes involved in fibrils formations but not LB formation.

Given that LBs have been consistently shown to have complex composition and organization and usually contain other non-proteinaceous material (lipids)^25, 26^ and membranous organelles^8, 27–32^, we hypothesized that the transition from fibrils to LB might require time and could be driven by post-translational modifications and structural rearrangements of the fibrils. By extending the characterization of this neuronal seeding model from 11–14 days to 21 days, we were eventually able to observe the transition from fibrils to α-syn rich inclusions that recapitulate the biochemical, morphological, and structural features of the *bona fide* human LBs, including the recruitment of membranous organelles and accumulation of phosphorylated and C-terminally truncated α-syn aggregates^33–38^.

By employing an integrated approach using confocal and CLEM imaging, as well as quantitative proteomic and transcriptomic analyses, we were able to investigate with spatiotemporal resolution the biochemical and structural changes associated with each step involved in LB biogenesis, from seeding to fibrillization to LB formation and maturation. Our ability to uncouple the different stages of LB formation in this model provided unique opportunities to investigate the cellular and functional changes that occur at each stage, thus paving the way for elucidating how the different events associated with LB formation and maturation contribute to α-syn-induced toxicity and neurodegeneration. Our work shows that the process of LB formation and maturation, rather than simply α-syn fibril formation, is a major contributor to α-syn-induced toxicity and neurodegeneration in PD and synucleinopathies. We believe that this new model of LB formation can be further developed and used as a powerful platform to further investigate the molecular determinants and cellular pathways that regulate the stages of LB formation and maturation and to screen for therapeutic agents based on the modulation of these pathways.

## Results

### At early stage, α-syn seeded aggregates resemble *bona fide* human LBs histochemically but not at the structural and organizational levels

To investigate the mechanisms of LB formation and maturation, we initially took advantage of a cellular seeding-based model of synucleinopathies developed by Virginia Lee and coworkers^22, 23^. As a first step, we sought to determine to what extent this model reproduces LB formation. Extracellular sonicated α-syn pre-formed fibrils (PFFs, length ∼50–100 nm) (Figure S1A-E) were added at a nanomolar concentration (70 nM) to primary neuronal cultures (Figures 1A). Once internalized into the neurons, the PFFs serve as seeds and induced endogenous α-syn to misfold and form intracellular aggregates in a time-dependent manner (Figures 1B and S1F). To establish the spatio-temporal features of α-syn seeded aggregates formed in neurons, we used ICC (Figures 1C-J and S3A-E) combined with high content imaging analysis (HCA), which allows quantitative assessment of the level of α-syn-pS129 in MAP2 positive neurons (Figure S3F-I). The epitopes of the antibodies used in our study are summarized in Supplemental Figure 2.

**Figure 1.**
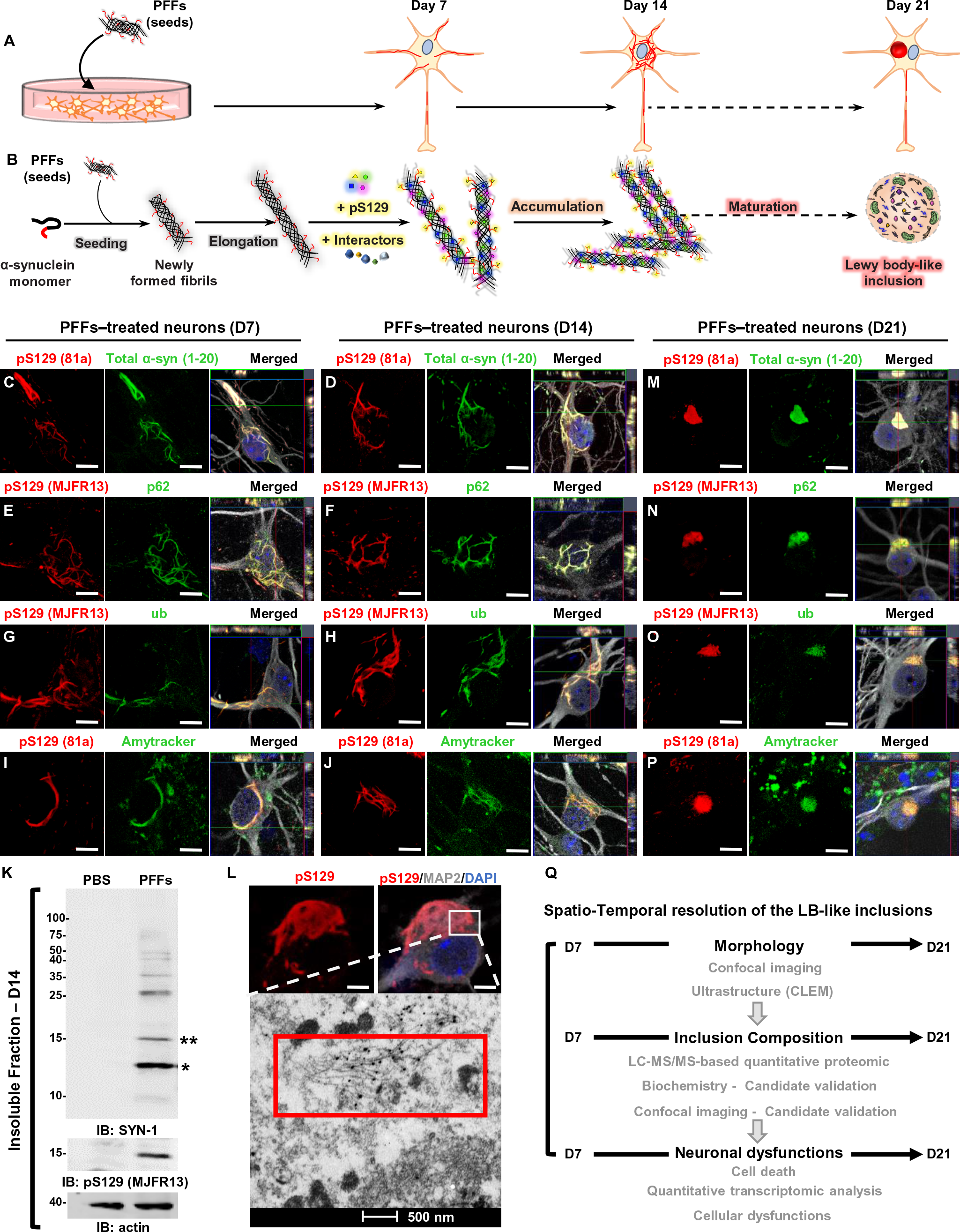
At early stage, α-syn-seeded fibrillar aggregates share the immunohistochemical but not the structural and morphological features of the *bona fide* human LBs. **A.** Seeding model in primary hippocampal neurons. 70 nM of mouse sonicated PFFs were added to neurons at DIV 5 (days *in vitro*). Control neurons were treated with PBS used to prepare PFFs. Following a defined period of 4 days (D4) during which no seeded aggregates are detected, α-syn aggregates first appear in the neuronal extension (between D4 to D7) before being detected in the neuronal cell bodies (D7–D14) where they eventually accumulate (D14 to D21).
**B.** The formation of α-syn aggregates in the context of the neuronal seeding requires a sequence of events starting with 1) the internalization and cleavage of PFF seeds (D0–D1) followed by 2) the initiation of the seeding by the recruitment of endogenous α-syn (D0–D4), 3) fibril elongation along with the incorporation of PTMs, such as phosphorylation at residue S129 (D4–D14), and 4) formation of large filamentous aggregates (D7–D14).
**C-J, M-P.** Temporal analysis of α-syn aggregates formed at D7 (C, E, G, I), at D14 (D, F, H, J), and at D21(M-P) in WT neurons after addition of α-syn PFFs. Aggregates were detected by ICC using pS129 (MJFR13 or 81a) in combination with total α-syn (epitope: 1-20) (**C-D, M**), p62 (**E-F, N**) or ubiquitin (**G-H, O**) antibodies. Aggregates were also stained with the Amytracker dye (**I-J, P**). Neurons were counterstained with microtubule-associated protein (MAP2) antibody, and the nucleus was counterstained with DAPI staining. Scale bars = 10 μm.
**K.** WB analysis of α-syn seeded-aggregates formed 14 days after adding mouse PFFs to WT neurons. Control neurons were treated with PBS. After sequential extractions, the insoluble fractions of neuronal cell lysates were analyzed by immunoblotting. Total α-syn, pS129, and actin were detected by SYN-1, pS129 (MJFR13), and actin antibodies, respectively. Monomeric α-syn (15 kDa) is indicated by a double asterisk; C-terminal truncated α-syn (12 kDa) is indicated by a single asterisk; the higher molecular weights (HMWs) corresponding to the newly formed fibrils are detected from 25 kDa to the top of the gel.
**L.** At D14, neurons were fixed, and ICC with a pS129 (81a) antibody was performed and imaged by confocal microscopy (upper images). Scale bar = 5 μm. The selected neuron was embedded and cut by an ultramicrotome. Serial sections were examined by EM (bottom image). Immunogold labeling allowed the detection of long filaments in the neuronal cell body that appeared as tightly packed bundles of fibrils that were positive for pS129-α-syn. Scale bar = 500 nm.
**Q.** In this study, we employed an integrated approach using advanced techniques in confocal imaging, EM, proteomics, transcriptomic, and biochemistry to investigate, at the spatio-temporal resolution (from D7 to D21), how the seeded intracellular aggregates recapitulate the Lewy body-like inclusion pathology at the morphological, biochemical, structural, and biological levels.

Consistent with previous studies, we observed that PFF-treated neurons only started to exhibit α-syn-pS129 positive inclusions 4 days after the addition of α-syn seeds (Figure S3A). These rare aggregates were exclusively detected in neurites (Figure S3A). After 7 days of treatment (D7), we observed an accumulation of filamentous-like α-syn aggregates in neurites with a limited number of cell-body inclusions (Figures 1C and S3B, G-I). At 14 days (D14), the number of filamentous aggregates continued to significantly increase in neurites but also in the neuronal cell bodies where they were distributed in the vicinity of the nucleus (Figures 1D and S3C, G-I).

The newly formed aggregates observed at D7 and D14 were all positive for the two well-established LB markers, p62 (Figure 1E-F) and ubiquitin (Ub) (Figure 1G-H)^39^ and were also stained by the Amytracker tracer, a fluorescent dye that binds specifically the *β*-sheet structure of amyloid-like protein aggregates (Figure 1I-J). Although previous studies have suggested the formation of specific morphologies (strains) of α-syn aggregates in seeding models, our ICC data revealed the formation of aggregates that are heterogeneous in size, shape, and subcellular distribution in the same population of PFF-treated neurons (Figures 1C-J and S3B-C). Biochemical analysis of the insoluble fraction of the PFF-treated neurons confirmed the accumulation of both truncated and high molecular weight (HMWs) α-syn species that were positively stained with a pan-synuclein antibody (SYN-1) and with an antibody against pS129 (Figures 1K and S7). Interestingly, these insoluble fractions displayed a potent seeding activity in neuronal primary culture (Figure S4). Compared to recombinant PFFs, we did not observe differences in the morphology and the subcellular distribution of the seeded-aggregates formed after addition of “native aggregates” (Figures S3C and S4B). These findings demonstrate that the newly formed pS129 positive α-syn filamentous aggregates contain α-syn seeding competent species, most likely in a fibrillar species.

To determine if the newly formed aggregates share the structural and morphological properties of the *bona fide* LBs observed in PD brains, we performed Electron Microscopy (EM) and immunogold labeling studies against α-syn pS129 at D14. As shown in Figure 1L, the newly formed aggregates in neurons appear as loosely organized uniform filaments (Figure 1L, bottom panel). Interestingly, at this point, we did not observe significant lateral association or clustering of the fibrils in the form of inclusions. Although previous studies have referred to the α-syn aggregates formed at 11–14 days post-treatment with PFFs as LB-like inclusions, our ICC and EM studies, which are consistent with previous studies using the same model^23, 40, 41^ and tools (e.g., antibodies and LB markers), clearly demonstrate the formation of predominantly α-syn fibrils, rather than LB-like inclusions. LBs are highly organized round structures that are composed of not only α-syn fibrils but also other proteins^42^, lipids^25, 26^, and membranous organelles, including lysosomal structures and mitochondria^8, 28, 29, 31, 43^. The absence of such structures at D14, even though the fibrillar aggregates are immunoreactive for the classical markers used to define LBs, pS129, ubiquitin, and p62, suggests that these markers cannot be used to accurately distinguish between α-syn fibrillar aggregates and mature LBs.

### At late stage, α-syn seeded fibrillar aggregates rearrange into inclusions that morphologically resemble to the *bona fide* human LBs

We hypothesized that the rearrangement and transition of the newly formed α-syn fibrils into LB-like inclusions might require more time. Therefore, we sought to investigate the structural and morphological properties of the newly formed aggregates over an extended period, up to 21 days (D21) post-treatment with PFFs (DIV 28). Interestingly, at D21, ICC combined with a HCA approach showed a significant increase in the number of aggregates detected in the neuronal cell bodies between D14 and D21 (Figure S3F-H). Furthermore, we observed major changes in the morphology of the newly formed aggregates. The seeded aggregates, up to 14 days of formation, were exclusively detected as long filamentous-like structures located either in the neurites and/or the neuronal cell bodies (Figure S3B-C). However, at D21, we identified three different morphologies; filamentous (∼30%), ribbon-like (∼45%), and round LB-like inclusions (∼ 22%) (Figure S3D-E). The formation of these types of aggregates in seeded neurons has not been reported, as previous studies limited their analysis up to D14. The α-syn rounded inclusions observed at D21 were all positive for p62, Ub, and the amyloid-specific dye (Amytracker) (Figure 1M-P). Furthermore, we also showed the presence of several classes of lipids in the pS129-positive inclusions using fluorescent probes (Figure S3 J-O). This finding is in line with several studies which have previously documented the presence of lipids within LBs and α-syn pathological inclusions observed in *post-mortem* PD human brain tissues^25, 26, 44–47^.

Finally, the LB-like inclusions were exclusively detected in the vicinity of the nucleus (Figures 1M-P and S3D-E). After D14, no further significant accumulation of α-syn aggregates was observed in the neurites (Figure S3G). Altogether, our data suggest that α-syn fibrillar aggregates initially formed in the neuronal extensions (D4) are transported over time (D7–D21) to the perinuclear region, where they eventually accumulate as LB-like inclusions (D21).

### The process of inclusion formation is accompanied by the sequestration of lipids, organelles and endomembranes structures

The transition from filamentous aggregates into ribbon and round-like aggregates suggests that the newly formed fibrils undergo major structural changes over time. To determine if these changes are associated with the evolution and maturation of α-syn fibrils into LBs, we next turned to correlative light and electron microscopy (CLEM) to characterize the ultrastructural properties of the seeded α-syn aggregates formed in neurons after 7, 14, or 21 days post-treatment with WT PFFs (Figures 1Q, 2 and S5A). As shown in Figure 2, the CLEM data revealed a marked reorganization of the newly formed α-syn fibrils over time. Indeed, at D7, EM images of neurites positive for pS129-positive α-syn seeded aggregates showed predominantly single fibril filaments distant from each other (Figures 2Aa). Aligned and randomly ordered fibrils were observed in the same neurite (Figures 2Aa). In comparison, in neurite negative for α-syn seeded aggregates, we observed the classical organization of the microtubules that appear as aligned bundle of long filaments parallel to the axis of the neurites (Figures 2Ab). In-depth measurements of the diameters of the filament-like structures observed in neurites confirmed that the newly formed α-syn fibrils exhibited an average diameter of 11.94 ± 3.97 nm (standard deviation, SD) which was significantly smaller than the average diameter of the microtubules (18.67 ± 2.94 nm) (Figure 2Ab, inset). Moreover, the range of fibril lengths (from 500 nm to 2 μM per slide section) (Figure 2A) confirmed that the fibrils detected in the cytosol were not simply internalized α-syn PFFs (size ∼40-120 nm, Figure S5B) but rather newly formed fibrils resulting from the seeding and fibrillization of endogenous α-syn. At D7, although most of the newly formed α-syn fibrils were observed in the neuritic extensions, pS129-positive α-syn aggregates were also detected in the cell bodies of few neurons (Figure 1C-I).

**Figure 2.**
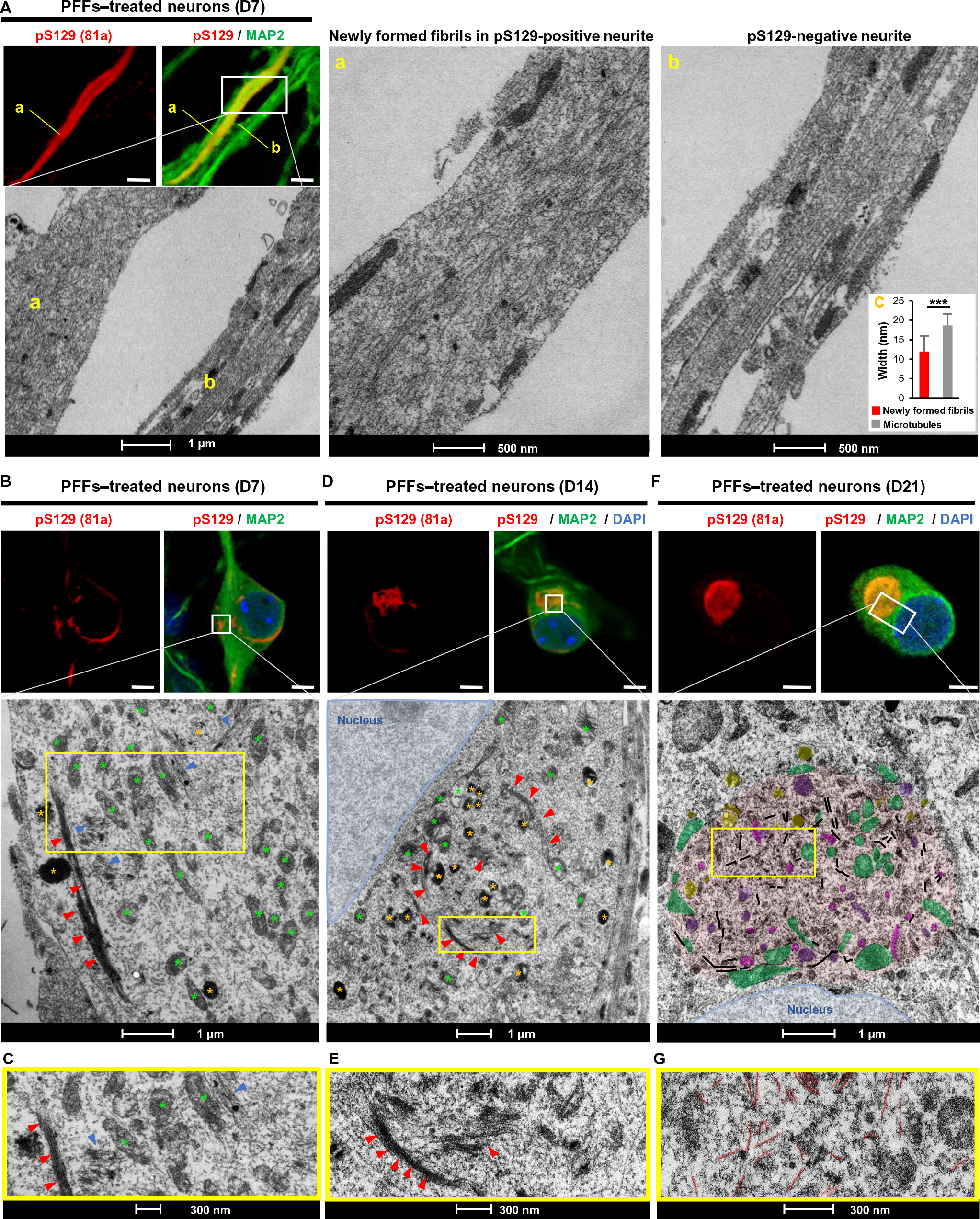
Formation and maturation of LB-like inclusions require the lateral association and the fragmentation of the newly formed α-syn fibrils over time. PFFs were added for 7, 14, and 21 days to the extracellular media of primary hippocampal neurons plated on dishes with alpha-numerical searching grids imprinted on the bottom, which allowed easy localization of the cells. At the indicated time, neurons were fixed, ICC against pS129-α-syn was performed and imaged by confocal microscopy (**A, B, D, F** top images; scale bar = 10 μm), and the selected neurons were embedded and cut using a ultramicrotome. Serial sections were examined by EM (**A, B, D**, **F** bottom images). **Aa**. Neurite with pS129-positive newly formed α-syn fibrils. **Ab**. Neurite negative for pS129 staining. **Ab, inset.** Graph representing the mean ± SD of the width of the microtubules compared to the newly formed fibrils at D7 (a minimum of 120 microtubules or newly formed fibrils were counted). Measurements confirmed that the width of newly formed α-syn fibrils of 11.94±3.97 nm (SD) is significantly smaller than the average width of the microtubules of 18.67±2.94 nm (SD). p<0.001=*** (student t-test for unpaired data with equal variance), indicating that this parameter can be used to discriminate the newly formed fibrils from the cytoskeletal proteins. Autophagolysosomal-like vesicles are indicated by a yellow asterisk (**D**) or colored in yellow (**F**), and mitochondrial compartments are indicated by a green asterisk (**D**) or colored in green (**F**). Accumulation of endomembranous compartment at a late stage (D21) are colored in pink (endosomes-like vesicles) and purple (lysosomes-like vesicles), respectively. Nuclei are highlighted in blue. **C-G.** Representative images at higher magnifications corresponding to the area indicated by a yellow rectangle in EM images shown in **B**, **D**, and **F** respectively. **A, B, D, F.** Scale bar = 1 μm; **Aa and Ab.** Scale bar = 500 nm; **C-G.** Scale bars = 300 nm.

CLEM showed, in the same neuronal cell body, α-syn fibrils randomly organized (Figure 2B-C, blue arrows) or ordered as long tracks of parallel bundles of filaments (Figure 2B-C, red arrows). At D14, most α-syn filaments were reorganized into laterally associated and tightly packed bundles of fibrils (Figure 2D-E, red arrows).

In some inclusions, these laterally associated bundles were seen to start to associate with or encircle organelles, including mitochondria, autophagosomes, and others endolysosomal-like vesicles (Figure 2D-E). Single fibrils, loosely distributed in the neuronal cell body, were also detected close to these organelles (Figure 2D-E).

The sequestration of organelles and membranous structures by α-syn fibrillar aggregates seems to gradually increase over time between D14 and D21 (Figures 2D-F and S5F). At D21, both the ribbon-like aggregates (Figures 3A-B, highlighted in red and S6A) and the LB-like inclusions (Figures 2F, 3C-D, highlighted in red and S6B) were composed of α-syn filaments (highlighted in black) but also of a high number of mitochondria (highlighted in green) and several types of vesicles, such as autophagosomes, endosome and lysosome-like vesicles (highlighted respectively in yellow, pink and purple) (Figures 2F, 3A-D and S6A-D). Interestingly, the mitochondria were either sequestered inside the inclusions or organized at the periphery of the inclusions. The presence of these organelles inside the LB-like inclusions was confirmed by ICC using specific markers for the late endosomes/lysosomes (Figure 3E, LAMP1), the mitochondria (Figure 3G, Tom20) and the autophagosome vesicles (Figure 1M, p62, and 3F, LAMP2A). BiP protein localized to the endoplasmic reticulum was also partially colocalizing to the LB-like inclusions (Figure 3H). Our data strongly suggest that the process of inclusion formation is not only driven by the mechanical assembly of newly formed α-syn fibrils but seems to be accompanied by the sequestration or active recruitment of other proteins, membranous structures, and organelles over time. This is in line with previous studies showing that LBs from PD brains are not only composed of α-syn filaments and proteinaceous material^42, 48, 49^ but also contain a mixture of vesicles, mitochondria, and others organelles ^8, 27–31, 43^ as well as lipids^25, 26^.

**Figure 3.**
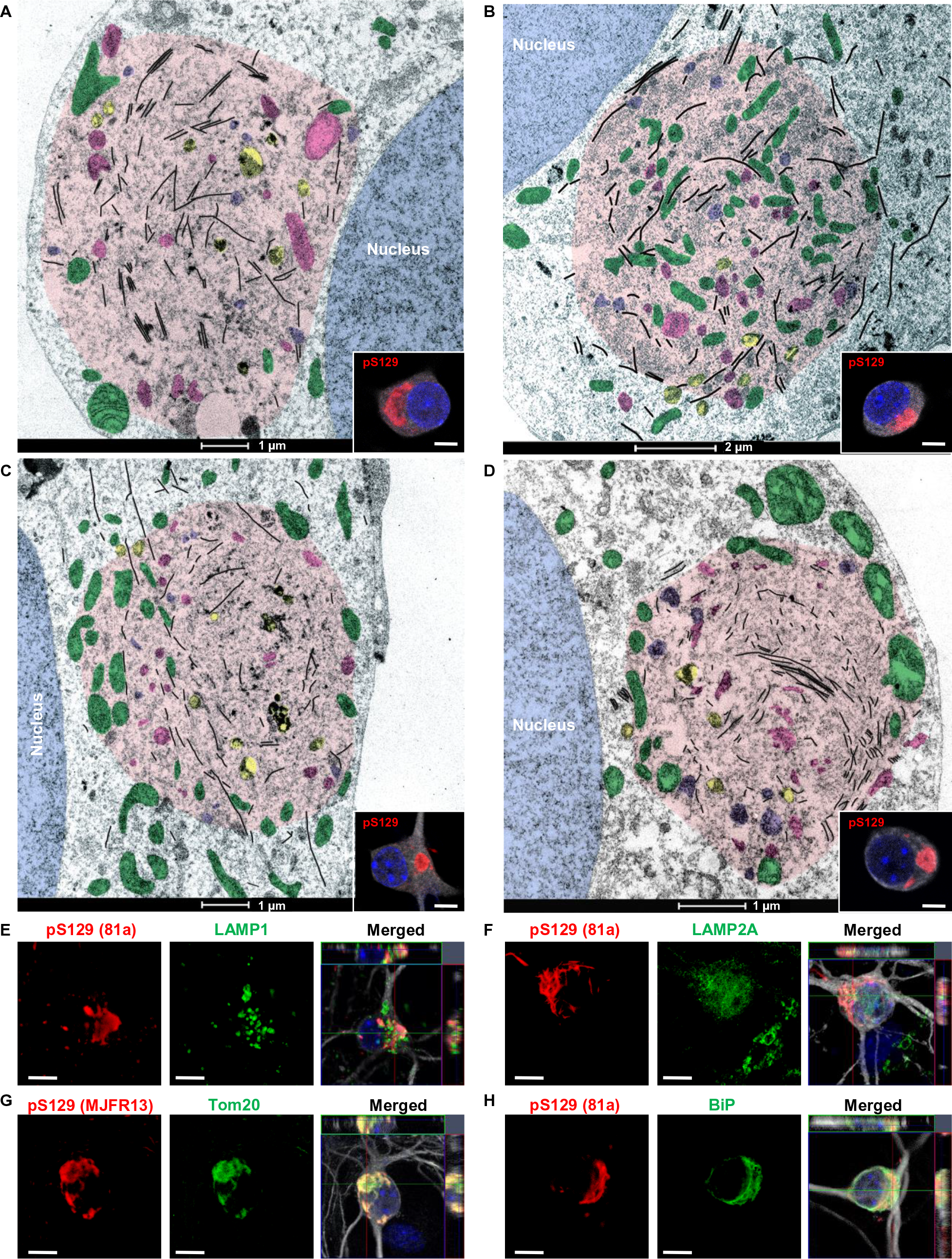
Maturation of α-syn-seeded aggregates into LB-like inclusions is accompanied by the sequestration of organelles and the endomembrane system over time. **A-D**. At D21, PFF-treated neurons were fixed, ICC against pS129-α-syn was performed and imaged by confocal microscopy (**A-D**, inset images; scale bar = 5 μm), and serial sections of the selected neurons were examined by EM. Representative images of LB-like inclusions which exhibited a ribbon-like morphology (**A-B**) or compact and rounded structures (**C-D**). A high number of mitochondria (highlighted in green) and several types of vesicles, such as autophagosomes, endosome, and lysosome-like vesicles (highlighted respectively in yellow, pink and purple) were detected inside or at the edge of these inclusions. **A, C-D.** Scale bar = 1 μm; **B.** Scale bar = 2 μm.
**E-H**. LB-like inclusions were detected by ICC using pS129 (MJFR13 or 81a) in combination with LAMP1 (late endosome/lysosome marker) (**E**), LAMP2A (autophagolysosome marker) (**F**), Tom20 (mitochondria marker) (**G**), or BiP (ER marker) (**H**) antibodies. Neurons were stained with microtubule-associated protein (MAP2) antibody, and the nucleus was counterstained with DAPI. Scale bars = 10 μm.

### Formation and maturation of LB-like inclusions require the lateral association and the fragmentation of the newly formed α-syn fibrils over time

The lateral association of the newly formed fibrils at D14 might represent an early stage in the process of packaging fibrillar aggregates into LB-like inclusions. Also, the length of the newly formed fibrils significantly decreased over time during the evolution of the inclusions (Figure S5G-J). At D7, the neurites contained long filaments of α-syn reaching up to 2 μm in length (Figures 2Aa, 2B-C and S5G, J). Strikingly, at D14, the size of α-syn fibrils was significantly reduced to an average length of 400 nm (Figures 2D-E and S5H, J). This was even more obvious in the compact inclusions observed in neurons treated for 21 days, in which only very short filaments were detected with an average size of 300 nm (Figures 2F-G and S5F, I-J). This suggests that the lateral association of the newly formed fibrils and their packing into higher-organized aggregates might require fragmentation of α-syn into shorter filaments. The presence of C-terminally truncated species of α-syn was confirmed by Western blotting (WB), as evidenced by the detection of the 12 kDa band by the pan α-syn antibody (SYN-1, epitope 91-99) but not by the antibody against pS129 (Figure S7). This is in line with our recent findings showing that specific post-fibrillization C-terminal cleavages serve as key regulators in the processing of α-syn seeds and the growth of newly formed α-syn fibrils in the neuronal seeding model^33^.

Altogether, our findings suggest that the formation of α-syn inclusions, in the context of this neuronal seeding model, requires a sequence of events starting with 1) the internalization of PFF seeds, followed by 2) the initiation of the seeding, 3) fibril elongation and post-fibrillization modification (phosphorylation at S129 and ubiquitination), 4) the fragmentation of the newly formed fibrils, which facilitates 5) the structural reorganization of the fibrils, their lateral assembly, and their packing into higher-order aggregates along with 6) the recruitment and sequestration of organelles and endomembranes (Figure 9). It is likely that many of these events are regulated by post-fibrillization post-translational dependent interactions between α-syn fibrils and other proteins and organelles. To the best of our knowledge, this is the first report of the formation of inclusions that recapitulate many of the cardinal features of LBs, including the presence of fibrillar α-syn, lipids, and membranous organelles.

### Quantitative proteomics reveals the temporal sequestration of membranous organelle, synaptic, and mitochondrial proteins into LB-like inclusions

Having demonstrated by CLEM imaging that the maturation of the newly formed fibrils (D14) into LB-like inclusions (D21) is accompanied by the active and gradual recruitment and sequestration of organelles and endomembranes over time (D14 to D21), we next sought to gain further insight into the molecular interactions and mechanisms that govern LB formation and maturation by mapping the proteome of α-syn inclusions over time. Toward this goal, we performed quantitative proteomic analysis on the insoluble fractions of PBS or PFF-treated neurons after 7, 14, or 21 days of treatment (Figure 4). The proteomic data generated from three independent experiments are presented as a volcano plot to depict the differentially expressed proteins based on the mean intensity differences versus t-test probability.

**Figure 4.**
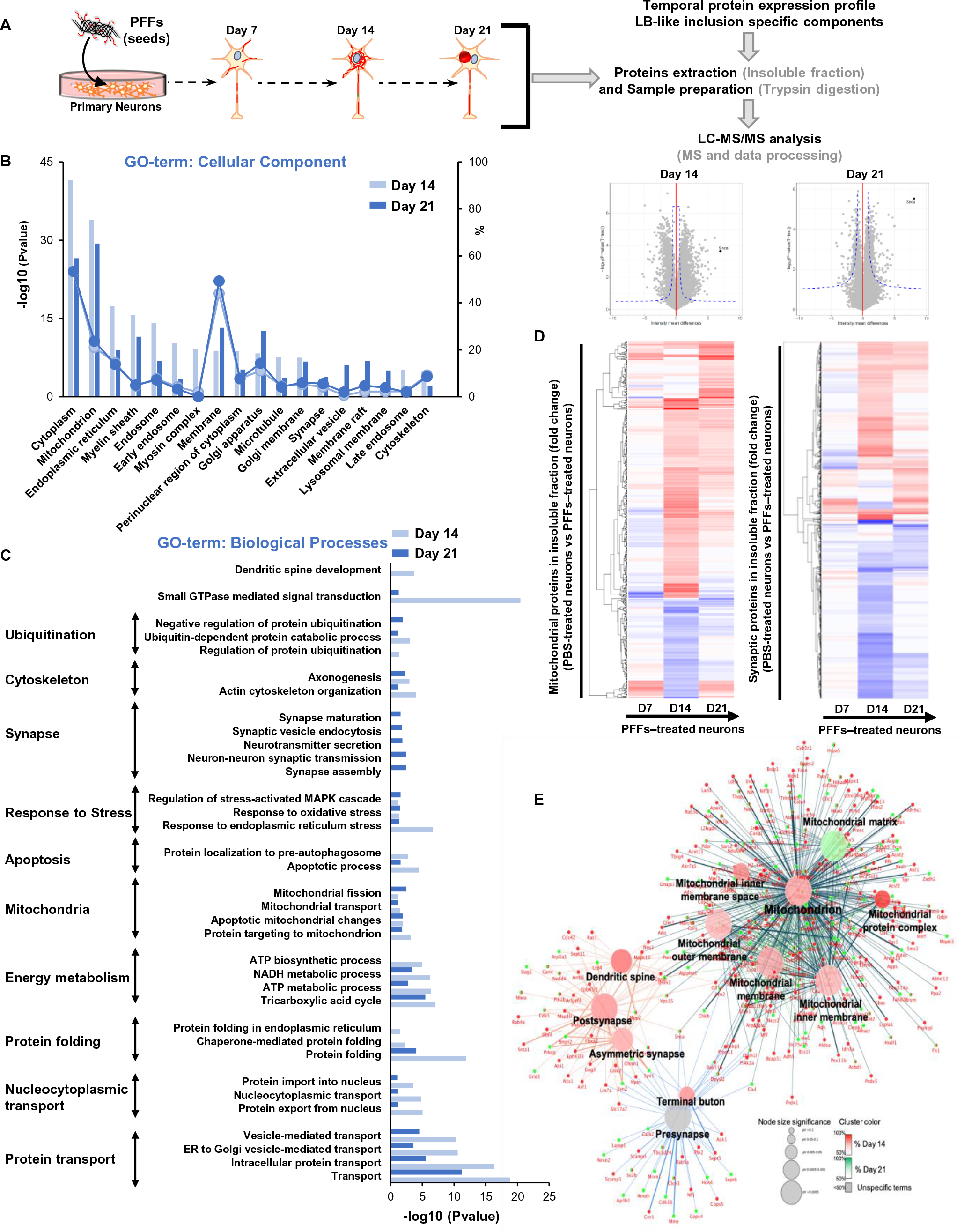
Temporal proteomic analyses of the protein contents found in the insoluble fraction of the PFF-treated neurons reveals a high increase in proteins related to the endomembrane system. **A.** Insoluble proteins from neurons treated with PBS and PFFs for 7, 14, and 21 days were extracted and analyzed using LC-MS/MS. Identified proteins were plotted using volcano plot. Mean difference (Log2 (Fold-Change) on the X-axis) between the insoluble fractions of PFF-treated neurons and PBS neurons treated for 14 (left panel) or 21 days (right panel) were plotted against Log_10_ (p-Value) on the Y-axis (T-Test). Dotted lines represent the FDR <0.05 (False Discovery Rate) and threshold of significance SO=1 assigned for the subsequent analysis. A detailed list of the hits shown in the volcano plot is available in Table 1.
**B-C.** Classification of the proteins significantly enriched in the insoluble fractions of the PFF-treated neurons at D14 and D21 by cellular component (**B**) and biological processes (**C**) using Gene Ontology (GO) enrichment analyses determined by DAVID analysis.
**D.** Heat map representing color-coded fold change levels of mitochondrial (right) and synaptic (left) proteins present in the insoluble fraction in neurons treated with PFFs 7, 14, and 21 days.
**E.** Cytoscape visualization of the proteins shared between 14 and 21 days PFFs treated insoluble fraction based on cellular compartment related to mitochondrial and synaptic function. The nodes (round shape) represent gene ontology terms; node color represents the group and node size represent the significance of each gene ontology term.

At D7, after multi comparisons correction using the Bonferroni method (FDR threshold <0.05, magnitude of proteins changes above 2 and a –log10 p-value >0.05), only α-syn and the Phospholipase C Beta 1 (Plcb1) were shown to be significantly enriched in the insoluble fraction with a nine- and two-fold increase in the insoluble fraction of the PFF-treated neurons, respectively (Figure S8A). No proteins from the endomembrane compartments were significantly enriched in the insoluble fraction of the PFF-treated neurons at D7. This finding is in line with our CLEM observations showing that at an early stage, α-syn seeded aggregates were mainly detected in the neurites as long filaments that did not appear to interact with intracellular organelles.

Next, we examined the protein contents of the α-syn inclusions at the intermediate stage, prior to (D14) and after (D21) the formation of LB-like inclusions (Figure 4). As shown in Figure 4A, the volcano plots show that 633 proteins and 568 proteins were significantly upregulated in the insoluble fraction of the PFF-treated neurons at D14 and D21, respectively. These proteins were then classified by cellular component (Figure 4B) and biological processes (Figure 4C) using the gene ontology (GO) term analysis and the database for annotation, visualization, and integrated discovery (DAVID) analysis and pathways (Figure S8B) using the Ingenuity Pathway Analysis (IPA). Classifications of the proteins by cellular compartment (Figure 4B) showed that the insoluble fractions of the PFFs-treated neurons at D14 and D21 were highly enriched by proteins that belong to the endomembrane compartments mainly from the mitochondria, the endoplasmic reticulum (ER), the Golgi and to a lesser extent the endolysosomal apparatus or synapses. These data are in line with the CLEM imaging showing that from D14 the newly formed fibrils start to interact with and encircled several organelles, including mitochondria, autophagosome, endosomes, lysosomes-like vesicles, and others endomembranous structures (Figures 2D) that are found later to be sequestered in the LB-like inclusions at D21 (Figures 2F and 3). Finally, several cytoskeletal proteins, including microtubule and myosin complexes, were also significantly enriched at D14 and D21 in the insoluble fraction of the PFF-treated neurons (Figure 4B). It is noteworthy that the close resemblance between the protein contents of the insoluble fraction of PFF-treated neurons at D14 and D21 is expected given that at D21 only ∼22% of the positive α-syn aggregates have completely transitioned to inclusions with an LB-like morphology (Figure S3E).

### Formation and maturation of LB-like inclusions are dynamic processes which involve proteins sequestration and alterations of key signaling pathways rather than simply α-syn fibrillization

Several studies have shown that protein aggregation and inclusion formation could contribute to neurodegeneration and disease progression through both gain of toxic function and loss of normal protein function, possibly through aberrant interactions involving protein aggregation or the sequestration of functional proteins within pathological inclusions^50–54^. Therefore, to gain insight into the mechanisms and pathways that are involved and affected by α-syn fibrillization and LB formation and maturation, the list of proteins found enriched in the PFF-treated neurons (Figure 4A) was further classified by biological processes (Figure 4C) and signaling pathways (Figure S8B). At the early stage of fibrils formation (D7), only α-syn and Plcb1 proteins were significantly enriched in the insoluble fraction of the PFF-treated neurons. At this stage, no biological processes were found to be altered (Figure S8A). Cellular compartment analysis showed that the insoluble fractions of PFF-treated neurons at D14 and D21 contained a large number of cytoskeletal proteins including several tubulin sub-units, microtubule-associated proteins, and motor proteins (kinesin, dynein, and myosin) (Figure 4B). This cluster of proteins was associated with the enrichment of several biological processes, including actin cytoskeleton organization, axonogenesis, and dendritic spine development (Figure 4C). The top canonical signaling pathways were also related to axonal guidance and microtubule regulation by the stathmin proteins (Figure S8B). The accumulation of the cytoskeletal proteins in the neuronal insoluble fraction is concomitant with the redistribution of newly formed α-syn aggregates from the neurites to the perinuclear region (Figures 1C-P, S3B-H). This suggests that the retrograde trafficking of newly formed α-syn fibrils is accompanied by the sequestration of the proteins involved in axonal transport. Impaired axonal transport has been shown to have dramatic effects on the intracellular trafficking of proteins, vesicles, and organelles^55, 56^. In line with this hypothesis, our temporal proteomic analysis revealed that, at D14 and D21, the most highly enriched terms for the biological processes were related to intracellular protein transport, including ER to Golgi mediated transport and the vesicle-mediated transport but also the nucleocytoplasmic protein transport (Figure 4C). Overall, the sequestration of proteins related to the intracellular transport inside α-syn inclusions seems to result in an impairment in the trafficking of organelles and vesicles, such as mitochondria, endosomes, lysosomes, and the synaptic vesicles^55, 56^, that eventually accumulate inside α-syn inclusions as evidenced by CLEM imaging (Figures 2 and 3). In line with this hypothesis, our proteomic data showed perturbation of the biological processes and signaling pathways related to mitochondria and synaptic compartments (Figures 4C and S8B). Interestingly, not only proteins involved in mitochondrial transport were enriched in the insoluble fraction of the PFF-treated neurons but also proteins related to mitochondrial dynamics, the mitochondrial apoptotic pathway, and the energy metabolism pathways (Tricarboxylic acid cycle, ATP and NADH metabolisms). This indicates that the process of inclusion formation that occurs throughout D14 to D21 might dramatically alter mitochondrial physiology, suggesting that mitochondrial dysfunction could be a major contributor to neurodegeneration in PD and synucleinopathies.

The upregulation of biological processes related to synaptic homeostasis (Figure 4C) was due to the accumulation of both pre and post-synaptic proteins (Figure 4E) mostly at D21 (Figure 4D). The processes involved in synaptic transmission, synapse assembly and synapse maturation (Figure 4C) were associated with an upregulation of the long-term synaptic depression and potentiation pathways but also of the endocannabinoid and CXCR4 signaling that regulate synapse function^57^ (Figure S8B). Several signaling pathways related to neurotoxicity were detected as upregulated in our analyses at D14, such as the Huntington’s disease signaling, mitochondrial dysfunction, Apoptosis signaling, the ER stress pathway, autophagy, and the neuroinflammation signaling pathway (Figure S8B).

Our proteomic and CLEM data provide strong evidence that formation of LB does not occur through simply the continued formation, growth, and assembly of α-syn fibrils but instead arises as a result of complex α-syn aggregation-dependent events that involve the active recruitment and sequestration of proteins and organelles over time. This process, which occurs mainly between 14-21 days, rather than simply fibril formation, which occurs as early as 7 days, seems to trigger a cascade of cellular processes, including changes in the physiological properties of the mitochondria and the synapses, that could ultimately lead to neuronal dysfunctions and toxicity.

### Transcriptomic analysis reveals multiple changes in gene expression during the formation of the newly formed fibrils and their maturation into LB-like inclusions

Next, to better understand the mechanisms of dysfunctions that ultimately lead to neurodegeneration, we investigated the transcriptomic changes in response to PFFs treatment over time. RNA sequencing (RNA-Seq) technology was used to explore the genome-wide transcript profiling in PBS and PFF-treated neurons after 7, 14, or 21 days of treatment. Data were generated from three independent experiments. RNA-Seq analysis (FDR threshold <0.01) identified a large number of genes that were differentially expressed between the PBS- and the PFF-treated neurons (Figure S9A). Interestingly, global transcriptome changes increased over time in the PFF-treated neurons. At D7, only 75 differentially expressed genes were identified, while at D14 the total number of genes increased to 435. At D21, 1017 genes were differentially expressed (Figure S9A), of which 455 were upregulated and 562 were downregulated (Figure S9A).

We first investigated the potential enrichment of cellular component (GO term analysis) for the differentially expressed genes in PFF-treated neurons in cellular compartment. At D7, the differentially expressed genes encoded for proteins located at the synapses, in the axons, or the secretory and exocytic vesicles (Figure 5A). These genes are thought to play regulatory roles in the neurogenesis processes, including the organization, the growth, and the extension of the axons and dendrites (Figure 5B). Thus, the presence of the newly formed fibrils that are mainly detected in the neuritic extensions at this stage could perturb the development and the differentiation of the neurites. At D14, among the differentially expressed genes, 329 genes (106 upregulated, 223 downregulated) were linked to the synaptic, neuritic, and vesicular cellular compartments (Figure 5A). Further analyses showed that these genes were associated with multiple biological processes, including neurogenesis, calcium homeostasis, synaptic homeostasis (organization, plasticity, and neurotransmission), cytoskeleton organization, response to stress, and neuronal cell death process (Figure 5B).

**Figure 5.**
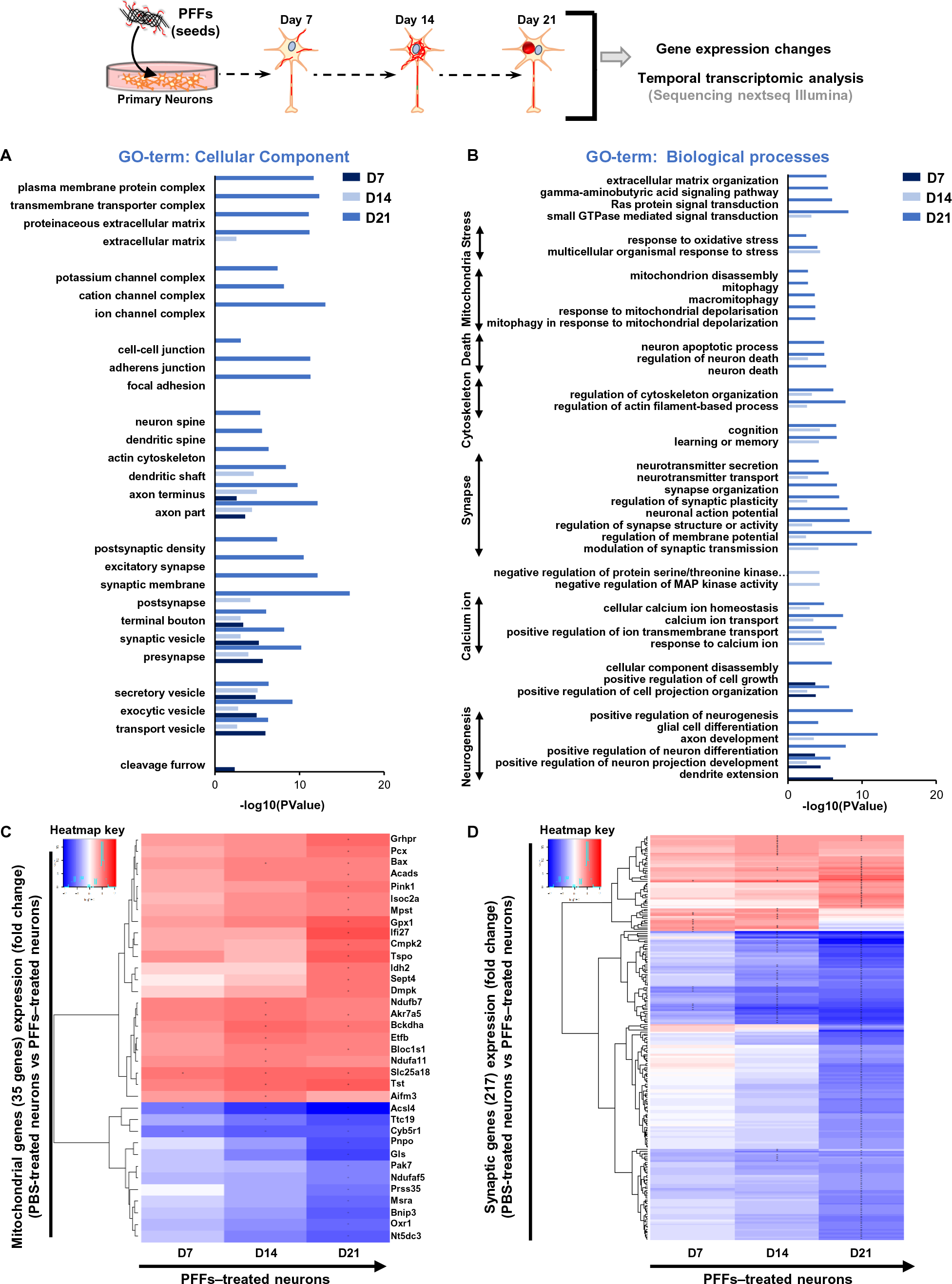
Gene expression level changes during the formation of the newly formed fibrils and their maturation into LB-like inclusions. Temporal transcriptomic analysis of the gene expression level in PBS-treated neurons vs. PFF-treated neurons treated for 7, 14, or 21 days. Genes with an absolute log2 fold-change greater than 1 and an FDR less than 0.01 were considered as significantly differentially expressed. Significant GO terms enrichment associated with cellular components (**A**) and biological processes (**B**) from differentially regulated genes were plotted against –log10 P value (t-test, PBS-treated neurons vs. PFF-treated neurons, Table 2). Log2 fold changes of the mitochondrial genes (**C and Table 3**), and the synaptic genes (**D and Table 4**) expression levels are represented over time. “+” and “-” indicate a significant upregulation or downregulation in the gene expression level respectively.

At D21, in addition to the axon and vesicles, our transcriptomic data showed an enrichment of genes that encode for proteins present in several cellular compartments, including the ion channels complex, plasma membrane protein complex, and the cell-cell junctions. More importantly, our data show that the expression level of 217 synaptic related genes was dramatically changed over time with a marked difference between D14 and D21 (Figures 5B, D, and S10B).

Most of the differentially expressed synaptic genes were significantly downregulated between D14 and D21 (Figures 5D and S9B). Strikingly, at D21, ∼20% of the genes differentially expressed in the PFF-treated neurons were related to synaptic functions, including neurotransmission processes and synapse organization. Also, at D21, biological processes related to the response to oxidative stress and mitochondria, including mitochondrion disassembly, mitophagy, and mitochondria depolarization, were significantly enriched (Figure 5B-C).

Finally, both transcriptomic and proteomic analyses revealed that shared biological processes (Figures 4C and 5B) are mostly involved in the cytoskeletal, synaptic, mitochondrial, and neuronal cell death functions. Altogether, our results show that alterations of the mitochondrial and synaptic functions, both at the proteomic and transcriptomic levels, occur primarily throughout D14 to D21, thus coinciding with inclusion formation, LB maturation, and cell death.

### Dynamics of Lewy body formation and maturation induce mitochondrial alterations

To validate our findings that mitochondrial dysfunctions are associated with the formation and the maturation of the LB-like inclusions, we assessed the mitochondrial activity over time in PFF-treated neurons. ICC for mitochondrial markers (VDAC1, TOM20, and TIM23) revealed strong co-localization of mitochondria with α-syn pS129-positive aggregates starting from D14 after PFF exposure. To assess whether this recruitment of mitochondrial components decreased mitochondrial function, we applied a combined protocol of high-resolution respirometry with Amplex Red based fluorometry to measure the production of mitochondrial reactive oxygen species (ROS).

Routine respiration of intact cells was significantly reduced at D21 (Figure 6B), while it was similar to PBS-treated control cells at the other assessed time points. Plasma membranes were subsequently permeabilized using digitonin, and substrates feeding into NADH-linked respiration were supplied. In the absence (N*_L_*) and presence (N*_P_*) of ADP, these respirational states did not significantly differ across all tested time points of PFF- and PBS-treatment.

**Figure 6.**
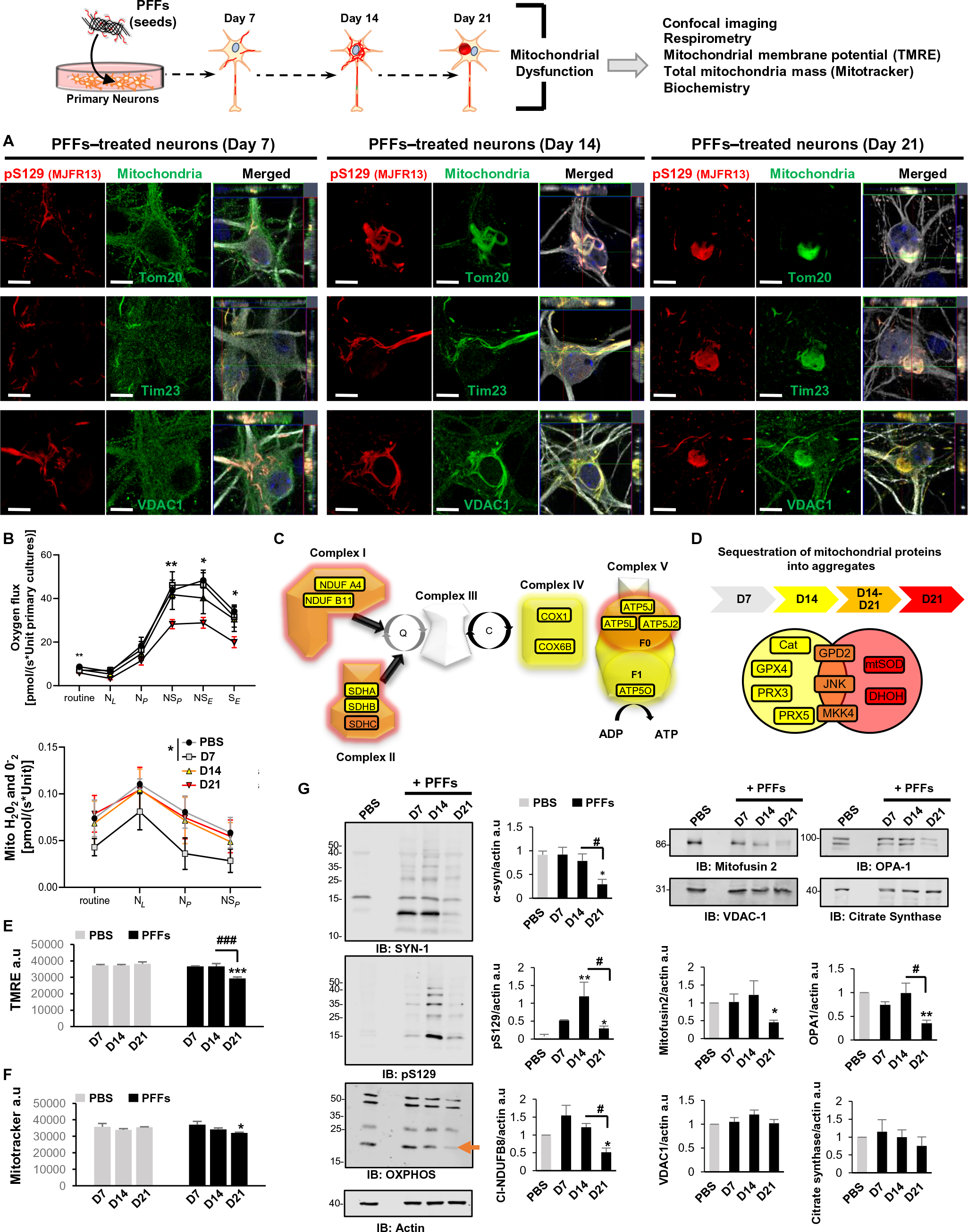
Dynamics of Lewy body formation and maturation induce mitochondrial alterations. Primary hippocampal neurons were treated with 70 nM of PFFs for 7, 14, or 21 days before assessment of mitochondrial parameters. **A.** ICC co-stainings of α-syn pS129 with VDAC1, TOM20, and TIM23 show increasing colocalization of mitochondrial components with pS129-positive aggregates over time. Scale bars = 5 μm. **B.** High-resolution respirometry revealed significant differences between PBS- and PFF-treated groups at D21 in several respirational states (**B**, upper panel). Concurrently assessed mitochondrial ROS-production was exclusively reduced at D7 (**B**, lower panel). Statistical results in **B** were obtained from 2-way repeated measurement ANOVAs based on a minimum of four independent experiments. **C-D**. Proteomic data on proteins implicated in mitochondrial dysfunction suggest proteins related to oxidative phosphorylation (**C**) and oxidative stress pathways (**D**) to be sequestered into aggregates only after D7 days. Mitochondrial membrane potential was assessed fluorometrically from cells loaded with TMRE (**E**); reduction in TMRE fluorescence intensity was observed at D21. Mitotracker green, labeling mitochondrial proteins independent of mitochondrial membrane potential, fluorescence (**F**) was moderately decreased at D21. **G**. WB analyses of total fractions derived from neuronal cultures. Components of the oxidative phosphorylation (OXPHOS) system (most notably the NDUF B8 subunit of complex I) are reduced at D21. A primary antibody mix for subunits of OXPHOS complexes I-V was used, the band indicated by an orange arrow represents CI-NDUF B8. Higher bands (from bottom to top) represent CII-SDHB, CIII-UQCRC2, and CV-ATP5A, respectively. Also, levels of the mitochondrial shape changing proteins mitofusin 2 and OPA1 were significantly decreased at D21. No significant changes in levels of citrate synthase or VDAC1 proteins were observed at any time. The graphs (**E-G**), represent the mean +/−SD of a minimum of three independent experiments. p<0.05 = *, p<0.005=**, p<0.0005=*** (ANOVA followed by Tukey HSD post-hoc test, PBS vs. PFF-treated neurons). p<0.05 = # p<0.05 = ### (ANOVA followed by Tukey HSD post-hoc test, D14 vs. D21 PFF-treated neurons).

Addition of succinate, drove overall respiration of all other groups to significantly higher levels than neurons at D21. This was the case for both maximum oxidative phosphorylation capacity (NS*P*) and maximum electron transport system capacity (NS*E*) in the uncoupled state (by CCCP). Blocking NADH-linked respiration by rotenone, yielding maximal succinate-driven respiration in the CCCP uncoupled state, maintained the significantly lower respiration of neurons at D21. Strikingly, the generation of mitochondrial ROS was significantly decreased at D7 but returned to baseline levels after longer periods of PFF treatment (Figure 6B, lower panel). In line with respiration results, analysis of proteomic data revealed significant recruitment of oxidative phosphorylation system components into aggregates starting from D14 (Figure 6C). Complexes I, II, and V were strongly affected up to D21, with complex II subunit SDHC being robustly detected in aggregates at D14 and D21 (Figure 6D). The strong sequestration of complex II components into aggregates might explain the stronger effects of α-syn pathology on succinate- than on NADH-linked substrate driven respiration.

The reduced mitochondrial ROS production at D7 (Figure 6B) correlated with the apparent lack of mitochondrial proteins involved in oxidative stress responses in aggregates (Figure 6D). Coinciding with the appearance of proteins and components of the JNK pathway in the aggregates, mitochondrial ROS production increased to baseline levels. Mitochondrial membrane potential was measured from attached cells by TMRE fluorometry (Figure 6E), and in line with respiration data, deterioration of mitochondrial membrane potential was apparent only at D21. At this time point, we also observed a slight reduction of (membrane-potential independent) binding of Mitotracker green to mitochondrial proteins (Figure 6F), indicating moderate loss of mitochondrial density.

To investigate the effects of the aggregation process on selected mitochondrial proteins in more detail, WB analyses were performed and the expression levels were assessed at distinct time points of this process (Figure 6G): outer mitochondrial membrane proteins (mitofusin 2 and VDAC1), matrix protein (Citrate synthase), inner mitochondrial membrane proteins, including OPA1 and subunits of oxidative phosphorylation complexes (OXPHOS), known to be labile when the respective complexes are not assembled. At D7, no altered mitochondrial protein levels were observed despite reduced mitochondrial ROS production and increases in α-syn pS129 and HMW species (Figure 6G). At D14, the levels of the investigated mitochondrial proteins were still essentially unaltered, while significant reductions were observed at D21 for pro-fusion protein OPA1 (long and short isoforms) and mitofusin 2 (Figure 6G). Similar effects were observed for subunits of oxidative phosphorylation complexes, in particular, complex I subunit NDUFB8 (Figure 6G). Importantly, levels of VDAC1 and citrate synthase, common markers of mitochondrial density, remained relatively unaffected, even in this period of massive respirational deficits and dropping mitochondrial membrane potential.

### Synaptic dysfunction is primarily linked to the formation and the maturation of the LB-like inclusions

Our transcriptomic and proteomic data strongly suggest that the reorganization of the newly formed fibrils into LB-like inclusions are concomitant not only with mitochondria defects but also with synaptic dysfunctions including neurotransmission, synaptic organization, and plasticity (Figures 4C and 5B). Therefore, we assessed the protein levels of pre- and post-synaptic markers [(respectively Synapsin I and Post-synaptic Density 95 (PSD95)]. We measured a dramatic reduction of these synaptic markers at D21 (Figure 7A).

ICC also revealed that the density of the synapses was reduced in neurons having an aggregates burden (Figure 7B-C). The loss of synapses correlates with the marked increase in neuronal cell death observed at D21.

Finally, both proteomic and transcriptomic data suggested dysregulation of the mitogen-activated protein kinase (MAPK) signaling pathways and the recruitment of the MAP kinases ERK1 (MAPK3) and ERK2 (MAPK1) within α-syn inclusions over time. These findings are especially relevant as the ERK signaling pathway is involved in regulating mitochondria fission and integrity^58–60^ and synaptic plasticity^61–64^, and is also an important player in PD pathogenesis^65–67^. WB analysis showed a significant reduction of the total protein level of the extracellular signal-regulated (ERK 1/2) proteins at D21. Interestingly most of the ERK proteins that were still expressed in neurons at D21 were highly phosphorylated, suggesting that the ERK pathway is still activated at the late stage of LB formation. Finally, ICC revealed that p-ERK 1/2 decorated the newly formed fibrils at D14 before being sequestered in the LB-like inclusions at D21 (Figure 7D). Also, our data showed that p-ERK 1/2 was only associated with the newly formed aggregates localized in the neuronal cell bodies, as evidenced by the absence of p-ERK 1/2 signal near the neuritic aggregates at D7 and D14. At D21, the granular p-ERK 1/2 immunoreactivity was mostly detected at the periphery of the inclusions as observed in the *bona fide* LBs found in human PD brain tissue^68, 69^.

### The formation and maturation of the LB-like inclusions are associated with neuronal cell death

We next assessed how the processes involved in the formation, remodeling, and interactome of α-syn aggregates could impact on the health of the cells at the distinct phases of α-syn fibrillization from the early stage of fibrils formation (D7), through the packing of the newly formed fibrils into aggregates (D7 to D14) and their maturation into LB-like inclusions (D14 to D21).

First, it is important to note that treatment of α-syn KO neurons with the PFFs (70 nM) did not induce neuronal death, even at D21, confirming that α-syn PFFs treatment and uptake is not toxic to the neurons (Figure S10). Cell death was only detectable in WT neurons treated with PFFs (Figure 8A-G). The first signs of toxicity in this neuronal population were only observed from D14, as indicated by the significant activation of caspase 3, a key marker of apoptosis.

**Figure 7.**
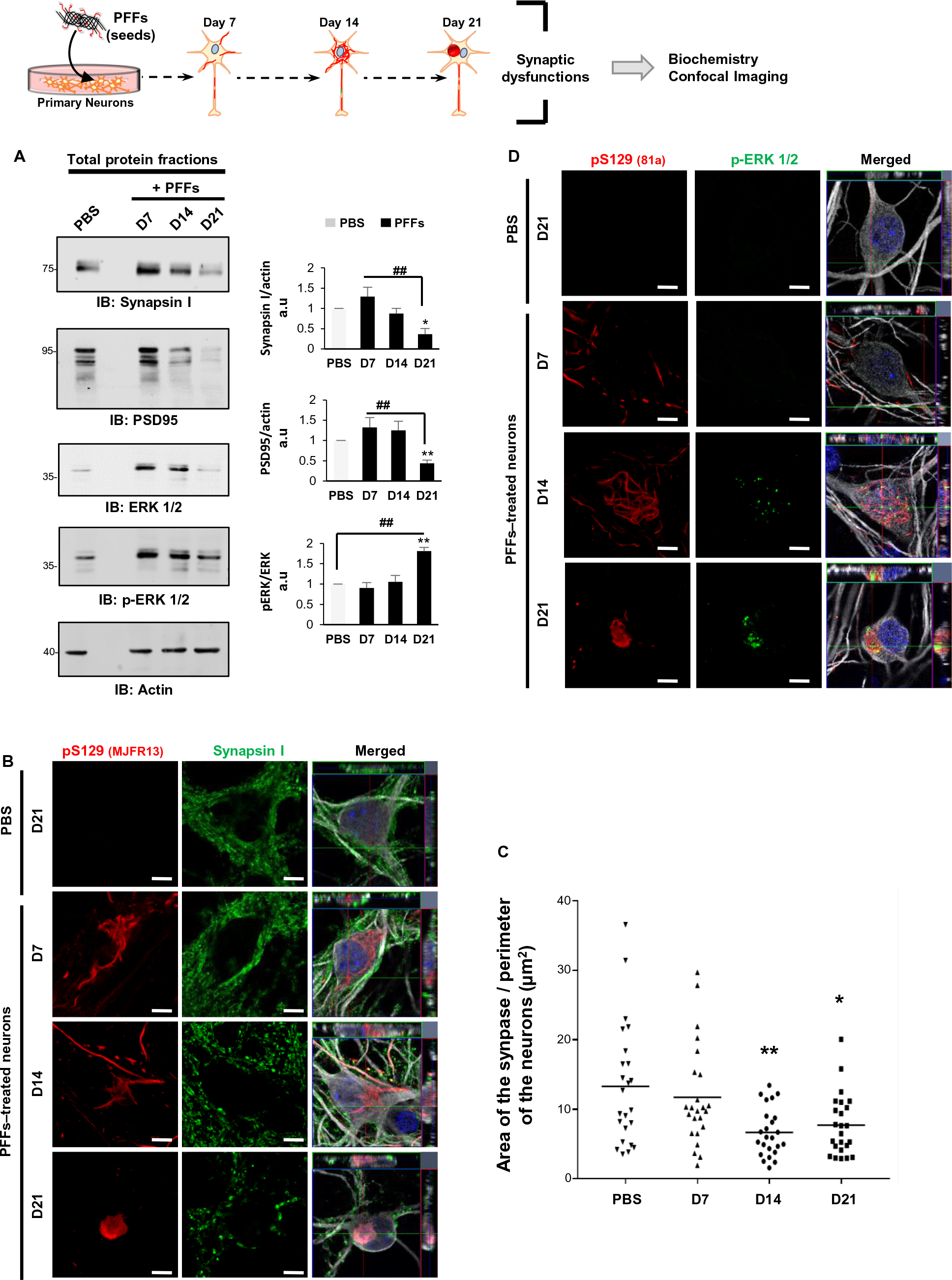
Synaptic dysfunctions were associated with the formation and maturation of the LB-like inclusions. Primary hippocampal neurons were treated with 70 nM of PFFs for 7, 14, or 21 days before assessment of synaptic parameters.
**A.** After total cell extraction, cells lysates were analyzed by WB. The levels of Synapsin I (pre-synaptic protein), PSD95 (post-synaptic protein), and ERK 1/2 (synaptic plasticity marker) were significantly decreased at D21. The phosphorylation level of ERK 1/2 (p-ERK 1/2) significantly increased over time. The graphs represent the mean ± SD of a minimum of three independent experiments. p<0.05 = *, p<0.005=** (ANOVA followed by Tukey HSD post-hoc test, PBS vs. PFF-treated neurons). p<0.05 = ### (ANOVA followed by Tukey HSD post-hoc test, D14 vs. D21 PFF-treated neurons).
**B-C.** Synaptic area decreases in PFF-treated neurons from D14. **B.** Aggregates were detected by ICC using pS129 (MJFR13) in combination with Synapsin I antibodies. Neurons were counterstained with microtubule-associated protein (MAP2) antibody, and the nucleus was counterstained with DAPI staining. Scale bars = 5 μm. **C.** Measurement of the synaptic area was performed over time. For each time-point, 24 neurons were imaged (z-step of 400 nm). The image analysis was performed in Fiji using a custom script to automatically analyze the z-stack images and measures the area of the synapse that was normalized to the perimeter of the neuron of interest. p<0.05 = *, p<0.005=** (ANOVA followed by Tukey HSD post-hoc test, PBS vs. PFF-treated neurons).
**D.** Aggregates were detected by ICC using pS129 (MJFR13) in combination with p-ERK 1/2 antibodies. ICC revealed the presence of p-ERK 1/2 associated with the newly formed fibrils at D14 before being sequestered in the LB-like inclusions at D21. Scale bars = 5 μm.

**Figure 8.**
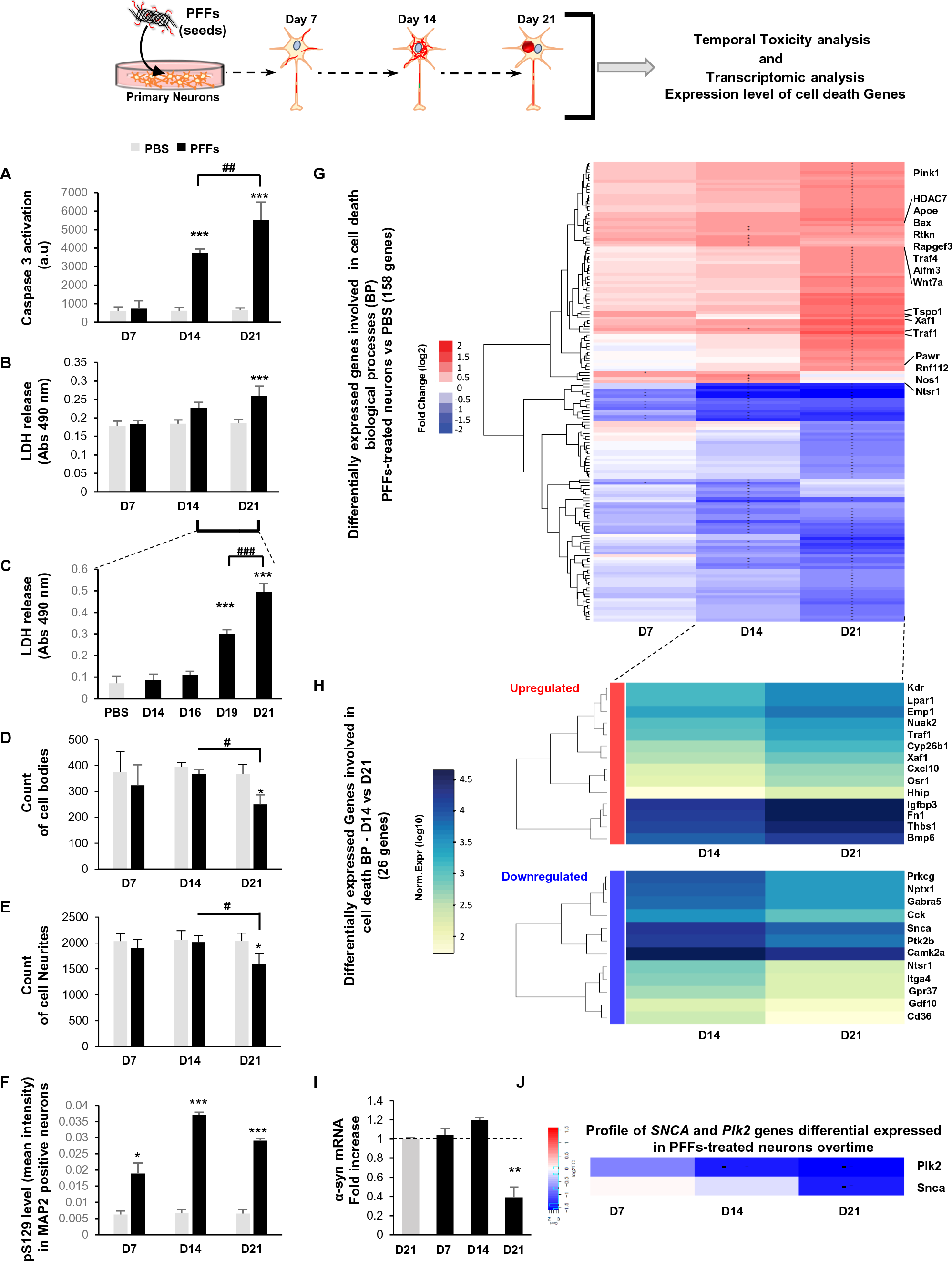
At late stages, the processes of formation and maturation of the LB-like inclusions are associated with neuronal toxicity. **A-F.** Addition of α-syn PFFs to primary neurons induces cell death in a time-dependent manner in WT neurons. **A-B.** Cell death levels were assessed in WT neurons treated with PFFs (70 nm) for up to D21 using complementary assays. Caspase 3 activation (**A**) was measured as an early event of apoptotic cell death, while late cell death events were assessed based on lactate dehydrogenase (LDH) release (**B-C**). For each independent experiment, triplicate wells were measured per condition. **D-F**. Neuronal loss was also assessed by high content image analysis (HCA) that allows us to determine the total number of neuronal cell bodies (**D**), neurites (**E**) and the total level of pS129 in MAP2 positive neurons (**F**) over time in PFF-treated primary culture. For each independent experiment, duplicated wells were acquired per condition, and nine fields of view were imaged for each well. Images were then analyzed using Cell Profile software to identify and quantify the number of neuronal cell bodies (DAPI and MAP2-positive areas) and the number of neurites (MAP2-positiveareas).
The graphs (**A-F**) represent the mean ± SD of three independent experiments. p<0.01=*, p<0.0001=*** (ANOVA followed by Tukey HSD post-hoc test, PBS vs. PFF-treated neurons). p<0.01= #, p<0.001= ## (ANOVA followed by Tukey HSD post-hoc test, PFF-treated neurons D14 vs. D21).
**G-H, J.** Temporal transcriptomic analysis of the gene expression level in PBS-treated neurons vs. PFF-treated neurons treated for 7, 14, or 21 days. Genes with an absolute log2 fold-change greater than 1 and an FDR less than 0.01 were considered as significantly differentially expressed. **G**. Log2 fold changes of the cell death genes at D7, D14, and D21 (**G**: PBS vs. PFF-treated neurons; **H**. Expression level of cell death genes D14 vs. D21. **J**. Fold changes of *SNCA* and *PLK2* genes (PBS vs. PFF-treated neurons). “+” and “-” indicate respectively a significant upregulation or downregulation in the gene expression level.
**I.** Quantitative RT-qPCR was performed with primers specific for α-syn at the indicated time-points after adding PFFs to WT neurons. Results are presented as fold change in comparison to the respective level in control neurons treated with PBS buffer. mRNA levels were normalized to the relative transcriptional levels of GAPDH and actin housekeeping genes. The graphs represent the mean ± SD of three independent experiments. p<0.005=** (ANOVA followed by Tukey HSD post-hoc test, PBS vs. PFF-treated neurons).

**Figure 9.**
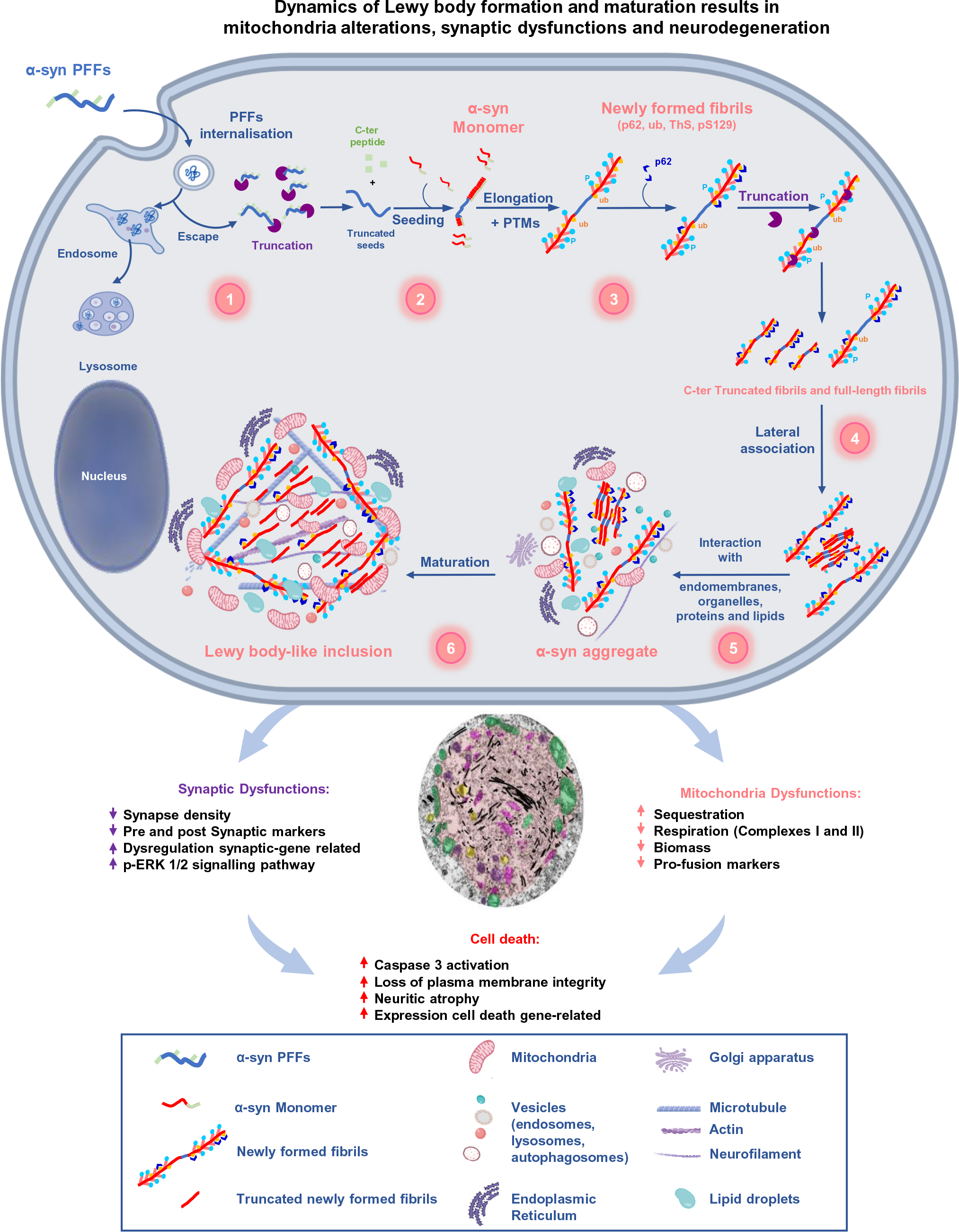
The dynamics of Lewy body formation, rather than simply α-syn fibrillization, is the primary cause of mitochondrial alterations, synaptic dysfunction, and neurodegeneration. Formation of α-syn inclusions in the context of the neuronal seeding requires a sequence of events starting with 1) the internalization and cleavage of PFF seeds (D0-D1); followed by 2) the initiation of the seeding by the recruitment of endogenous α-syn (D0-D4); 3) fibril elongation along with the incorporation of PTMs, such as phosphorylation at residue S129 (D4-D14); 4) formation of seeded filamentous aggregates (D7) that are fragmented and laterally associated over time (D7-D21); 5–6) the packing of the fibrils into LB-like inclusions (D14-D21) that have a morphology similar to the *bona fide* LB detected in human synucleinopathies (D21) is accompanied by the sequestration of organelles, endomembranes, proteins and lipids. Our integrated approach using advanced techniques in EM, proteomics, transcriptomic, and biochemistry clearly demonstrate that LB formation and maturation impaired the normal functions and biological processes in PFF-treated neurons causing mitochondrial alterations and synaptic dysfunctions that result in a progressive neurodegeneration.

From D14 to D21, the level of activated caspase 3 was further increased (Figure 8A-B). The loss of plasma membrane integrity was observed starting at D19, as reflected by the release of Lactate dehydrogenase (LDH) that significantly increased at D21 (Figure 8B-C). ICC combined with HCA confirmed the neuronal loss at D21, as evident by the significant decrease in the number of neuronal cell bodies and neurites (Figure 8D-E). In all the cell death assays, the presence of α-syn positive aggregates was confirmed by the measurement of pS129 level in MAP2-positive neurons (Figure 8F).

Strikingly, our transcriptomic analysis revealed that 156 genes involved in signaling pathways related to the regulation of neuronal cell death were also significantly changed over time in PFF-treated neurons (Figure 8G). These genes started to be significantly dysregulated at D14. At D21, 70 genes involved in cell death signaling pathways were upregulated in PFF-treated neurons compared to PBS-treated neurons (Figure 8G). Among these genes, 24 were directly related to apoptotic pathways, which is in line with our cell death assays showing significant activation of caspase 3 at D21. Between D14 and D21, 26 genes were differentially expressed in PFF-treated neurons (Figure 8H).

Our data also revealed a significant reduction of α-syn mRNA levels at D21 (Figure 8I), concomitant with the downregulation of the transcript level of the Polo-like kinase 2 (Plk2) (Figure 8J), the major kinase involved in the phosphorylation of α-syn at S129 residue. This suggests that several cellular responses are deployed by the neurons to prevent further aggregation and inclusion formation, either by downregulation of the protein level of α-syn or its phosphorylation state.

Overall, our data suggest a progressive neurodegeneration that coincides with the first signs of organelle accumulation and assembly of the LB-like inclusions.

Altogether our data demonstrate that the formation of fibrillar α-syn positive aggregates and their maturation into highly organized LB-like inclusions are accompanied by mitochondrial defects, synaptic dysfunctions, and the dysregulation of associated signaling pathways that ultimately lead to toxicity and neurodegeneration.

## Discussion

### The majority of existing cellular models are models of α-syn fibril formation rather than LBs

Several *in vitro* assays^70–73^ and cellular and animal models^12, 17, 22, 24, 71, 74^ of α-syn aggregation and pathology formation have been developed and are commonly used to study these processes. While many of these models reproduce specific aspects or stages of LB pathology formation, none has been shown to form pathological aggregates exhibiting the biochemical and organizational complexity of α-syn pathologies, including LBs, found in *post-mortem* brains of patients with synucleinopathies^8, 25, 26, 29–31, 39, 43, 74, 75^. For the most part, classification of α-syn aggregates as having LB-like features has been limited to very few biomarkers^12, 13, 17^, namely pS129, ubiquitin, and p62 immunoreactivity and the detection of high molecular weight aggregates in WB^13, 22, 40, 41^. In very few cases, detailed studies were performed to characterize the nature of the aggregates and their morphological properties. Failure of these models to reproduce the transition of α-syn from the aggregated/fibrillar stage to LB-like inclusions has hampered efforts to elucidate the molecular determinants and cellular pathways that regulate the different stages of LB pathology formation and to determine the role of this process in the pathogenesis of PD.

The use of PFFs to seed the aggregation of endogenous α-syn has enabled the reliable induction of α-syn fibril formation in neurons and other cell types^74, 76^. Based on the use of the LB markers discussed above, these seeding models are often presented as models of LB pathology. However, EM studies^22, 40, 41^ of the seeded aggregates, including our own (Figure 2A-F), clearly show the presence of mainly fibrillar aggregates at the time points usually used to assess seeding in this model (10–14 days) (Figures 1-2). These observations demonstrate that the α-syn modifications and interacting proteins commonly used to define LB pathology represent markers of α-syn fibril formation and, not necessarily, LB pathology, although these markers persist with α-syn fibrils during their transition to LBs and other types of α-syn pathologies.

### Extending the seeding process enabled the reconstitution of LB-like inclusions in neurons

While investigating the mechanism of seeding and pathology spreading in mice inoculated with PFFs, we observed a time-dependent shift in the morphology and localization of α-syn pathology from a filamentous neuritic fibrillar pathology to predominantly compact α-syn in cell body inclusions. Similarly, we observed that by simply extending the seeding process in primary neurons from 14 to 21 days, we began to see major changes in the localization and biochemical properties of the fibrils that eventually led to the formation rounded LB-like inclusions (Figures 2F, S5F and 3) that exhibit similar biochemical, structural, and architectural features as patient-derived LBs, as previously described by EM^7, 8, 30, 31, 46, 75, 77, 78^, IHC- or ICC^39, 42, 79^-based imaging, WB^34–38^, and proteomic^48, 49^ analyses. This includes the cleavage^33^ and accumulation of truncated fibrillar α-syn species^34–38^, the presence of many of the proteins found in LBs^42, 48, 49^ and the active recruitment of membranous organelles (e.g., mitochondria, the endolysosomal apparatus, and autophagolysosome-like vesicles)^8, 25, 26, 29–31, 39, 43, 74, 75, 78, 80, 81^. Although previous reports have established the presence of lipids in LBs from *post-mortem* PD tissues using different techniques, including Coherent Anti-Stokes Raman Scattering (CARS) microscopy^45, 46^ or synchrotron Fourier transform infrared micro-spectroscopy (FTIRM)^47^, very few studies have explored the different types of lipids and their distribution in LBs. Using different probes that target different types of lipids, we identified several classes of lipids that colocalize within the LB-like structures. Consistent with our confocal and CLEM imaging data, we observed enrichment of lipids (methyl ester staining) that are specific for mitochondria, ER and Golgi apparatus membranes within the LB-like inclusions. In addition, the phospholipids (cell membranes)^26^, sphingolipids (ER and Golgi complex)^26^, neutral lipids^25, 26^ (lipid droplets) and the cholesteryl esters^26^ (cholesterol transport) which have been previously reported as components of the *bona fide* LB inclusions were also detected reinforcing the similarities between the patient-derived LB pathologies and the LB-like inclusions formed in the seeding model. These findings underscore the critical importance of reassessing the role of lipids in the different stages of α-syn aggregation and LB formation and maturation.

Altogether our findings are in agreement with ultrastructure-based studies showing that LBs imaged in the *substantia nigra*^27, 29, 31, 46, 75, 78^, the hippocampal CA2 region^82^, or the Stellate ganglion^8^ from PD patients or non-PD patients^30^ exhibit organized structures that are enriched in deformed and damaged mitochondria, cytoskeletal components, and lipidic structures, including vesicles, fragmented membranes of organelles, and presynaptic structures. Recently, in *substantia nigra*, CLEM approach showed some LBs rich in lipids and membranous structures but devoid of α-syn fibrils^46^. However, a STED microscopy study of similar LBs conducted by the same group revealed the presence of a highly dense and organized shell of phosphorylated α-syn at the periphery of these LBs^45^. These observations raise the possibility that the CLEM procedure could have led to the disruption and removal of this peripheral aggregated pS129-α-syn. One alternative explanation that is consistent with our findings in the neuronal model is that the fibrils become increasingly fragmented over time (Figure 2, ∼50–350 nm in length at D21 in PFF-treated neurons). Such small fibrils, which are also C-terminally truncated^33^, might not be recognized and/or could be missed, especially when C-terminal antibodies were used to identify LBs (pS129 antibody or total α-syn antibody against amino acids 115–122)^46^. We cannot rule out the possibility that LB structures devoid of fibrillar α-syn aggregates exist and could represent a rare type of α-syn pathologies in PD and synucleinopathies. Indeed, several reports have confirmed the existence of a large morphological spectrum of LBs^3, 30, 39, 75, 77, 83–85^.

Altogether our results suggest that the neuronal seeding model recapitulates many of the key events and processes that govern α-syn seeding, aggregation, and LB formation (Figures S13 and 9). This model also allows disentangling of the two processes of fibril formation and Lewy body formation, thus paving the way for systematic investigation of the molecular and cellular determinants of each process and their contributions to neuronal dysfunction and degeneration.

### The proteome of the neuronal LB-like inclusions overlaps with that of PD and points to failure of the protein degradation machinery to clear α-syn inclusions

To further characterize the pathological relevance of the inclusions formed in the neuronal seeding model, we compared our proteomic data with results from previous studies on the proteome of brainstem or cortical LBs from PD human brain tissues. Approximately one-fourth of the proteins identified in our inclusion proteomic data at D14 and D21 were previously described as components of the LBs based on immunohistochemical-based studies ^1, 42, 86^ (Figure S11). When comparing our proteomic data of the LB-like inclusions with the proteome of human LBs, we found that ∼15–20% of the proteome overlaps with that of LBs from human PD brain tissues^48, 49^ (Figure S6). This is not surprising given the differences in the time of LB inclusion formation and maturation between primary neurons and in the human brain. It is also evident from our CLEM studies that the neuronal LB-like inclusions have not reached the same mature stage as the PD LBs. Interestingly, even previous studies that investigated the protein content of human cortical LBs showed ∼20% of similarities in the proteome of the inclusions. Overall, most of the proteins found in common between LBs from PD human brain tissues and the LB-like inclusions from the neuronal seeding model could be classified into three main categories: 1) proteins from the mitochondria and synaptic compartments or belonging to the endomembrane system; 2) cytoskeleton constituents and proteins associated with the intracellular trafficking of proteins and vesicles and the nucleocytoplasmic transport; and 3) proteins involved in the protein quality control (PQC) and the degradation/clearance machineries (Figures S11–S12). These results suggest that the sequestration of proteins related to the intracellular transport inside α-syn inclusions might result in the trafficking impairment of organelles and vesicles, such as mitochondria, endosomes, lysosomes, and the synaptic vesicles. Interestingly, dysfunction of the intracellular trafficking system and, in particular, defects in the ER-Golgi caused by α-syn have been shown to underlie the pathogenesis of PD at the early stage^56^. Previously published proteomic analyses using the same neuronal seeding model also indicated a potential role of the microtubule-dependent transport of vesicles and organelles^87^ during the process of aggregate formation (Figure S8C). Finally, the accumulation of large components of the proteins involved in the PQC and degradation/clearance machinery inside LB-like inclusions and the human *bona fide* LBs point toward a potential failure of the protein degradation machinery to clear off α-syn inclusions, leading to the intracellular accumulation of the aggregates as failed aggresomes-like inclusions. In line with this hypothesis, it has been shown that a conditional KO of 26S proteasome in mice led to the formation of LB-like inclusions consisting of filamentous α-syn, mitochondria and membrane-bound vesicles in neurons^88^. This is also consistent with our proteomic study showing a marked enrichment of the proteasomal system between D14 and D21 in the insoluble fractions PFFs-treated neurons (Figures 4 and S12). Altogether these findings highlight the potential role of the proteasomal system in driving the formation of the LB.

### The processes associated with LB formation and maturation, rather than simply α-syn fibril formation, are the major drivers of α-syn toxicity in neurons

Several studies have suggested that the formation of LBs represents a protective mechanism whereby aggregated and potentially toxic α-syn species are actively recruited into aggresome-like structures to prevent their aberrant interactions with other cytosolic proteins and their deleterious effects on cellular organelles^89^. A protective role for LBs is plausible if one assumes that this process is efficient. However, if this process stalls for any reason, then this will likely expose neurons to deleterious effects mediated by the presence of toxic proteins and damaged organelles and vesicles. To test this hypothesis, it is crucial to develop a neuronal and animal model system that enables uncoupling of the different stages of α-syn aggregation, fibrillization, and LB formation. One key advantage of neuronal models, as shown in this study, is that they permit detailed investigation of the molecular and cellular changes that occur during LB formation with high temporal resolution.

Our results show that formation of α-syn fibrils occurs as early as days 4–7, primarily in neurites, where they accumulate as long filamentous fibrils (Figures 1C–J and 2Aa). At the early stage of the seeding and fibrillization process (D7), the presence of these newly formed fibrils did not result in significant alteration of the proteome (Figures 4 and S8A), and induced only limited genetic and molecular perturbations (Figures 4 and 5) that did not impact neuronal viability up to D14 (Figures 8A-F and S13). These findings are in line with previous studies where no cell death was reported before D14^22, 40, 41, 90, 91^ and suggest that α-syn fibrillization is not sufficient to trigger neuron death, despite the large accumulation of α-syn fibrils throughout the cytosol of neurons.

However, the progressive accumulation of the filamentous fibrils in the neuronal cell bodies from D7 to D14 was accompanied by the shortening of the fibrils, their lateral association, and their interaction with surrounding organelles (Figure 2). These structural changes were concomitant with significant perturbations at the proteomic (Figure 4) and transcriptomic levels (Figure 5). Despite the interaction of the newly formed fibrils with a large number of proteins associated with cytoskeleton organization, mitochondrial functions, intracellular trafficking of proteins, vesicles, and nucleocytoplasmic transport, only the early stages of cell death were activated at D14 as evidenced by the significant activation of caspase 3 without yet the loss of plasma membrane integrity (Figures 5A-F and S13). The enrichment of proteins related to the chaperones machineries, the autophagy-lysosomal pathway, and the ubiquitin-proteasome system in the insoluble fraction of the PFF-treated neurons at D14 (Figures S12-S13) could suggest that the engagement of the protein quality control machinery and other related processes represent early cellular responses to prevent the formation and/or the accumulation of the fibrils in the cytosol. Similarly, transcriptomic analyses revealed that the formation of α-syn seeded aggregates induced (between D7 and D14) major dysregulations in the expression of genes involved in neurogenesis, calcium and synaptic homeostasis, cytoskeleton organization, response to stress, and neuronal cell death process (Figure 5). Therefore, the lack of noticeable toxic effects during the fibrillization process (D7–D14) might reflect the ability of neurons to activate multiple pathways to counter any negative effects induced by the fibrils.

In our extended neuronal seeding model, we observed that the major changes in the structural properties of the newly formed fibrils occur between days 14 and 21. This suggests that alternative cellular mechanisms take over to allow the structural reorganization of the fibrils and their sequestration within LB-like inclusions. This could represent an ideal detoxification mechanism by reducing aberrant interactions of the fibrils with cytosolic proteins or organelles. However, our CLEM (Figure 3) and proteomic analyses (Figure 4) showed that these structural changes were accompanied by significant recruitment and aberrant sequestration of intracellular proteins related to the intracellular transport inside α-syn inclusions. This seems to result in the trafficking impairment of organelles and vesicles such as mitochondria, endosomes, lysosomes, and the synaptic vesicles^56^ that eventually accumulate within LBs-like inclusions (Figures 2-4). This process was coupled with altered expression of genes associated with the cytoskeleton, mitochondria, and synaptic pathways together with an increase of the neuronal cell death pathways and neurodegeneration (Figures 5-8 and S13). Our data suggest that post-fibrillization post-translational modifications could play important roles in these processes by regulating the interactome of the fibrils and their interactions with cellular organelles. The sequestration of proteins and organelles eventually leads to a widespread loss of the biological functions, ultimately resulting in neurodegeneration. Finally, the reorganization of the newly formed fibrils into LB-like inclusions led to a significant decrease of α-syn both at the protein (Figure S7) and mRNA (Figure 8I) levels. While depletion of endogenous α-syn could prevent further aggregation and inclusion formation, it could also lead to loss of important physiological function(s) of α-syn with cellular dysfunctions as consequences.

### The dynamics of LB formation and maturation induce mitochondrial alterations

The prominent accumulation of mitochondria and mitochondrial components within the LB-like inclusions prompted us to further investigate how the various stages of α-syn fibrillization and LB formation influence mitochondrial functions (Figure S14). Although mitochondrial dysfunctions have been strongly implicated in PD, and other neurodegenerative diseases^92, 93^, the mechanisms by which mitochondrial dysfunction is induced and how it promotes pathology formation and/or neurodegeneration are still unclear.

Based on our CLEM results and literature evidence, we propose that interactions between the newly formed α-syn aggregates and mitochondria (D7 to D14) result in the increasing sequestration of mitochondrial organelles and proteins during the formation and the maturation of the LB-like inclusions leading to severe mitochondria defects observed at D21 (Figures 3-4, 6 and S14). This is in line with proteomic^48, 49^, IHC^42^ and EM^29, 31, 46, 75, 77^ characterizations of patient LBs, for which the presence of mitochondrial proteins and entire mitochondrial organelles have also been demonstrated.

Based on our results we suggest a working model on how subtle mitochondria-related events across the formation and maturation of LB-like inclusions might induce neurodegeneration (Figure S14).

A reduction in mitochondrial complex I, and of the pro-fusion proteins mitofusin 2 and OPA1 protein levels by WB (Figure 7) was only observed on D21, indicating an eventual failure of compensatory mechanisms during the formation of LB-like inclusions. These observations correlate with reduced mitochondrial membrane potential and a breakdown of mitochondrial respiration at this time point. Decreased respiration has already been shown in previous studies in the neuronal seeding model. However, respiration was assessed only between 8/9 and 14 days after PFFs treatment using mouse cortical neurons^94^ or rat midbrain neurons^95^, respectively. Importantly, in these previous studies high PFF concentrations (140nM or 350nM) were used, which in our hands induce cell death events faster. The concentration applied here (70nM) allowed investigation at a state, where LB-like inclusions occur. While mitochondrial respiration in these reports was assessed only in intact cells, here we measured maximum respiratory capacities at states requiring plasma membrane permeabilization. This allowed the additional determination of mitochondrial complexes I and complexes II to respiration capacity. Our data are in line with *post-mortem* studies^96–98^ reporting the deficiency of the respiratory chain in SN neurons from PD patients.

Given that we were able to detect different types of changes in mitochondrial functions and biochemical properties during the different stages associated with α-syn aggregation, fibrillization and formation of LB-like inclusions, we believe that the seeding model provides a powerful platform for dissecting the role of mitochondrial dysfunctions in synucleinopathies.

### Synaptic functions are impaired during the formation and the maturation of the LB

Several studies on *post-mortem* PD brain tissues and animal models have recently suggested that neuritic and synaptic degeneration correlate with early disease progression^99^. High presynaptic accumulation of insoluble α-syn has been proposed as a greater contributor to the development of clinical symptoms than LBs. Our proteomic (Figure 4) and transcriptomic analyses (Figure 6) suggest that mitochondrial defects are accompanied by an alteration of synapse-related RNA and protein levels. However, it remains unclear which pathways in the mitochondria-synapse signaling loop is dysregulated first by the formation and the maturation of α-syn pathological inclusions^100^. Consistent with *postmortem* studies of PD patients^101, 102^, during the transition from fibrils to LB-like inclusion, we showed a reduction of synaptic density, alterations of the protein level of both pre- and post-synaptic markers (Figure 8), and dysregulation of the synaptic transcriptome (Figure 6). Unlike mitochondrial dysfunction, which are manifested mainly once newly formed α-syn fibrils are reorganized into LB-like inclusions (D14-D21), early synaptic changes at the transcriptomic level were observed concomitant with the beginning of the α-syn fibrilization process and the appearance of α-syn-seeded aggregates inside the neuritic extensions at the early stage of the seeding model (D4–D7) (Figures 5E and S14). This suggests that synaptic dysfunction or different pathways linked to synaptic dysfunction are affected at the different stages of α-syn aggregation and LB formation and occur before the early onset of neurodegeneration, supporting the hypothesis that synaptopathy is an initial event in the pathogenesis of PD and related synucleinopathies^102, 103^. Synaptotoxicity reflected by the downregulation of the synaptic activity and the loss of synapses was also recently reported in seeding neuronal models^22, 91^.

Finally, the ERK 1/2 signaling was also shown to be upregulated during both the formation and the maturation of the LB-like inclusions (Figure 4). The ERK 1/2 pathway regulates neuronal cell death, oxidative stress, mitochondria fission, and integrity, as well as synaptic plasticity^65^, which are the major biological processes dysregulated in PD. Similar to what has been previously reported in *bona fide* LB from PD patients^68, 69^, we found that p-ERK 1/2 was recruited inside the LB-like inclusions (Figure 7D). Therefore, dysregulation of this central signaling pathway during the formation and the maturation of the LB-like inclusions could affect mitochondrial and synaptic functions, potentially explaining our observation in the neuronal seeding model and PD brains.

In conclusion, our work provides a comprehensive characterization of a highly reproducible neuronal model that recapitulates many of the key biochemical, structural, and organizational features of LB pathologies in PD brains. Using integrative imaging approaches, we were able to dissect the key processes involved the formation of LB-like pathologies and provide novel insight into their contribution to neuronal dysfunction and neurodegeneration. To our knowledge, this is the first report that shows the formation of such LB-like inclusions and presents a comprehensive characterization of the different stages of LB formation, from seeding to fibrillization to the formation of LB-like inclusions at the molecular, biochemical, proteomic, transcriptomic, and structural levels. These advances, combined with the possibility to uncouple the three key processes (seeding, fibrillization, and LB formation) and investigate them separately, provide unique opportunities to 1) investigate the molecular mechanisms underpinning these processes; 2) elucidate their role in α-syn-induced toxicity and potential contributions to the pathogenesis of PD; and 3) to screen for therapeutic agents based on the modulation of these pathways. The use of a model system that allows the investigation of these processes over a longer period of time (e.g., iPSC derived neurons) could enable further insights into the molecular mechanisms that regulate LB pathology formation, maturation, and pathological diversity in PD and synucleinopathies^3^, especially if the development of such model systems is driven by insight from human pathology.

## Material and Methods

### Antibodies and compounds

Information and RRID of the primary antibodies used in this study are listed in Figure S2. Tables include their clone name, catalog numbers, vendors, and respective epitopes.

### Expression and Purification of Mouse α-syn

pT7-7 plasmids were used for the expression of recombinant mouse α-syn in *E. coli*. Mouse wild type (WT) α-syn was expressed and purified using anion exchange chromatography (AEC) followed by reverse-phase High-Performance Liquid Chromatography (HPLC) and fully characterized as previously described^104, 105^.

### Primary culture of hippocampal neurons and treatment with mouse α-syn fibrils

Primary hippocampal neurons were prepared from P0 pups of WT mice (C57BL/6JRccHsd, Harlan) or α-syn KO mice (C57BL/6J OlaHsd, Harlan) and cultured as previously described^106^. The neurons were seeded in black, clear-bottomed, 96-well plates (Falcon, Switzerland), 6-wells plates, or onto coverslips (CS) (VWR, Switzerland) previously coated with poly-L-lysine 0.1% w/v in water (Brunschwig, Switzerland) at a density of 300,000 cells/mL. After 5 days in culture, the WT or α-syn KO neurons were treated with extracellular mouse α-syn fibrils to a final concentration of 70 nM and cultured for up to 21 days as previously described^22, 23^. All procedures were approved by the Swiss Federal Veterinary Office (authorization number VD 3392).

### Immunocytochemistry (ICC)

After α-syn PFFs treatment, HeLa cells or primary hippocampal neurons were washed twice with PBS, fixed in 4% PFA for 20 min at RT, and then immunostained as previously described^105^. The antibodies used are indicated in the corresponding legend section of each figure. The source and dilution of each antibody can be found in Figure S2. The cells plated on CS were then examined with a confocal laser-scanning microscope (LSM 700, Carl Zeiss Microscopy, Germany) with a 40× objective and analyzed using Zen software (RRID:SCR_013672). The cells plated in black, clear-bottom, 96-well plates were imaged using the IN Cell Analyzer 2200 (with a ×10 objective). For each independent experiment, two wells were acquired per tested condition, and in each well, nine fields of view were imaged. Each experiment was reproduced at least three times independently.

### Quantitative high-content wide-field cell imaging analyses (HCA)

After α-syn PFFs treatment, primary hippocampal neurons plated in black, clear-bottom, 96-well plates (BD, Switzerland) were washed twice with PBS, fixed in 4% PFA for 20 min at RT and then immunostained as described above. Images were acquired using the Nikon 10×/ 0.45, Plan Apo, CFI/60 of the IN Cell Analyzer 2200 (GE Healthcare, Switzerland), a high-throughput imaging system equipped with a high-resolution 16-bits sCMOS camera (2048×2048 pixels), using a binning of 2×2. For each independent experiment, duplicated wells were acquired per condition, and nine fields of view were imaged for each well. Each experiment was reproduced at least three times independently. Images were then analyzed using Cell profiler 3.0.0 software (RRID:SCR_007358) for identifying and quantifying the level of LB-like inclusions (stained with pS129 antibody) formed in neurons MAP2-positive cells, the number of neuronal cell bodies (co-stained with MAP2 staining and DAPI), or the number of neurites (stained with MAP2 staining).

### Correlative Light and Electron Microscopy (CLEM)

Primary hippocampal neurons were grown on 35 mm dishes with alphanumeric searching grids etched to the bottom glass (MatTek Corporation, Ashland, MA, USA) and treated with WT PFFs. At the indicated time point, cells were fixed for 2 h with 1% glutaraldehyde and 2.0% PFA in 0.1 M phosphate buffer (PB) at pH 7.4. After washing with PBS, ICC was performed (for more details, see the corresponding section in the Materials and Methods). Neurons with LB-like inclusions (positively stained for pS129) were selected by fluorescence confocal microscopy (LSM700, Carl Zeiss Microscopy) for ultrastructural analysis. The precise position of the selected neuron was recorded using the alpha-numeric grid etched on the dish bottom. The cells were then fixed further with 2.5% glutaraldehyde and 2.0% paraformaldehyde in 0.1 M PB at pH 7.4 for another 2 h. After washing (5 times for 5 min) with 0.1 M cacodylate buffer at pH 7.4, cells were post-fixed with 1% osmium tetroxide in the same buffer for 1 h and washed with double-distilled water before contrasting with 1% uranyl acetate water for 1 h. The cells were then dehydrated in increasing concentrations of alcohol (twice at 50%, once at 70%, once at 90%, once at 95%, and twice at 100%) for 3 min each wash. Dehydrated cells were infiltrated with Durcupan resin diluted with absolute ethanol at 1:2 for 30 min, at 1:1 for 30 min, and 2:1 for 30 min, and twice with pure Durcupan (Electron Microscopy Sciences, Hatfield, PA, USA) for 30 min each. After 2 h of incubation in fresh Durcupan resin, the dishes were transferred into a 65°C oven for the resin to polymerize overnight. Once the resin had hardened, the glass CS on the bottom of the dish was removed by repeated immersion in hot (60°C) water, followed by liquid nitrogen. The cell of interest was then located using the alphanumeric coordinates previously recorded and a razor blade used to cut this region away from the rest of the resin. This piece was then glued to a resin block with acrylic glue, trimmed with a glass knife using an ultramicrotome (Leica Ultracut UCT, Leica Microsystems), and then ultrathin sections (50–60 nm) were cut serially from the face with a diamond knife (Diatome, Biel, Switzerland) and collected onto 2 mm single-slot copper grids coated with formvar plastic support film. Sections were contrasted with uranyl acetate and lead citrate and imaged with a transmission electron microscope (Tecnai Spirit EM, FEI, The Netherlands) operating at 80 kV acceleration voltage and equipped with a digital camera (FEI Eagle, FEI).

### Immunogold Staining

Primary hippocampal neurons were grown on 35 mm dishes with alphanumeric searching grids etched to the bottom glass (MatTek Corporation, Ashland, MA, USA) and treated with WT PFFs. At the indicated time point, cultured cells were fixed for 2 h with a buffered mix of paraformaldehyde (2%) and glutaraldehyde (0.2%) in PB (0.1M), then washed in PBS (0.01M). After washing with PBS, ICC was performed as described^105^. Briefly, cells were incubated with pS129 antibody (81a, Figure S4B) for 2 h at RT diluted in the blocking solution [(3% BSA and 0.1% Triton X-100 in PBS), (PBS-BSA-T)]. After five washes with PBS-BSA-T, neurons were incubated with the Alexa Fluor^647^ Fluoronanogold secondary antibody (Nanoprobes, USA) for 1 h at RT. Neurons were washed five times with PBS-BSA-T and mounted in polyvinyl alcohol mounting medium supplemented with anti-fading DABCO reagent. Neurons with LB-like inclusions (positively stained for pS129) were selected by fluorescence confocal microscopy (LSM700, Carl Zeiss Microscopy) for ultrastructural analysis. The precise position of the selected neuron was recorded using the alpha-numeric grid etched on the dish bottom. The cells were next washed in phosphate buffer (0.1M) and fixed again in 2.5% glutaraldehyde before being rinsed in distilled water. A few drops of a silver enhancement solution (Aurion, Netherlands) were then added to each CS to cover all of the cells and left in the dark for 1 h at RT. The solution was then removed, and the CS washed in distilled water followed by 0.5% osmium tetroxide for 30 min and 1% uranyl acetate for 40 min. Following this, the CS were dehydrated through increasing concentrations of ethanol and then transferred to 100% epoxy resin (Durcupan, Sigma Aldrich) for 4 h and then placed upside down on a glass CS in a 60°C oven overnight. Once the resin had cured, the CS was removed, and regions of interest were cut away with a razor blade and mounted onto a blank resin block for thin sectioning. Serial thin sections were cut at 50 nm thickness with a diamond knife (Diatome, Switzerland) in an ultramicrotome (UC7, Leica Microsystems) and collected onto a pioloform support firm on copper slot grids. These were further stained with lead citrate and uranyl acetate. Sections were imaged with a digital camera (Eagle, FEI Company) inside a transmission electron microscope (Tecnai Spirit, FEI Company) operating at 80 kV.

### Cell lysis and WB analyses of primary hippocampal neurons

#### Preparation of soluble and insoluble fractions

After α-syn PFFs treatment, primary hippocampal neurons were lysed as described in Volpicelli-Daley *et al.*^22, 23^. Briefly, treated neurons were lysed in 1% Triton X-100/ Tris-buffered saline (TBS) (50 mM Tris, 150 mM NaCl, pH 7.5) supplemented with protease inhibitor cocktail, 1 mM phenylmethane sulfonyl fluoride (PMSF), and phosphatase inhibitor cocktail 2 and 3 (Sigma-Aldrich, Switzerland). After sonication using a fine probe [(0.5-s pulse at an amplitude of 20%, ten times (Sonic Vibra Cell, Blanc Labo, Switzerland], cell lysates were incubated on ice for 30 min and centrifuged at 100,000 g for 30 min at 4°C. The supernatant (soluble fraction) was collected while the pellet was washed in 1% Triton X-100/TBS, sonicated as described above, and centrifuged for another 30 min at 100,000 g. The supernatant was discarded whereas the pellet (insoluble fraction) was resuspended in 2% sodium dodecyl sulfate (SDS)/TBS supplemented with protease inhibitor cocktail, 1 mM PMSF, and phosphatase inhibitor cocktail 2 and 3 (Sigma-Aldrich, Switzerland), and sonicated using a fine probe (0.5-s pulse at amplitude of 20%, 15 times).

#### Total cell lysates

After α-syn PFFs treatment, primary hippocampal neurons were lysed in 2% sodium dodecyl sulfate (SDS)/TBS supplemented with protease inhibitor cocktail, 1 mM PMSF, and phosphatase inhibitor cocktail 2 and 3 (Sigma-Aldrich, Switzerland) and boiled for 10 min. The bicinchoninic acid (BCA) protein assay was performed to quantify the protein concentration in the total, soluble, and insoluble fractions before addition of Laemmli buffer 4× (4% SDS, 40% glycerol, 0.05% bromophenol blue, 0.252 M Tris-HCl pH 6.8, and 5% β-mercaptoethanol). Proteins from the total, soluble, and the insoluble fractions were then separated on a 16% tricine gel, transferred onto a nitrocellulose membrane (Fisher Scientific, Switzerland) with a semi-dry system (Bio-Rad, Switzerland), and immunostained as previously described^105^.

### Proteomic identification of the proteins enriched in the insoluble fraction of the PFF-treated neurons

Primary hippocampal neurons were treated with PBS or PFFs for 7, 14, or 21 days. Cells were lysed at the indicated time-points into the soluble and insoluble fractions as described above. Proteins from the insoluble fraction were then separated by SDS-PAGE on a 16% polyacrylamide gel, which was then stained with Coomassie Safestain (Life Technologies, Switzerland). Each gel lane was entirely sliced, and proteins were in-gel digested as previously described^107^. Peptides were desalted on stageTips^108^ and dried under a vacuum concentrator. For LC-MS/MS analysis, resuspended peptides were separated by reversed phase chromatography on a Dionex Ultimate 3000 RSLC nano UPLC system connected in-line with an Orbitrap Lumos (Thermo Fisher Scientific, Waltham, USA). Label-free quantification (LFQ) was performed using MaxQuant 1.6.0.1^109^ against the UniProt mouse database (UniProt release 2017_05). Perseus was used to highlight differentially quantified proteins^110^. Reverse proteins, contaminants, and proteins only identified by sites were filtered out. Biological replicates were grouped together. Protein groups containing a minimum of two LFQ values in at least one group were conserved. Empty values were imputed with random numbers from a normal distribution (Width: 0.5 and Downshift: 1.7). Significant hits were determined by a volcano plot-based strategy, combining t-test p-values with ratio information^111^. Significance curves in the volcano plot corresponding to a SO value of 0.4 (D7 and D21) or 0.5 (D14) and 0.05 FDR were determined by a permutation-based method. Further graphical displays were generated using a homemade program written in R (https://www.R-project.org/). Table 1 contains the final results.

### Quantification of cell death in primary hippocampal neurons

Primary hippocampal neurons were plated in a 96-well plate and treated with α-syn PFFs (70 nM or up to 500 nM). After 7, 14, and 21 days of treatment, cell death was quantified using complementary cell death assays.

#### Quantification of active caspase 3

The CaspaTag fluorescein caspase 3 activity kit (ImmunoChemistry Technologies, MN, USA) allows the detection of active effector caspase (caspase 3) in living cells. These kits use the specific fluorochrome peptide inhibitor (FAM-DEVD-fmk) of the caspase 3 (FLICA). This probe passively enters the cells and binds irreversibly to the active caspases. Neurons were washed twice with PBS and incubated for 30 min at 37°C with FAM-DEVD-FMK in accordance with the supplier’s instructions. Fluorescein emission was quantified using a Tecan infinite M200 Pro plate reader (Tecan, Maennedorf, Switzerland) with excitation and emission wavelengths of 487 nm and 519 nm respectively.

#### Quantification of LDH release

Using a CytoTox 96® Non-Radioactive Cytotoxicity Assay (Promega, Switzerland), the lactic acid dehydrogenase (LDH) released into culture supernatants was measured following the manufacturer’s instructions. After a 30 min coupled enzymatic reaction, which results in the conversion of a tetrazolium salt (INT) into a red formazan product, the amount of color formed, that is proportional to the number of damaged cells, was measured using a Tecan infinite M200 Pro plate reader (Tecan, Maennedorf, Switzerland) at a wavelength of 490 nm.

### Quantitative real-time RT-PCR and transcriptomic analysis

Primary hippocampal neurons were treated with WT PFFs or PBS buffer for 7, 14, and 21 days. Total RNA was isolated using the RNeasy Mini Kit (Qiagen, Switzerland) according to the manufacturer’s protocol. The concentration of each sample was measured using a NanoDrop (NanoDrop Technologies, Wilmington, DE, USA), and the purity was confirmed using the ratios at (260/280) nm and (260/230) nm.

#### Quantitative real-time RT-PCR

RNA (2 µg) was used to synthesize cDNA using the High-Capacity RNA-to-cDNA Kit (Life Technologies) following the manufacturer’s instructions. To quantify α-syn mRNAs levels, we used the SYBR Green PCR master mix (Pack Power SYBR Green PCR mix, Life Technologies). Q-PCR was performed using the following primers synthesized by Microsynth (Balgach, Switzerland):

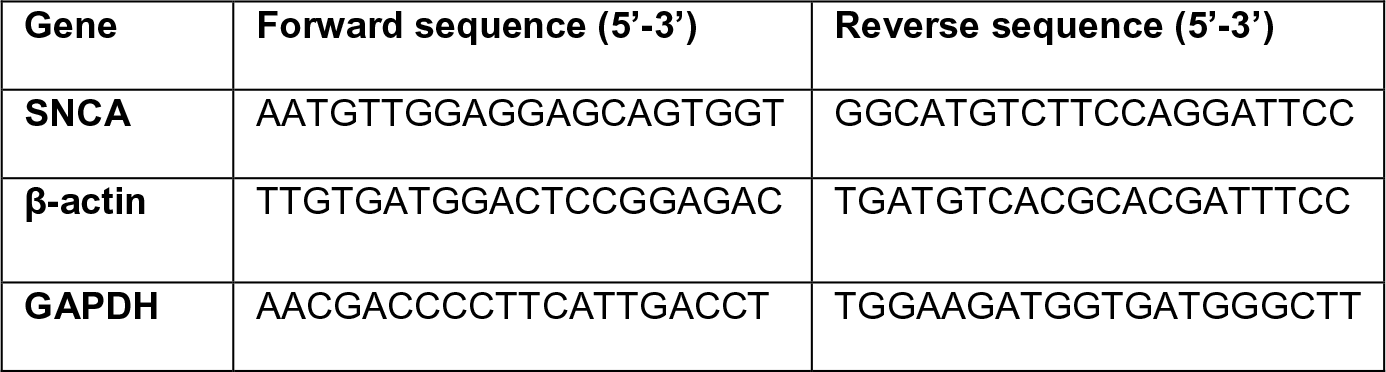

Forty cycles of amplification were then performed in an ABI Prism 7900 (Applied Biosystem, Foster City, USA) using a 384-well plate, which allowed the simultaneous analysis of the genes of interest and the housekeeping genes (β-actin and GAPDH) that serve as references for the normalization step. For each independent experiment, triplicated wells were acquired per condition, and each experiment was reproduced at least three times independently. To quantify the expression level of the genes of interest in the different conditions tested, the comparative 2^−ΔΔ*CT*^ method was used where ΔΔCT = ΔCT(target gene)−ΔCT(reference gene) and ΔCT = CT(target gene)−CT(reference gene). Results were expressed as the fold change relative to control neurons (2^−ΔΔ*CT*^). The geNorm method (RRID: SCR_006763, https://genorm.cmgg.be/) was performed to assess the most stable reference gene that should be used for the normalization of the gene expression^112^.

#### Temporal transcriptomic analysis

Libraries for mRNA-seq were prepared according to manufacturer’s instructions with the TruSeq stranded mRNA kit (Illumina) starting from 300 ng of good-quality total RNAs (RNA quality scores >8.9 on the TapeStation 4200). Libraries were subsequently loaded at 1.44 pM on two High Output flow cells (Illumina) and sequenced in a NextSeq 500 instrument (Illumina) according to manufacturer instructions, yielding 75 nt paired-end reads.

RNA-seq reads were aligned to the mm10 mouse reference genome (GRCm38.p5) with <u>STAR</u> (version 2.5)^113^. An average of 52 million of uniquely mapped reads per sample was then assigned to genes (<u>gencode primary assembly v16</u>) using <u>HTSeq-count</u> (version 0.9)^114^. The analysis was focused on protein-coding and antisense RNA genes only, representing a total of 23,057 studied genes. Lowly expressed genes (CPM < 5) and genes present in less than three samples were excluded prior to further processing. Normalization and differential expression analysis of the remaining 11,767 genes were done with the R package DEseq2 (version 1.2)^115^, using the internal DEseq and the results methods. Genes with an absolute log2 fold-change greater than 1 and an FDR less than 0.01 were considered as significantly differentially expressed. Table 2 [combined_DEseq2_FC1_FDR_0_01_PFatLeastOnce.xls] contains the final results. Tables 3-4 contains respectively analyses of mitochondrial and synaptic genes significantly enriched.

### Respirometry on detached primary neurons

Primary hippocampal neurons were plated 4 weeks before respiration experiments. At 21, 14, or 7 days before respiration, neurons were exposed to 70 nM PFFs. Immediately before respiration experiments, cells were carefully washed with PBS, and exposed to 0.05% trypsin for 35–40 min, until detachment of cells was clearly visible, after which DMEM F12 medium was used to neutralize the trypsin. Cells were gently spun down (300 g for 3 min), and the cells of 1.5 wells of 6 well plates were resuspended in DMEM F12 and used for high-resolution respirometry. The cell suspension was transferred to MiR05 (0.5 mM EGTA, 3 mM MgCl2, 60 mM potassium lactobionate, 20 mM taurine, 10 mM KH2PO4, 20 mM HEPES, 110 mM sucrose and 0.1% (w/v) BSA, pH 7.1) at a concentration of 2.5% cell suspension in MiR05 directly in a 2 ml oxygraphy chamber. Respiration was measured in parallel to mitochondrial ROS production (O_2_^−^ and H_2_O_2_) at 37°C in the Oroboros O2k (calibrated for each experiment) equipped with the O2K Fluo-LED2 Module (Oroboros Instruments, Austria). Mitochondrial ROS measurements were performed as described previously^116^ using LEDs for green excitation in the presence of 10 μM Amplex Red, 1 U/ml horseradish peroxidase, and 5 U/ml superoxide dismutase in 2 ml MiR05 per O2K chamber. Calibration was performed by titrations of 5 μL of 40 μM H2O2.

Routine respiration and associated mitochondrial ROS production were assessed from unpermeabilized cells, after which plasma membranes were permeabilized by application of an optimized (optimal accessibility of substrates to mitochondria while maintaining mitochondrial outer membranes intact) concentration of digitonin (5 μg/mL).

A substrate-uncoupler-inhibitor-titration (SUIT) protocol was applied to measure oxygen flux at different respirational states on permeabilized cells, as described previously^117, 118^. Briefly, NADH-pathway (N) respiration in the LEAK and oxidative phosphorylation (OXPHOS) state was analyzed in the presence of malate (2 mM), pyruvate (10 mM) and glutamate (20 mM) before and after the addition of ADP (5 mM), respectively (N*L*, N*P*). Addition of succinate (10 mM) allowed assessment of NADH- and Succinate-linked respiration in OXPHOS (NS*P*) and the uncoupled state (NS*E*) after incremental (Δ0.5 μM) addition of carbonyl cyanide m-chlorophenyl hydrazine (CCCP). Inhibition of Complex I by rotenone (0.5 μM) yielded succinate-linked respiration in the uncoupled state (S*E*). Tissue-mass specific oxygen fluxes were corrected for residual oxygen consumption, *Rox*, measured after additional inhibition of the mitochondrial electron transfer system, ETS, Complex III with antimycin A. For further normalization, fluxes of all respiratory states were divided by ETS-capacity to obtain flux control ratios, *FCR*. Terminology was applied according to http://www.mitoglobal.org/index.php/Gnaiger_2019_MitoFit_Preprint_Arch Mitochondrial ROS values were corrected for background fluorescence, and respirational states before the addition of uncoupler were used for analysis.

### Measurement of the mitochondrial membrane potential and the mitochondrial mass in PFF-treated neurons

Primary hippocampal neurons were plated in a 96-well plate and treated with α-syn PFFs (70 nM). After 7, 14, and 21 days of treatment, mitochondrial membrane potential and mitochondrial mass were quantified using the TMRE (tetramethylrhodamine, ethyl ester) assay kit (Abcam, UK) and Mitotracker Green tracer (Life Technologies, Switzerland) assays respectively.

#### TMRE assay

TMRE is a permeant dye that accumulates in the active mitochondria. TMRE was added to cells at a final concentration of 200 nM and incubated for 20 min 37°C. After a wash with PBS, TMRE staining was quantified using a Tecan infinite M200 Pro plate reader (Tecan, Maennedorf, Switzerland) with excitation and emission wavelengths of 549 nm and 575 nm respectively.

#### Mitotracker Green assay

Mitotracker Green was added to cells at a final concentration of 20 nM and incubated for 30 min at 37°C. After a wash with PBS, TMRE staining was quantified using a Tecan infinite M200 Pro plate reader (Tecan, Maennedorf, Switzerland) with excitation and emission wavelengths of 487 nm and 519 nm respectively.

### Measurement of the synaptic area

Images were acquired on an LSM 700 microscope (Carl Zeiss Microscopy, Germany), with a 40× objective (N.A. = 1.30). The pixel size was set to 78 nm and a z-step of 400 nm; the pinhole was set at 1 µm (equivalent 1A.U, channel 2).

To automatically analyze the z-stack images, the image analysis was performed in Fiji^119^, using a custom script (in ImageJ macro language). Briefly, the script binarizes the different channels of the image using thresholds and measures the area and perimeter. The script binaries user selected channels of the image using a threshold, either a fixed value or a value defined by an automatic threshold method (https://imagej.net/Auto_Threshold) (user defined the appropriate threshold by visual inspection, c1 = DAPI, c2 = Synapsin I, c3 = MAP2, c4 = pS129). From these resulting images, the script applies median filtering to remove noisy pixels and outputs the final masks (user defined the radius of the median filter for each channel, c1 = DAPI, c2 = Synapsin I, c3 = MAP2, c4 = pS129). Finally, the script measures the area, the perimeter, and multiplies the perimeter value by the z-step (to get an estimate of the surface). The generated masks are merged and saved as an output for visual validation of the thresholding.

### Relative Quantification of WBs and Statistical Analysis

The level of total α-syn (15 kDa, 12 kDa, or HMW) or pS129-α-syn were estimated by measuring the WB band intensity using Image J software (U.S. National Institutes of Health, Maryland, USA; RRID:SCR_001935) and normalized to the relative protein levels of actin. All the experiments were independently repeated three times. The statistical analyses were performed using Student’s *t*-test or ANOVA test followed by a Tukey-Kramer *post-hoc* test using KaleidaGraph (RRID:SCR_014980). The data were regarded as statistically significant at p<0.05.

## Acknowledgments

This work was supported by funding from EPFL and UCB (H.A.L, A.L.M, J.B, N.M, and L.W). We thank F.A for the production and the characterization of α-syn *monomers* and fibrils depicted in Figure S1. We are grateful to Dr. Arne Seitz and his staff at the Bio-imaging Core Facility (BioP, EPFL) for their technical support and helpful discussions. We thank Dr Romain Guiet (BioP, EPFL) for the development of a custom script (in ImageJ macro language) used for post-processing imaging analysis. We thank the Proteomics Core Facility (EPFL): Dr Marc Moniatte for the discussions, and Romain Hamelin and Dr. Florence Armand for handling the samples and for their valuable discussions. We thank Dr Gerardo Turcatti and his staff at the Biomolecular Screening Core Facility (EPFL) for the discussions and their technical support. We thank Dr. Bastien Mangeat and his staff at Gene expression Core Facility (EPFL) for their technical support in handling the samples. We also thank Stéphanie Rosset and Anaëlle Dubois at the Bio-EM Core facility (EPFL) for their technical support. Figures 9 and S14 have been prepared partially created using with BioRender.

## Author contributions

H.A.L conceived and supervised the study. H.A.L and A.L.M.M designed all the experiments and wrote the paper. A.L.M.M performed and analyzed the experiments shown in Figures 1-5, 6A, 6E-G, 7-9, and Figures S1B-S13. J.B designed, performed, and analyzed the experiments shown in Figures 6B-D and S14. N.M analyzed the experiments shown in Figures 4 and S8, S12-S13. L.W performed and analyzed the experiments shown in Figures 6G and 7A. M.L. analyzed the transcriptomic data and participated to the Figures 4D, 5, 8G-H, I and S9. F.K developed the pipeline for High Content Image Analysis (HCA) using Cell Profiler software shown in Figures S3 G-I and Figure 8D-F. M.C prepared the samples for CLEM analysis and acquired EM images in Figures 2, 3, and S5. G.K supervised the experiments shown in Figures 1L, 2-3, and S5 and contributed to the interpretation of the data. All authors reviewed and contributed to the writing.

