## Supplemental information for "The process of Lewy body formation, rather than simply alpha-synuclein fibrillization, is the major driver of neurodegeneration in synucleinopathies"

### *The dynamics of Lewy body formation, rather than simply alpha-synuclein fibrillization, is the primary cause of mitochondrial alterations, synaptic dysfunction and neurodegeneration*

***Anne-Laure Mahul-Mellier[1], Johannes Burtscher[1], Niran Maharjan[1], Laura Weerens[1], Marie Croisier[2], Fabien Kuttler[3], Marion Leleu[4], Graham Knott[2], and Hilal A. Lashuel[1]****

[1]Laboratory of Molecular and Chemical Biology of Neurodegeneration, Brain Mind Institute, Ecole Polytechnique Fédérale de Lausanne (EPFL), 1015 Lausanne, Switzerland. [2]BioEM Core Facility and Technology Platform, EPFL, Lausanne 1015, Switzerland. [3]BiomolecularScreening Core Facility and Technology Platform, EPFL, Lausanne 1015, Switzerland. [3] Gene expression Core Facility and Technology Platform, EPFL, Lausanne 1015, Switzerland.
*To whom correspondence should be addressed at: Laboratory of Molecular and Chemical

Biology of Neurodegeneration, Brain Mind Institute, Ecole Polytechnique Fédérale de Lausanne, 1015 Lausanne. Tel: +41216939691, Fax: +41216939665,

**Running title:**

The grammar of Lewy body formation.

**Keywords:** alpha-synuclein (α-synuclein), Parkinson’s disease, aggregation, post-translational modification (PTM), mitochondria, synapse

#### Supplemental information – Material and Methods

##### Preparation of α-syn seeds from PFF-treated primary hippocampal neurons

70 nM of mouse α-syn PFFs were added to hippocampal primary neurons at *DIV* 5-6. After 10 days of treatment, seeded neurons were lysed in 1% Triton X-100/ Tris-buffered saline (TBS) (50 mM Tris, 150 mM NaCl, pH 7.5) supplemented with protease inhibitor cocktail, 1 mM phenylmethane sulfonyl fluoride (PMSF), and phosphatase inhibitor cocktail 2 and 3 (Sigma-Aldrich, Switzerland) and sequential biochemical fractionation of cell extracts was performed as described previously[1]. After sonication using a fine probe [(0.5-s pulse at an amplitude of 20%, ten times (Sonic Vibra Cell, Blanc Labo, Switzerland], cell lysates were incubated on ice for 30 min and centrifuge at 100,000 g for 30 min at 4°C. The supernatant (soluble fraction) was collected while the pellet was washed in 1% Triton X-100/TBS, sonicated as described above, and centrifuged for another 30 min at 100,000 g. The supernatant was discarded whereas the pellet (insoluble fraction) was washed twice with PBS, pH 7.4. The pellet was then sonicated in PBS, pH 7.4 using a fine probe [(0.5-s pulse at an amplitude of 20%, ten times (Sonic Vibra Cell, Blanc Labo, Switzerland]. The bicinchoninic acid (BCA) protein assay was performed to quantify the protein concentration in the soluble and insoluble fractions. 2 µg of the insoluble fraction was then added to hippocampal primary neurons at *DIV* 5-6 to induce seeding.

**Supplemental information – Titles of the Figures and Legends**

**Figure S1. Preparation and characterization of recombinant monomeric and PFFs α-syn species (related to Figures 1 to 8)**

**A. Purity and characterization of α-syn monomers.**

Recombinant WT mouse α-syn was produced in *E. coli* and purified by anion exchange chromatography and size-exclusion chromatography, followed by a final chromatographic step using reverse-phase HPLC, as previously described[2]. The purity of recombinant monomeric α-syn after purification was assessed by ESI-LC/MS, which showed the expected mass.

**B-E. Purity and characterization of α-syn fibrils.**

α-syn fibrils were formed by incubation of monomeric α-syn for 5 days at 37°C under constant agitation at 1000 rpm. (**B**). After sonication, fibril formation was assessed by ThT fluorometry. All data represent the average ± SD (n=3). (**C**). Purity of α-syn fibrils was verified by SDS-PAGE and Coomassie blue staining. After sonication, fibril preparations were centrifuged, and the presence of the fibrils was verified in the pellet fraction, while the absence of monomer release after the sonication step was assessed in the supernatant fraction or after filtration through a 100 kDa filter (filtration). (**D-E**). α-syn fibrils were characterized by transmission electron microscopy (TEM) imaging. (**D**) Representative images of negatively stained α-syn fibrils before and after sonication. All α-syn fibrils showed the characteristic rigid non-branched fibrillar morphology. Scale bars = 100 nm. (**E**) Average length of the fibrils after sonication.

**F. Treatment of hippocampal primary neurons by α-syn PFFs seeds.**

Schematic of experiments for the time course of the primary culture treatment with α-syn PFFs seeds. Morphological[3-5], physiological[6-8], gene expression[9] and proteome[10] changes are observed during the different developmental stages of primary neurons in culture (Days *In vitro*, DIV). Therefore, we adapted the time-course of the PFFs treatment in primary neurons to be able to analyse the spatio-temporal effects of the seeding mechanism at the biochemical, proteomic, transcriptomic and ultrastructural level uncoupled to the intrinsic changes related to neuronal development and maturation. PFFs were added to the neuronal cell culture media at different DIV and PFFs-treated neurons were all harvested at the DIV 26. This ensures that the developmental stage of primary neurons in culture was similar at the harvest time.

**G. Image analysis workflow.** Images were analyzed using CellProfiler software. **a.** Raw images were acquired in green channel (MAP2 staining), blue channel (DAPI) and red channel (pS129 staining) for each well and each field of view. **b.** An RGB color image was produced and rescaled from individual grayscale images, MAP2 in green, DAPI in blue and pS129 in red, for visualization purpose. **c.** Bulk of neurons area was first identified based on MAP2 staining using otsu thresholding. **d.** MAP2 image was masked to exclude all non-neuronal areas, based on bulk neurons segmentation, and tubeness method was used to enhance neurites. **e.** DAPI image was rescaled and masked to exclude all non-neuronal areas, therefore excluding all glial cells from the analysis, and neuronal nuclei were segmented using minimum cross entropy thresholding. **f.** Neurites are then segmented on image **d** using minimum cross entropy thresholding and using a watershed algorithm with neuronal nuclei as seeds. Cell Bodies are identified as the perinuclear region by expanding by 3 pixels the previously identified neuronal nuclei. **g.** Image from pS129 staining is rescaled and used to segment individual a-Syn aggregates in neurites and cell bodies using the segmentation from **f**. **h.** Features are then extracted and quantified from all identified objects: nuclei, neurons, neurites, cell bodies, aggregates in neurites, aggregates in cell bodies. Measurements were made on pS129 channel and composed of counts, size, shape and intensity criteria. **i.** For control / visualization purpose, an overlay image was finally created and recorded, composed of the color image from **b** and the outlines of some segmented objects: cell bodies and a-Syn aggregates in cell bodies and neurites.

**Figure S2. List of the antibodies used in this study (related to Figures 1 to 8)**

**A**. Antibodies used for the detection of total α-syn.

**B**. Antibodies used for the detection of α-syn phosphorylated on S129 residue.

**C**. Other antibodies used in the study.

**D**. Secondary antibodies or dyes used for immunoblotting or confocal imaging.

**Figure S3. Morphological and spatial distribution changes of α-syn seeded aggregates formed in PFFs-treated neurons overtime (related to Figure 1)**

**A.** 70 nM of mouse sonicated PFFs were added to neurons at DIV 5 (days *in vitro*). α-syn aggregates appear first in the neuronal extension after 4 days of PFFs-treatment. Aggregates were detected by ICC using pS129 (MJFR13). Neurons were counterstained with microtubule-associated protein (MAP2) antibody, and the nucleus was counterstained with DAPI staining. Scale bars = 10 μm.

**B-D.** Temporal analysis of α-syn aggregates formed at D7 (**B**), at D14 (**C**) and at D21(**D**) in WT neurons after addition of α-syn PFFs. Aggregates were detected by ICC using pS129 (MJFR13, 81a or GenTex antibodies) in combination with total α-syn (epitopes: 1-20 or 34-45 or NAC/ or 108-120) antibodies. Neurons were counterstained with microtubule-associated protein (MAP2) antibody, and the nucleus was counterstained with DAPI staining. Scale bars = 10 μm.

**E.** Quantification of the different types of morphologies observed by ICC (**D**) for α-syn seeded aggregates in PFFs-treated neurons at D21. A minimum of 300 α-syn seeded aggregates were measured in three independent experiments. $p<0.01$=*, $p<0.0001$=*** (ANOVA followed by Tukey HSD post-hoc test, filamentous vs ribbon-like vs round LB-like inclusions).

**F-I.** Spatial distribution of the α-syn seeded aggregates formed in PFFs-treated neurons overtime was assessed by high content imaging analysis (HCA) that allow the total count of α-syn seeded aggregates (**F**) in neurites (**G**) or in cell bodies (**H**) overtime in PFFs-treated primary culture. **I**. Total level of pS129 in MAP2 positive neurons (neurites + cell bodies).

For each independent experiment, duplicated wells were acquired per condition, and nine fields of view were imaged for each well. Images were then analysed using Cell Profiler software to identify and quantify the number of α-syn seeded aggregates in neuronal cell bodies (DAPI and MAP2-positive cells) or in neurites (MAP2-positive).

The graphs (**G-I**) represent the mean +/- SD of three independent experiments. $p<0.01$=*, $p<0.01$=**, $p<0.0001$=*** (ANOVA followed by Tukey HSD post-hoc test, PBS vs PFFs-treated neurons). p<0.001= ##, p<0.001= ### (ANOVA followed by Tukey HSD post-hoc test, PFFs-treated neurons D14 vs D21).

**J-O.** Colocalization of lipids with pS129-positive inclusions at D21. Ceramide (**J**), neutral lipid (**K**), cholesteryl (**L**), phospholipids (**M**), methyl ester (**N**) and sphingomyelin (**O**) were stained using specific fluorescent probes (Figure S2E). pS129-positive aggregates were stained with MJFR13 antibody. Neurons were counterstained with microtubule-associated protein (MAP2) antibody, and the nucleus was counterstained with DAPI staining. Scale bars = 10 µm.

**Figure S4. α-syn seeds prepared from PFF-seeded primary neurons have high seeding activity (related to Figure 1)**

**A.** 70 nM of mouse α-syn PFFs were added to hippocampal primary neurons at *DIV* 5-6. After 14 days of treatment, seeded neurons were lysed and sequential biochemical fractionation of cell extracts was performed. The pellet corresponding to the insoluble fraction was resuspended in PBS and dispersed by sonication. 2 µg of this fraction was then added to naïve hippocampal primary neurons at *DIV* 5-6 for 14 days.

**B.** Newly formed fibrils were detected by ICC using pS129 (81a antibody) in combination with total α-syn (epitope: 1-20) antibody. Neurons were counterstained with microtubule-associated protein (MAP2) antibody, and the nucleus was counterstained with DAPI staining. Neurons highlighted in yellow in the top panel was observed at higher magnification (bottom panel). Top panel, scale bar = 40 µm. Bottom panel, scale bar = 10 µm.

**Figure S5. Temporal profiling of the LB-like inclusions formation by correlative light electron microscopy (related to Figure 2)**

**A.** PFFs were added for 7, 14, and 21 days to the extracellular media of hippocampal neurons plated on dishes with alpha-numerical searching grids imprinted on the bottom, allowing an easy localization of the cells. Neurons were fixed at the indicated time and imaged by confocal microscopy (**C-E**, top images, scale bars = 10 µm). The selected neurons were embedded and cut by an ultramicrotome. Serial sections were examined by TEM.

**B.** Graph representing the mean +/- SD of the width of the microtubules compared to the newly formed fibrils at D7 (a minimum of 120 microtubules or newly formed fibrils were counted). Measurements confirmed that the width of newly formed α-syn fibrils of 11.94±3.97 nm (SD) is significantly smaller than the average width of the microtubules of 18.67±2.94 nm (SD). $p<0.001$=*** (student t-test for unpaired data with equal variance), indicating that this parameter can be used to discriminate the newly formed fibrils from the cytoskeletal proteins.

**C-E**. Representative confocal images (top images) and EM micrographs (bottom images) of the control neurons treated with PBS buffer for 7 days (**C-D**) or neurons treated with PFFs for 14 days (**E**). Microtubule is highlighted in orange (**C**), and nucleus is highlighted in blue. **E.** CLEM imaging confirmed that PFFs did not accumulate on the outer side of the plasma membrane when used at a nanomolar concentration, as previously shown when higher concentrations were applied to the neurons[11].

**F.** Raw EM images of α-syn inclusions formed at D21 shown in Figure 2F.

**G-J.** The length of newly formed fibrils was measured over time. A minimum of 160 fibrils were counted for each condition.

**C-D.** Scale bar = 500 nm. **E**. Scale bar = 5 μm **F.** Scale bar = 1 μm

**Figure S6. LB-like inclusions imaged by correlative light electron microscopy at D21 (related to Figure 3)**

**A-D.** Raw EM images of α-syn inclusions formed at D21 shown in Figure 3A-D.

**A-B, D.** Scale bars = 1 μm. **C**. Scale bar = 2 μm.

**Figure S7. WB analysis of α-syn seeded-aggregates formed 7, 14 or 21 days after adding mouse PFFs to WT neurons (related to Figures 1-3)**

Control neurons were treated with PBS. After sequential extractions, the insoluble fractions of neuronal cell lysates were analyzed by immunoblotting. Total α-syn, pS129 and actin were respectively detected by SYN-1, pS129 (MJFR13), and actin antibodies. The higher molecular weights (HMWs) corresponding to the newly formed fibrils are detected from 25 kDa to the top of the gel. A minimum of three independent experiments was performed. p<0.001=**, p<0.0001=*** (ANOVA followed by Tukey HSD post-hoc test, PBS vs D7 or D14 or D21).

**Figure S8. Temporal proteomic analyses of the protein contents found in the insoluble fraction of the PFFs-treated neurons reveals a high increase in proteins related to the endomembrane system (related to Figure 4)**

**A.** Insoluble proteins from neurons treated with PBS and PFFs for 7 days were extracted and analyzed using LC-MS/MS. Identified proteins were plotted using volcano plot. Mean difference ($log_2$) between the insoluble fractions of PFFs-treated neurons and PBS neurons treated for 7 days, were plotted against $–log_{10}$ P value (T-Test). Dotted lines represent the FDR <0.05 (False Discovery Rate) and threshold of significance SO=1 assigned for the subsequent analysis. At D7, only α-syn and Plcb1 proteins were significantly enriched in the insoluble fraction of the PFFs-treated neurons.

**B.** Canonical pathways enriched in the insoluble fractions of the PFFs-treated neurons at D14 and D21 using Ingenuity Pathway Analysis (IPA).

**C.** Comparison of our proteomic results with previous published proteomic analyses using the same neuronal seeding model**.** 8 out of the 12 proteins shown by Henderson et al[12] to be significantly enriched in the insoluble fraction of PFFs-treated neurons were also present in our proteomic data.

**Figure S9. Gene expression level changes during the formation of the newly formed fibrils and their maturation into LB-like inclusions (related to Figure 5)**

**A.** Temporal transcriptomic analysis of the gene expression level in PBS-treated neurons vs PFFs-treated neurons treated for 7, 14 or 21 days. Genes with an absolute log2 fold-change greater than 1 and an FDR less than 0.01 were considered as significantly differentially expressed. The table depicts the number of genes up or down-regulated in PFFs-treated neurons overtime.

**B.** The heatmap depicts the synaptic genes that are up- or down-regulated in PFFs-treated neurons between D14 and D21. “+” and “-“ indicate respectively a significant upregulation or downregulation in the gene expression level. (T-Test, D14 PFFs-treated neurons vs D21 PFFs-treated neurons).

**Figure S10. Addition of α-syn PFFs to primary neurons does not induce cell death in KO neurons (related to Figure 8)**

**A-B.** Cell death level was assessed in KO neurons treated with PFFs (70 nm) for up to D21 (**A**) using lactate dehydrogenase (LDH) release assay (**B**). For each independent experiment, triplicate wells were measured per condition. A minimum of three independent experiments was performed.

**Figure S11. The proteome of the neuronal LB-like inclusions overlaps significantly with that of LBs from human PD brain tissues (related to Figure 4 and Discussion)**

The table depicts the comparison of the proteins enriched in the insoluble fraction of PFFs-treated neurons (at D14 and D21) to the proteins contents of human *bona fide* LBs established by IHC[13] or by proteomic[14,15] analyses.

**Figure S12. Temporal proteomic analyses of the protein contents found in LB-like inclusions formed in PFFs-treated neurons (related to Figure 4 and Discussion)**

The table depicts the proteins involved in the protein degradation machinery that were enriched in the insoluble fraction of PFFs-treated neurons at D14 and D21.

**Figure S13. The dynamics of Lewy body formation, rather than simply alpha-synuclein fibrillization, is the primary cause of mitochondrial alterations, synaptic dysfunction and neurodegeneration (related to Figures 1-8 and discussion)**

The table recapitulates the main events that occur during the formation of α-syn seeded aggregates and their maturation into LB-like inclusions overtime.

**Figure S14. Model of how the interaction of mitochondria and LB-like inclusions might induce neurodegeneration (related to Figure 6 and discussion)**

Treatment of neurons with α-syn PFFs induces formation of fibrillar newly formed α-syn aggregates.

Aα-syn aggregates interact with mitochondrial membranes, mitochondrial transmembrane proteins and soluble α-syn localized in or close to mitochondria, accelerating α-syn aggregation and the formation of LB-like inclusions. Aggregation kinetics and perturbation of mitochondrial membrane integrity (potentially caused by mitochondrial reactive oxygen species) induce damage signalling that on D7 is translated in transcriptional upregulation of both pro-apoptotic factors (e.g. BAX, Aifm3) and potentially protective factors, including subunits of mitochondrial complex I, as well as mitophagy-related PINK1 and oxidative response genes (e.g. Gpx1). These processes are accompanied by reduced mitochondrial reactive oxygen species (ROS) generation and maintenance of efficient oxidative phosphorylation.

**B.** Continuous recruitment of α-syn to mitochondria and stimulated aggregation in presence of mitochondrial membranes cause maturation of LB-like inclusions with detectable sequestration of mitochondrial proteins, such as mitofusin2 (MFN2) or proteins related to oxidative phosphorylation (OXPHOS), and proteins related to REDOX-mechanisms. Cellular resilience is still sufficient to secure oxidative phosphorylation performance.

**C**. Aggregating α-syn mechanically disrupts mitochondrial membranes resulting in dense assemblies of fibrillary α-syn, mitochondrial components and other sequestered constituents of LB-like inclusions. Increasing recruitment of mitochondria to inclusions renders the organelles dysfunctional and results in massive respirational deficits, associated with reduced levels of numerous mitochondrial proteins and the induction of the apoptotic machinery.

**Figure S1. Related to Figures 1-8**

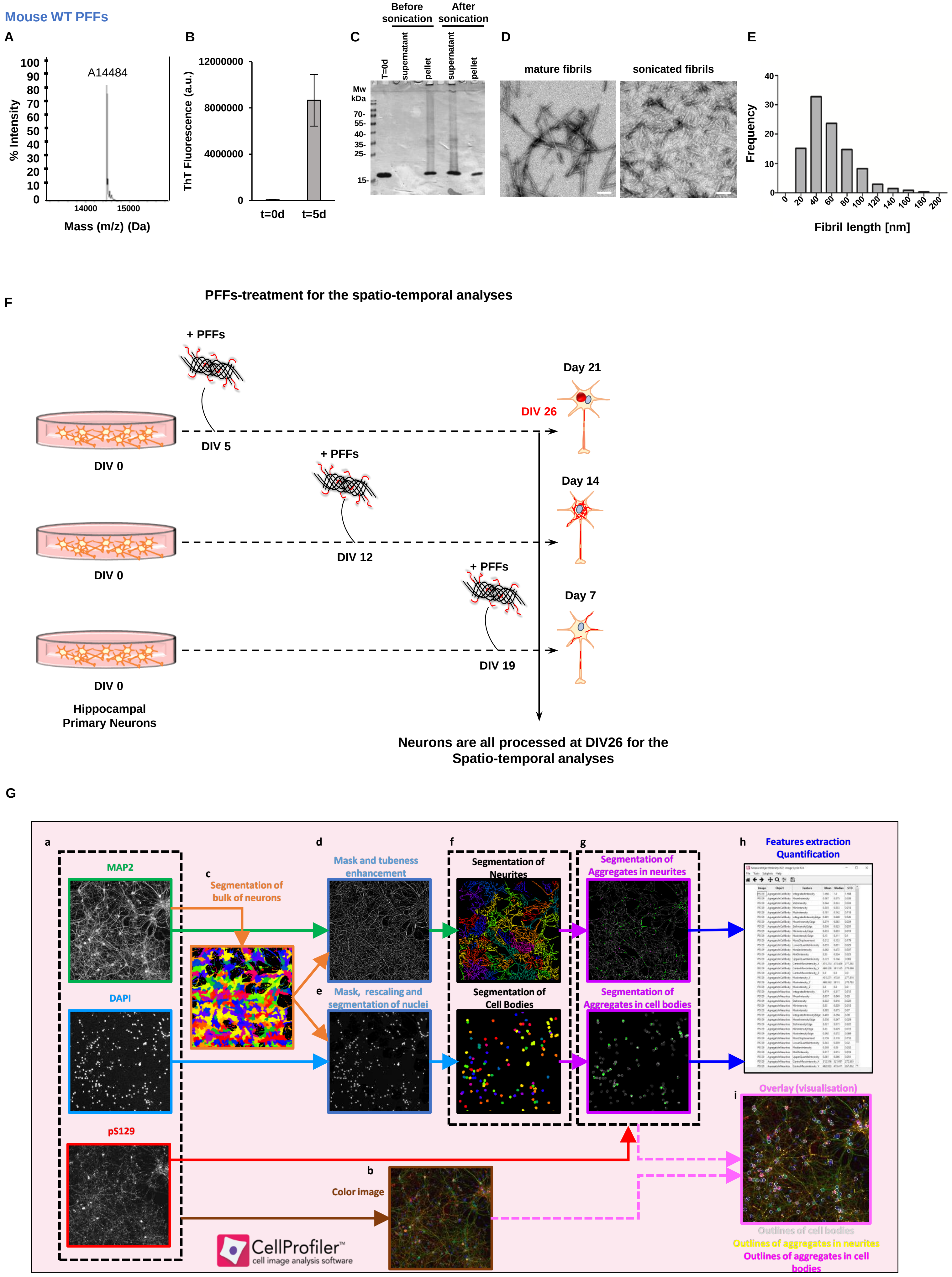

#### Figure S2. Related to Figures 1 to 8

**A**

| Primary Antibody | Catalog # | Company | Clone | RRID | Host | Concentration | WB dilution | ICC dilution | Epitope |
|---|---|---|---|---|---|---|---|---|---|
| anti-α-syn **total** | LASHUEL | - | LASH-EGT1-20 | - | Rabbit | Not provided | 1:1000 | 1:1000 | **1-20** |
| anti-α-syn **total** | LASHUEL | - | LASH-BL-A15110B | - | Mouse | 1.2 mg/ml | 1:1000 | 1:500 | **34-45** |
| anti-α-syn **total** | LASHUEL | - | LASH-BL-A15115A | - | Mouse | 0.8 mg/ml | 1:1000 | 1:500 | **80-96** |
| anti-α-syn **total** | 610787 | BD | SYN-1 | RRID:AB_398108 | Mouse | 0.25 mg/ml | 1:1000 | 1:1000 | **91-99** |
| anti-α-syn **total** | AB5336P | Millipore | - | RRID:AB_2192954 | Sheep | 1 mg/ml | 1:500 | 1:500 | **108-120** |
| anti-α-syn **total** | ab131508 | Abcam | - | RRID:AB_11155736 | Rabbit | 1 mg/ml | 1:500 | 1:500 | **134-138** |

**B**

| Primary Antibody | Catalog # | Company | Clone | RRID | Host | Concentration | WB dilution | ICC dilution | Epitope |
|---|---|---|---|---|---|---|---|---|---|
| anti-**pS129**-α-syn | Ab168381 | Abcam | MJF-R13 | RRID:AB_2728613 | Rabbit | 4.229 mg/ml | 1:3000 | 1:1000 | Not provided |
| anti-**pS129**-α-syn | 825701 | BioLegend | P-syn/81A | RRID:AB_2564891 | Mouse | 1.0 mg/ml | 1:1000 | 1:1000 | AYEMPpSEEGYQ |
| anti-**pS129**-α-syn | GTX82738 | GeneTex | - | AB_11176838 | Rabbit | - | 1:500 | 1:1000 | Synthetic phospho-peptide corresponding to amino acid residues surrounding Ser129 conjugated to KLH |

**C**

| Primary Antibody | Catalog # | Company | Clone | RRID | Host | Concentration | WB dilution | ICC dilution | Epitope |
|---|---|---|---|---|---|---|---|---|---|
| anti-**actin** | ab6276 | Abcam | AC-15 | RRID:AB_2223210 | Mouse | 2.2 mg/ml | 1:5000 | Not tested | DDDIAALVIDNGSGK |
| anti-**MAP2** | ab92434 | Abcam | - | RRID:AB_2138147 | Chicken | Not provided | Not tested | 1:2000 | Recombinant full length protein |
| anti-**p62** | H00008878 | Abnova | 2C11 | RRID:AB_437085 | Mouse | 1 mg/ml | 1:1000 | 1:500 | Raised against a full length recombinant SQSTM1 |
| anti-**LC3** | ab48394 | Abcam | - | - | Rabbit | 1 mg/ml | 1:500 | 1:500 | A synthetic peptide made to an Nterminal portion of the human LC3 protein sequence (between residues 1-100) |
| anti-**ubiquitin** | Sc-8017 | Santa-Cruz | P4D1 | RRID:AB_628423 | Mouse | 0.2 mg/ml | 1:500 | 1:500 | 1-76 |
| anti-**OXPHOS** | ab110413 | Abcam | - | - | Mouse | 1.5 mg/ml | 1:1000 | - | Cocktail of high quality antibodies for analyzing relative levels of OXPHOS complexes |
| anti-**Tom 20** | sc-17764 | Santa-Cruz | F-10 | RRID:AB_628381 | Mouse | 0.2 mg/ml | 1:500 | 1:200 | Raised against amino acids 1-145 |
| anti-**Tim 23** | sc-514463 | Santa-Cruz | H-8 | - | Mouse | 0.2 mg/ml | 1:500 | 1:200 | Raised against amino acids 31-209 |
| anti-**VDAC1** | ab14734 | Abcam | 20B12AF2 | - | Mouse | 1 mg/ml | 1:500 | 1:200 | Recombinant full length protein corresponding to Human VDAC1/ Porin. |
| anti-**OPA1** | 612606 | BD biosciences | 18/OPA1 | RRID:AB_399888 | Mouse | 0.25 mg/ml | 1:1000 | - | Raised against amino acids 708-830 |
| anti-**Citrate Synthase** | ab96600 | Abcam | - | - | Rabbit | 1.03 mg/ml | 1:1000 | - | Recombinant fragment within Human Citrate synthetase aa 44-316 |
| anti-**Mitofusin 2** | ab50843 | Abcam | - | - | Rabbit | 1.1 mg/ml | 1:1000 | - | Synthetic peptide corresponding to Human Mitofusin 2 aa 557-576 |
| anti-**Synapsin I** | ab64581 | Abcam | - | - | Rabbit | 1 mg/ml | 1:1000 | 1:500 | Synthetic peptide corresponding to Rat Synapsin I aa 600-700 |
| anti-**PSD95** | MAB N68 | Millipore | K28/43 | - | Mouse | 1 mg/ml | 1:1000 | 1:500 | Recombinant protein corresponding to human PSD95 |
| anti-**Synaptophysin** | ab8049 | Abcam | SY38 | - | Mouse | 1 mg/ml | 1:1000 | 1:500 | Not provided |
| anti-**ERK** | 4696 | Cell signalling | L34F12 | - | Mouse | - | 1:1000 | - | Synthetic peptide corresponding to the sequence of p42 MAP Kinase |
| anti-**p-ERK** | 9101 | Cell signalling | - | - | Rabbit | - | 1:1000 | - | Synthetic phosphopeptide corresponding to residues surrounding Thr202/Tyr204 of human p44 MAP kinase |
| anti-**LAMP1** | ab62562 | Abcam | - | RRID:AB_2134489 | Rabbit | | - | 1:500 | A 15 amino acid peptide from near the centre of human LAMP1 |
| anti-**BIP/Grp78** | ab21685 | Abcam | - | RRID:AB_2119834 | Rabbit | 1 mg/ml | 1:1000 | 1:500 | - |

**D**

| Secondary Antibody | Catalog # | Company | RRID | Concentration | WB dilution | ICC dilution |
|---|---|---|---|---|---|---|
| Goat anti-**mouse Alexa Fluor 680** | A21058 | Invitrogen | RRID:AB_2535724 | 2 mg/ml | 1:5000 | - |
| Goat anti-**rabbit Alexa Fluor 680** | A21109 | Invitrogen | RRID:AB_2535758 | 2 mg/ml | 1:5000 | - |
| Goat anti-**mouse Alexa Fluor 800** | 926-32210 | Li-Cor | RRID:AB_621842 | 1 mg/ml | 1:5000 | - |
| Goat anti-**rabbit Alexa Fluor 800** | 926-32211 | Li-Cor | RRID:AB_621843 | 1 mg/ml | 1:5000 | - |
| Goat anti-**mouse Alexa Fluor 647 + nanogold particles** | 7502 | Nanoprobes | - | | - | 1:800 |
| Donkey anti-**mouse Alexa Fluor 568** | A10037 | Invitrogen | RRID:AB_2534013 | 2 mg/ml | - | 1:800 |
| Donkey anti-**rabbit Alexa Fluor 568** | A10042 | Invitrogen | RRID:AB_2534017 | 2 mg/ml | - | 1:800 |
| Donkey anti-**rabbit Alexa Fluor 647** | A31573 | Invitrogen | RRID:AB_2536183 | 2 mg/ml | - | 1:800 |
| Donkey anti-**mouse Alexa Fluor 647** | A31571 | Invitrogen | RRID:AB_162542 | 2 mg/ml | - | 1:800 |
| Donkey anti-**Sheep Alexa Fluor 647** | A21448 | Invitrogen | RRID:AB_1500712 | 2 mg/ml | - | 1:800 |
| Donkey anti-**chicken Alexa Fluor 405** | 703-475-155 | Jackson Immunoresearch | RRID:AB_2340373 | 1 mg/ml | - | 1:400 |
| Donkey anti-**chicken Alexa Fluor 488** | 703-545-155 | Jackson Immunoresearch | RRID:AB_2340375 | 1 mg/ml | - | 1:400 |
| Goat anti-**chicken Alexa Fluor 568** | A11041 | Invitrogen | RRID:AB_2534098 | 2 mg/ml | - | 1:500 |
| Donkey anti-**chicken Alexa Fluor 647** | 703-605-155 | Jackson Immunoresearch | RRID:AB_2340379 | 1 mg/ml | - | 1:500 |

#### Figure S2. Related to Figures 1 to 8

**E**

| Dyes | Catalog # | Company | RRID | Concentration | WB dilution | ICC dilution |
|---|---|---|---|---|---|---|
| Amytracker**680** | Amytracker™680 | Ebba Biotech | - | | - | 1:500 |
| Mitotracker **Green FM** | M7514 | Invitrogen | - | 1 mg/ml | - | 20 nM |
| Neutral lipid **Lipidox Red** | H34476 | Invitrogen | - | | - | 1:500 |
| Ceramide **BODIPY TR** | B34400 | Invitrogen | - | | - | 1:500 |
| Phospholipid **Lipidox Red** | H34158 | Invitrogen | - | | - | 1:500 |
| Cholesteryl **BODIPY 524/563** | C12680 | Invitrogen | - | | - | 1:500 |
| C12-Sphingomyelin **BODIPY FL** | D7711 | Invitrogen | - | | - | 1:500 |
| Methyl Ester **BODIPY FL** | C34556 | Invitrogen | - | | - | 1:500 |

Figure S3. Related to Figure 1

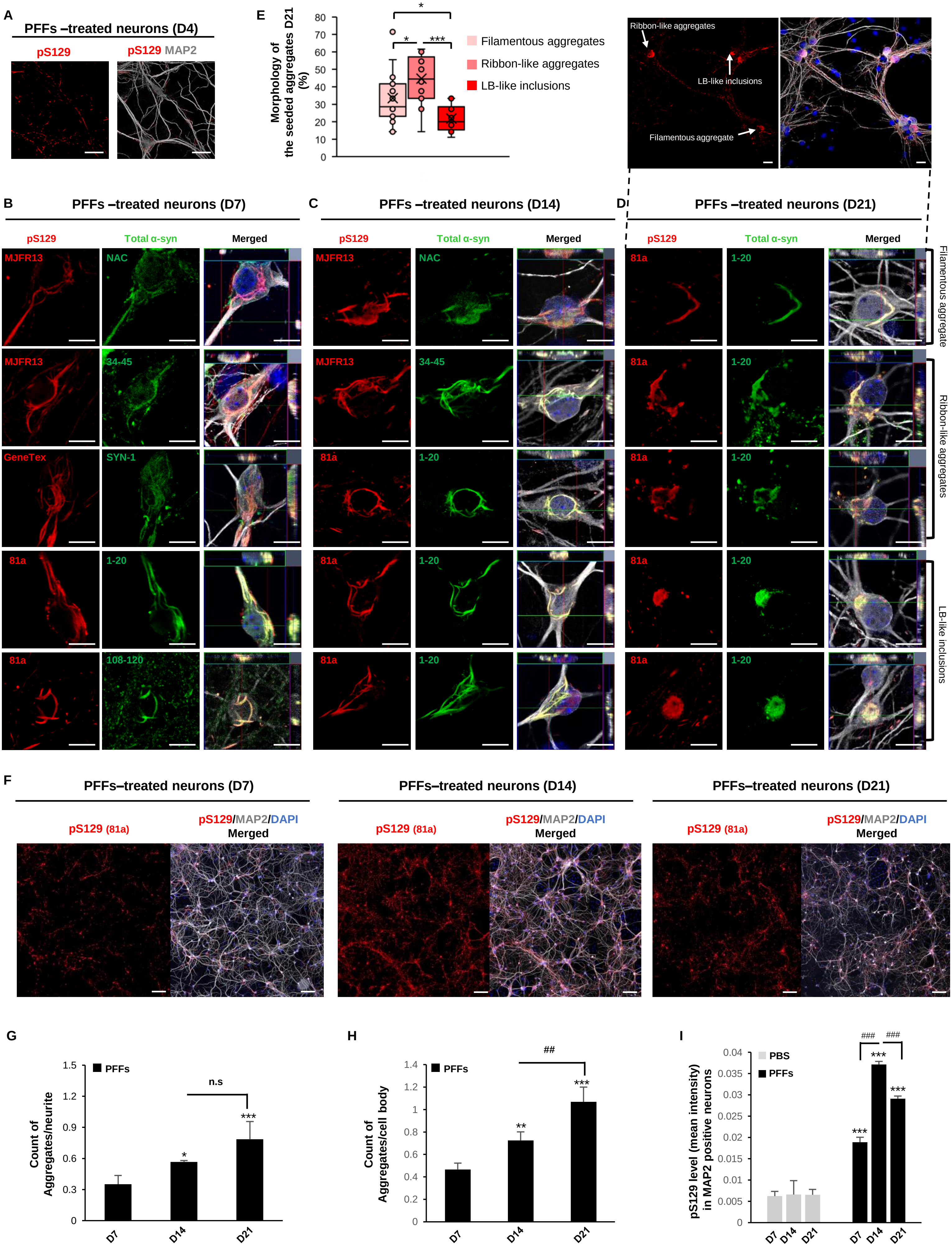

Figure S3. Related to Figure 1

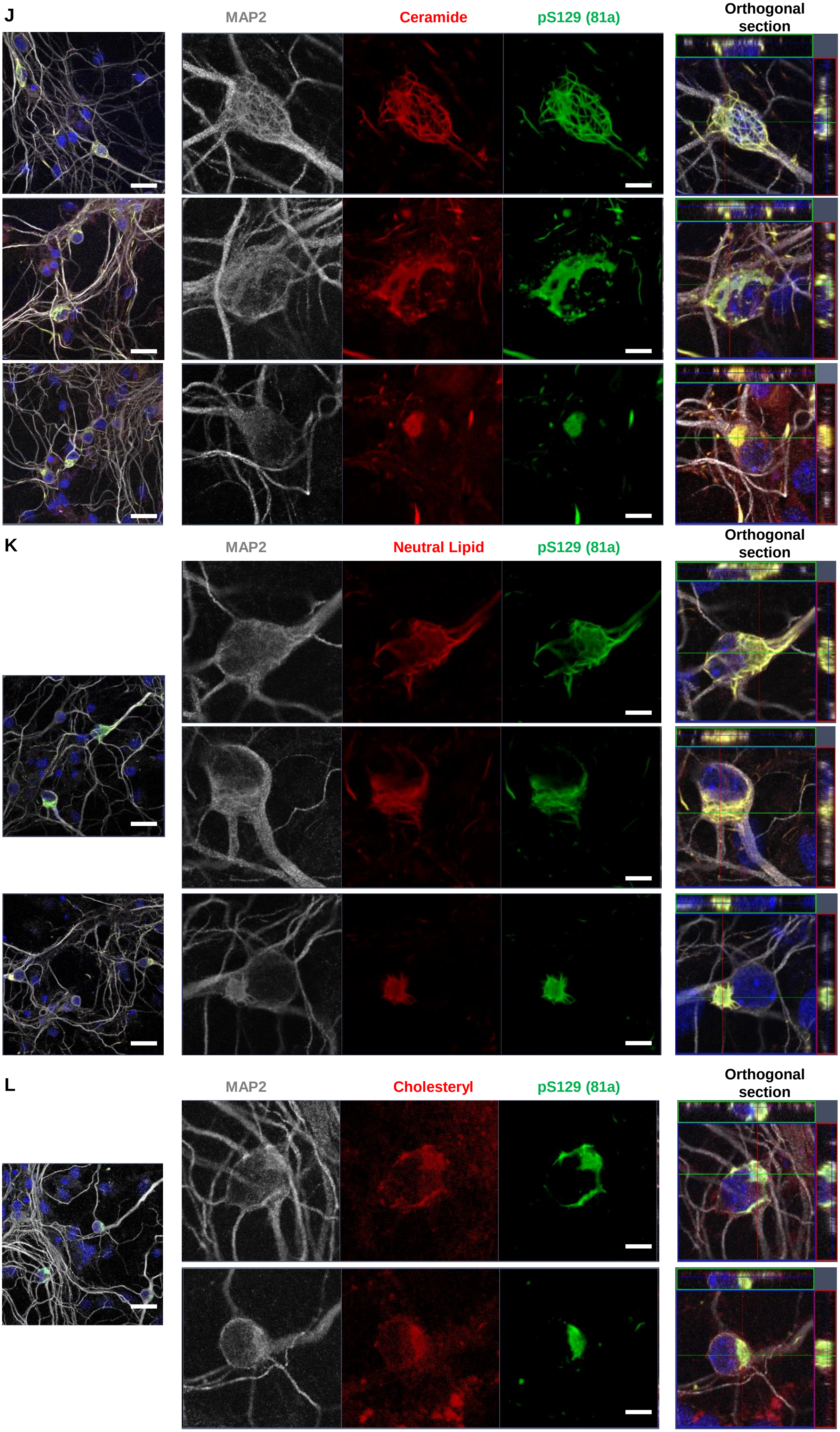

Figure S3. Related to Figure 1

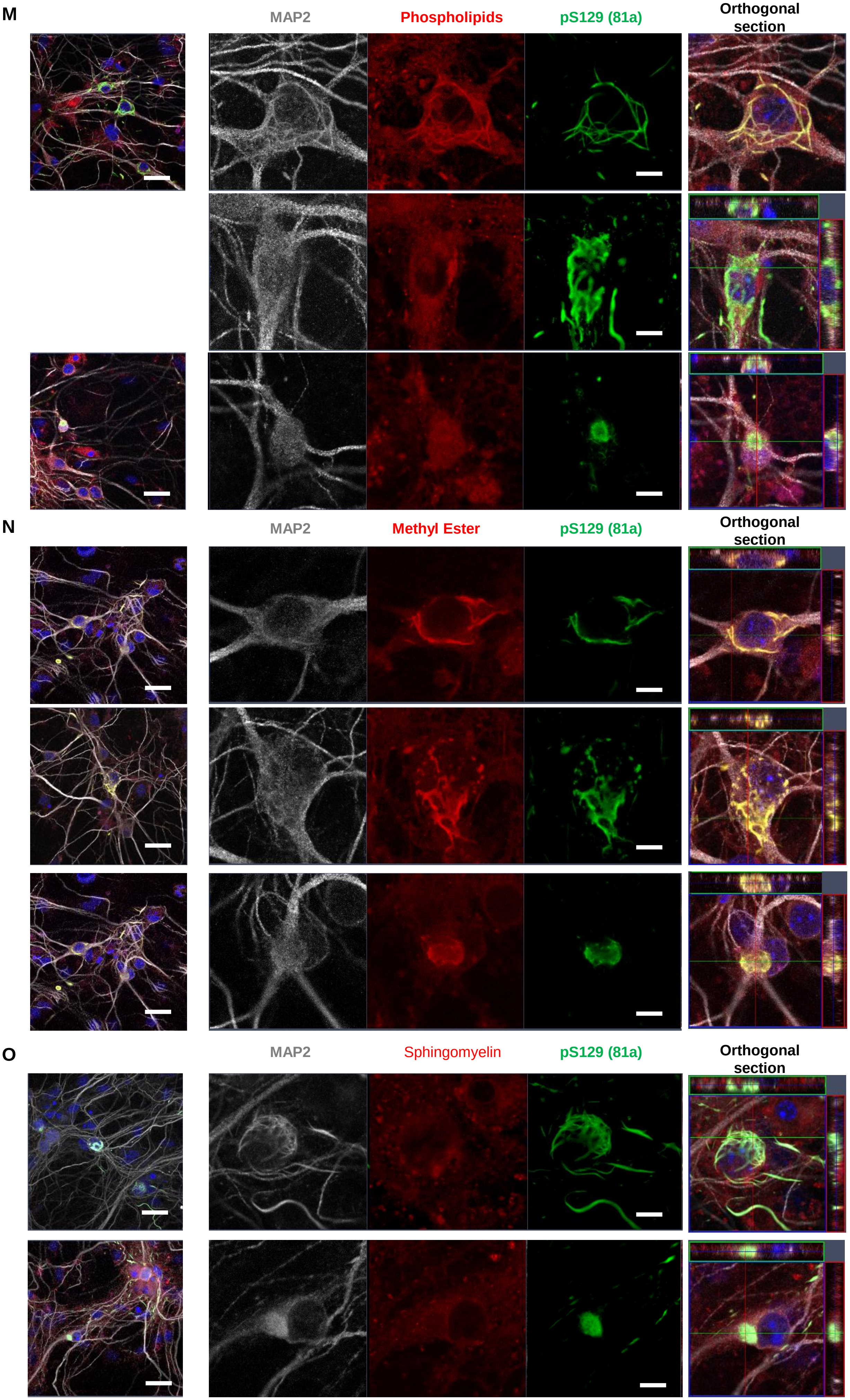

Figure S4. Related to Figure 1

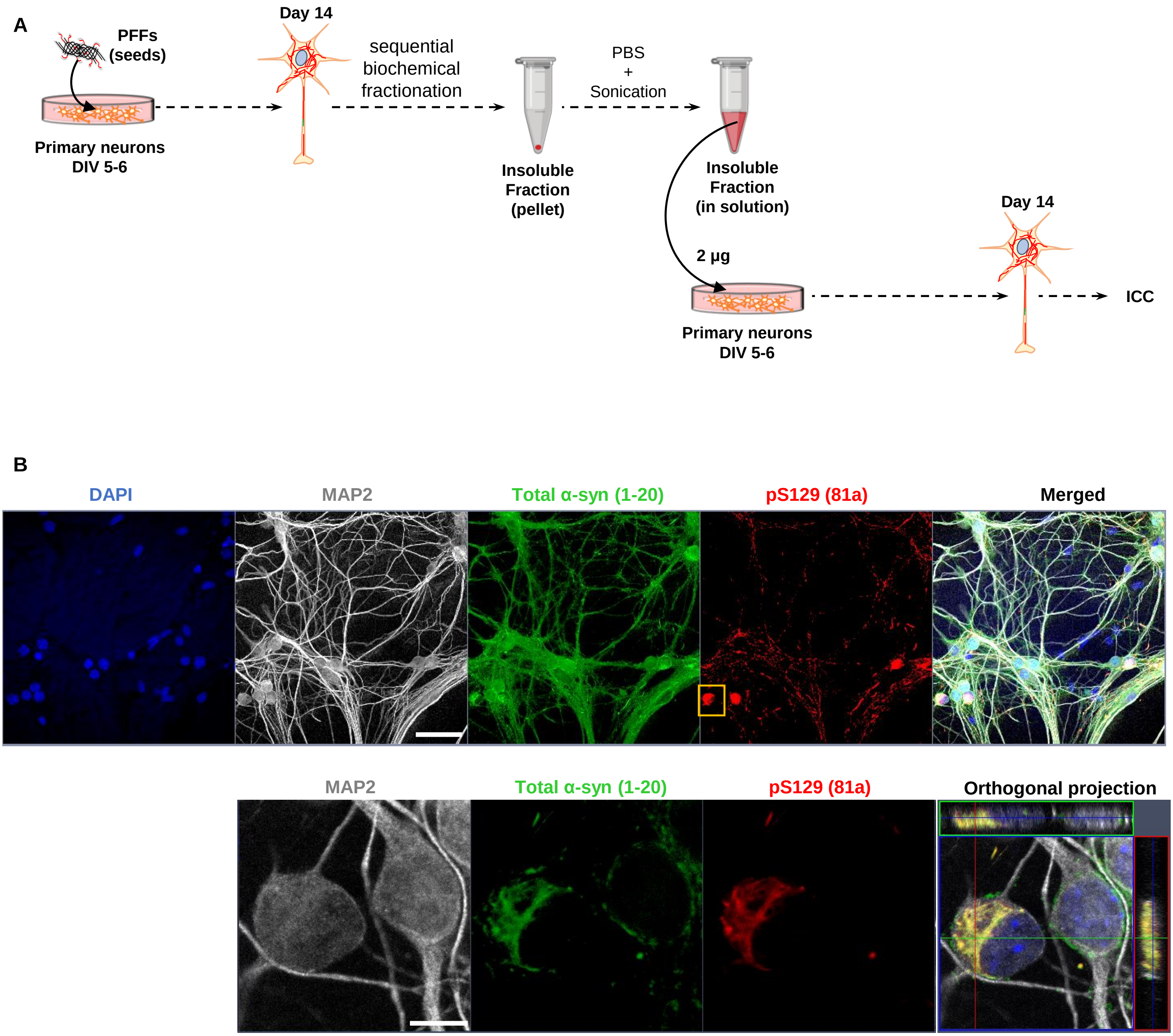

Figure S5. Related to Figure 2

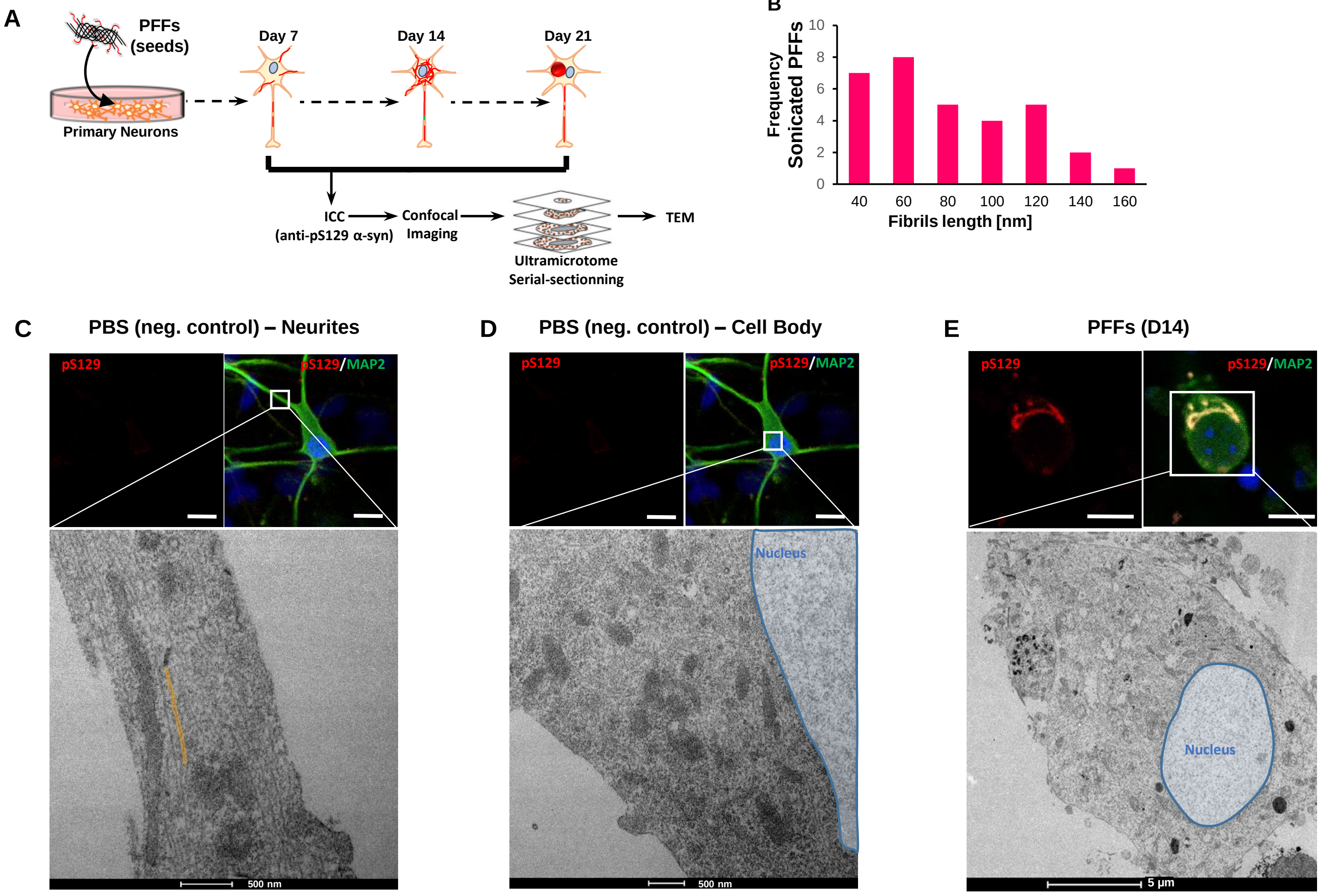

Figure S5. Related to Figure 2

F

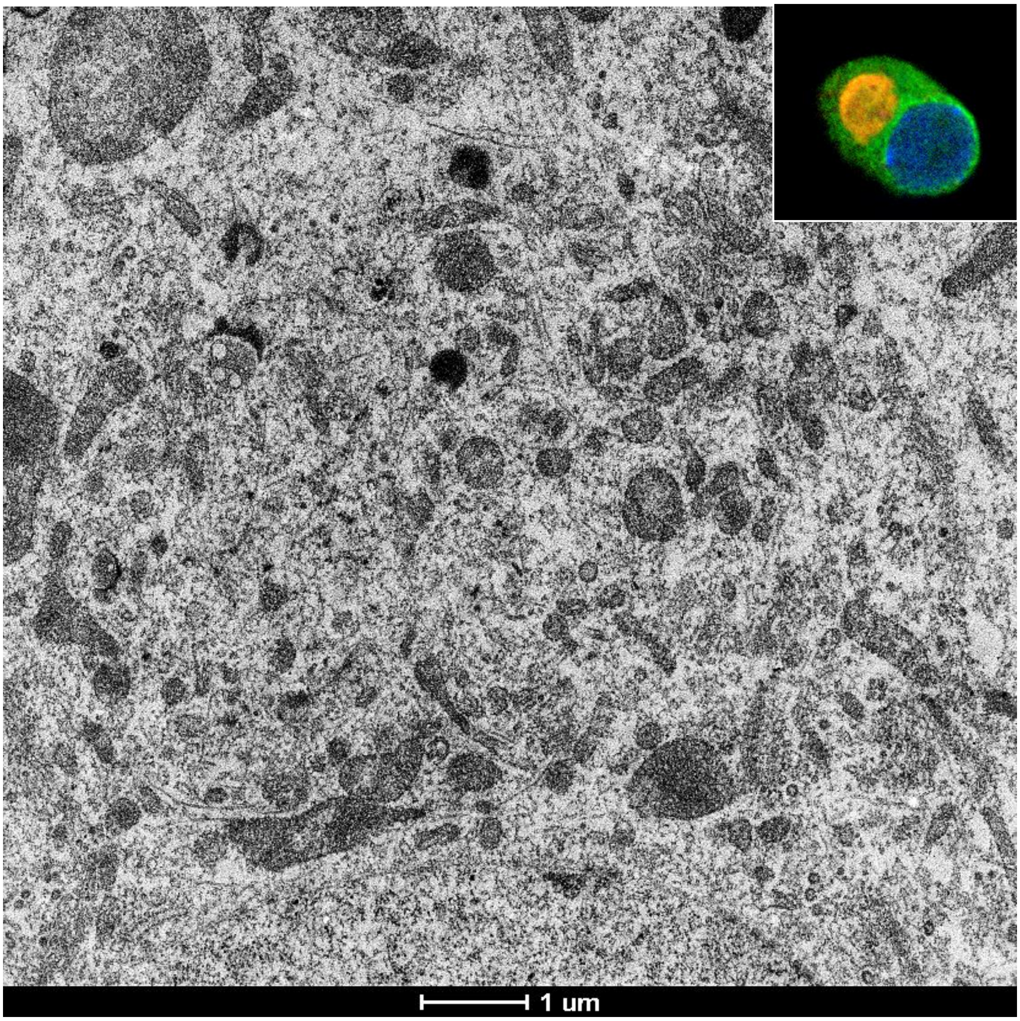


G

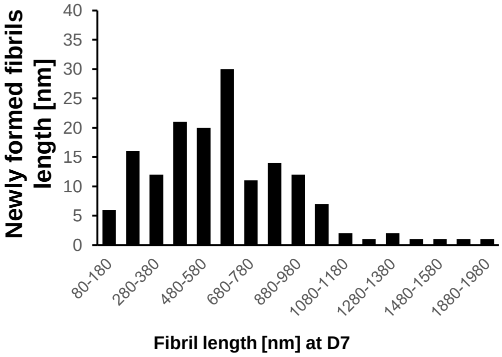


H

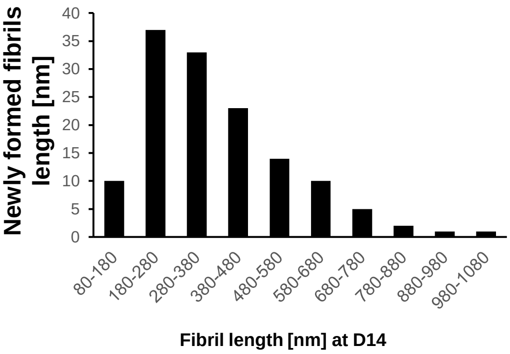


I

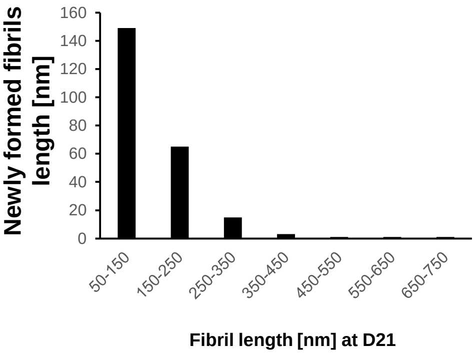


J

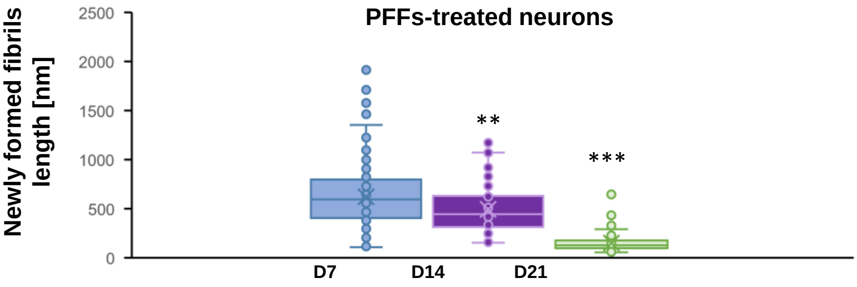

**Figure S6. Related to Figure 3**

A

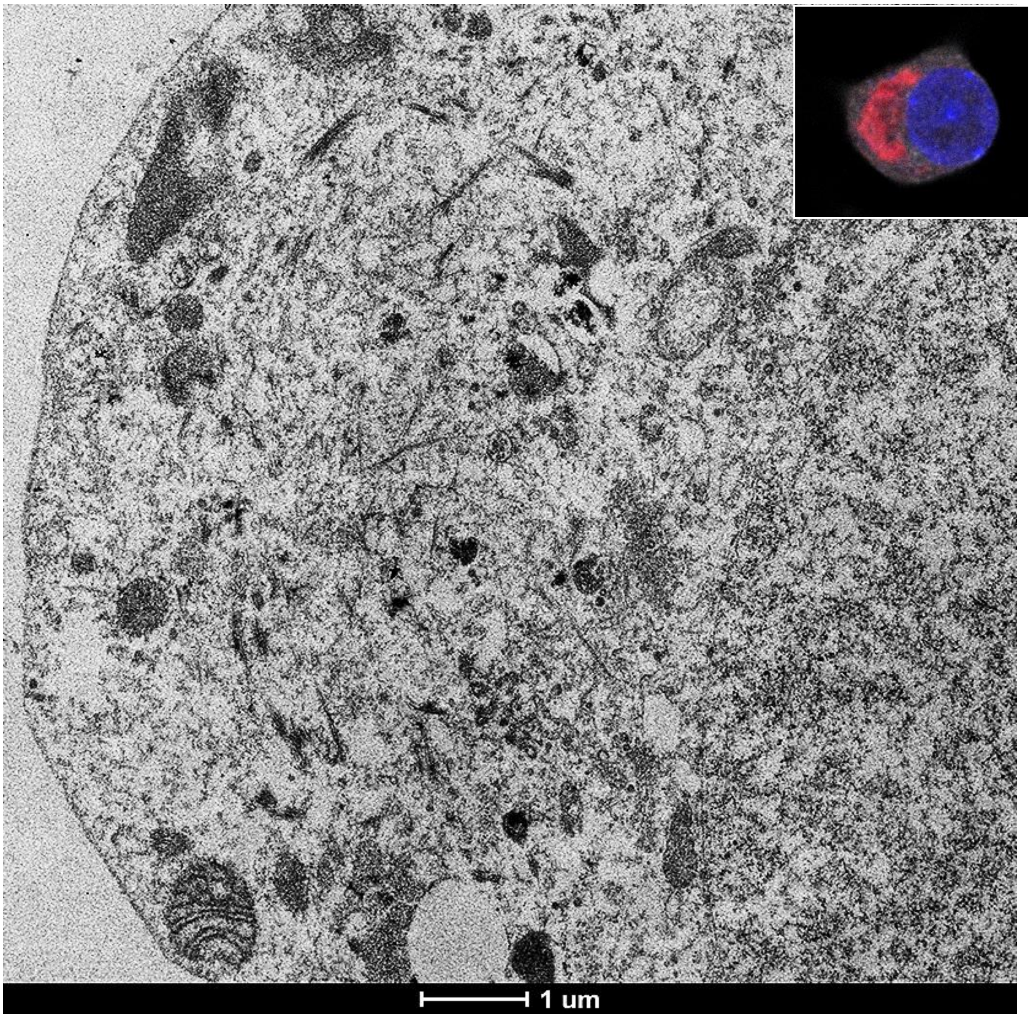


B

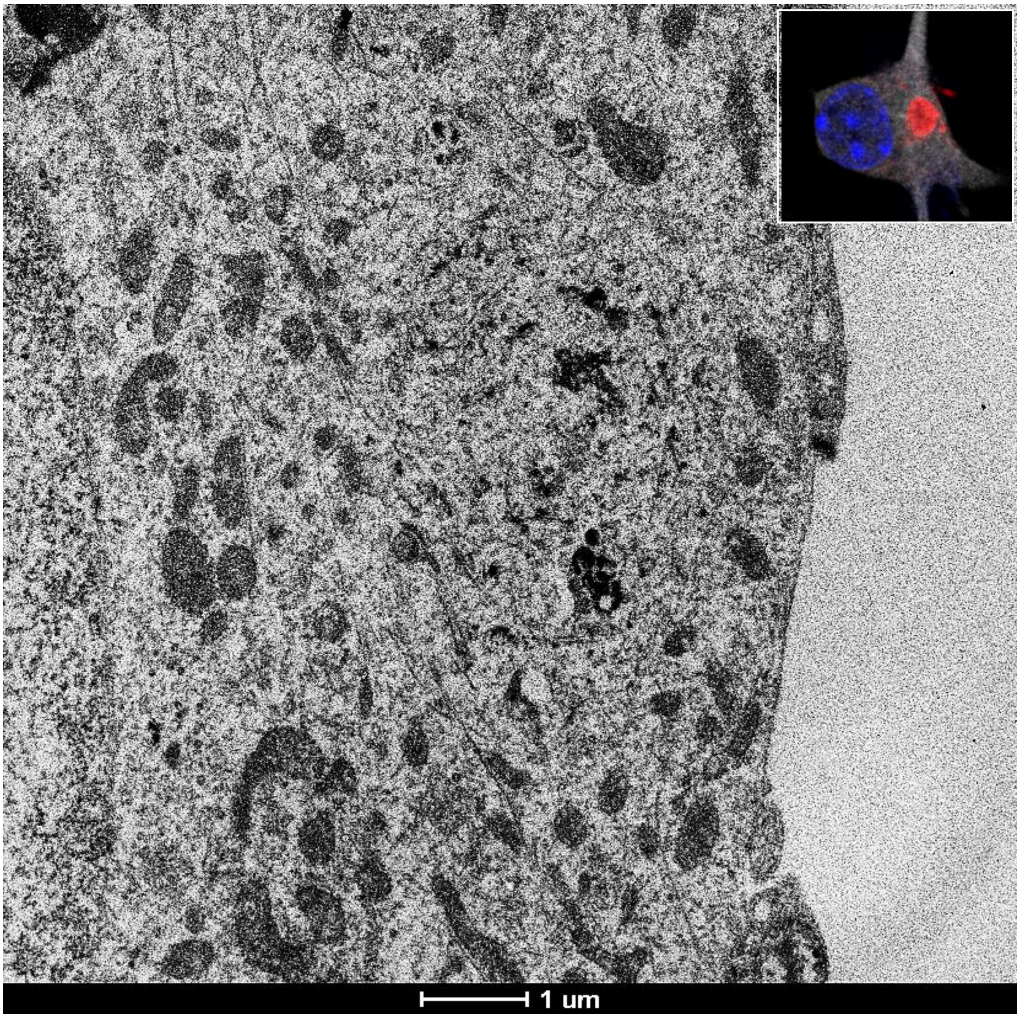

Figure S6. Related to Figure 3

C

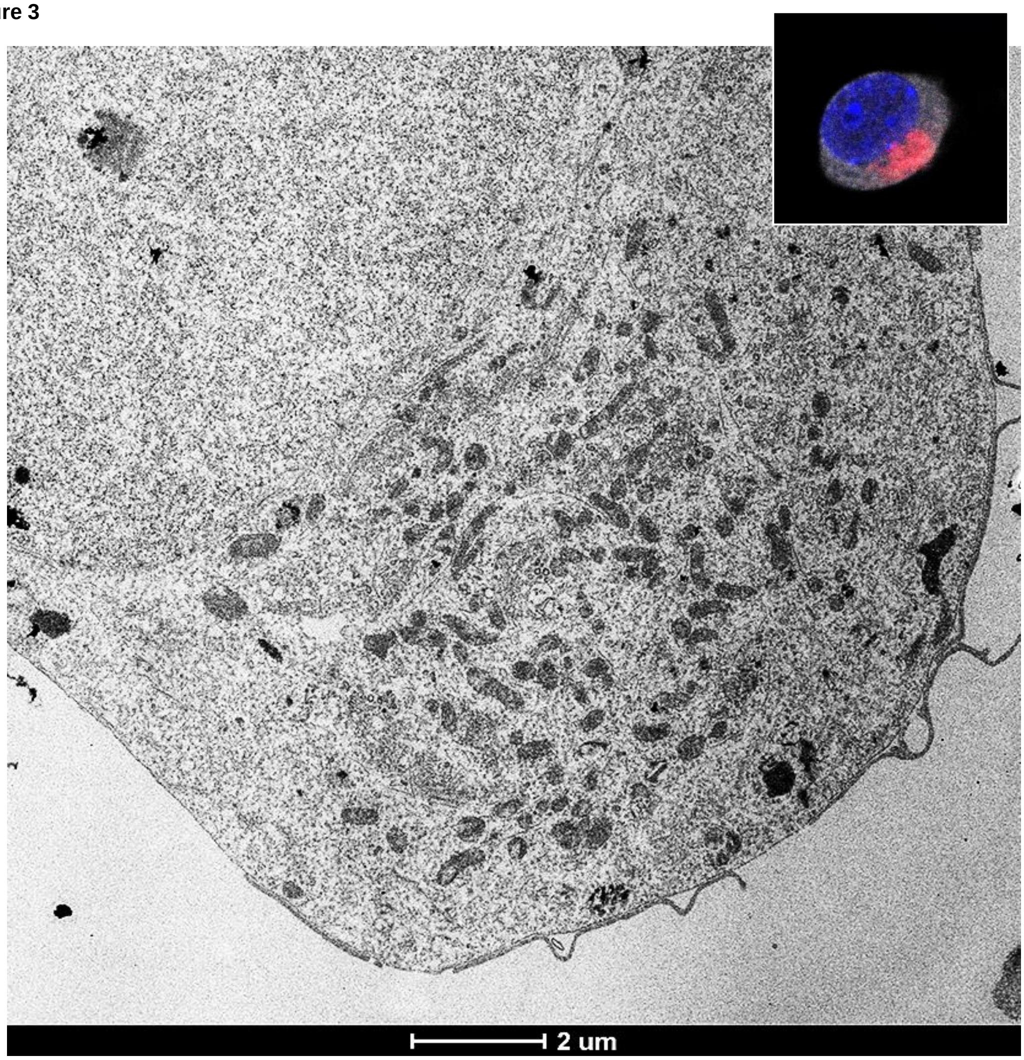


D

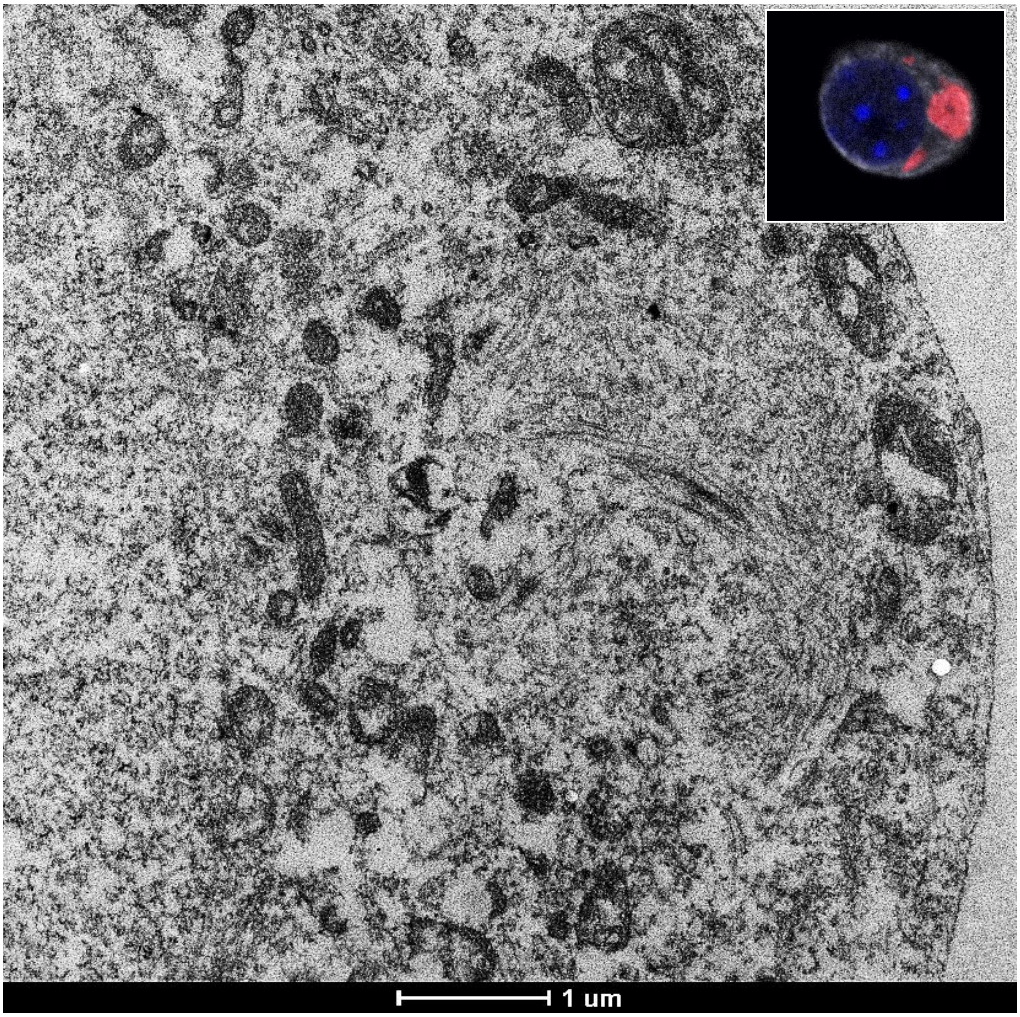

**Figure S7. Related to Figures 1-3**

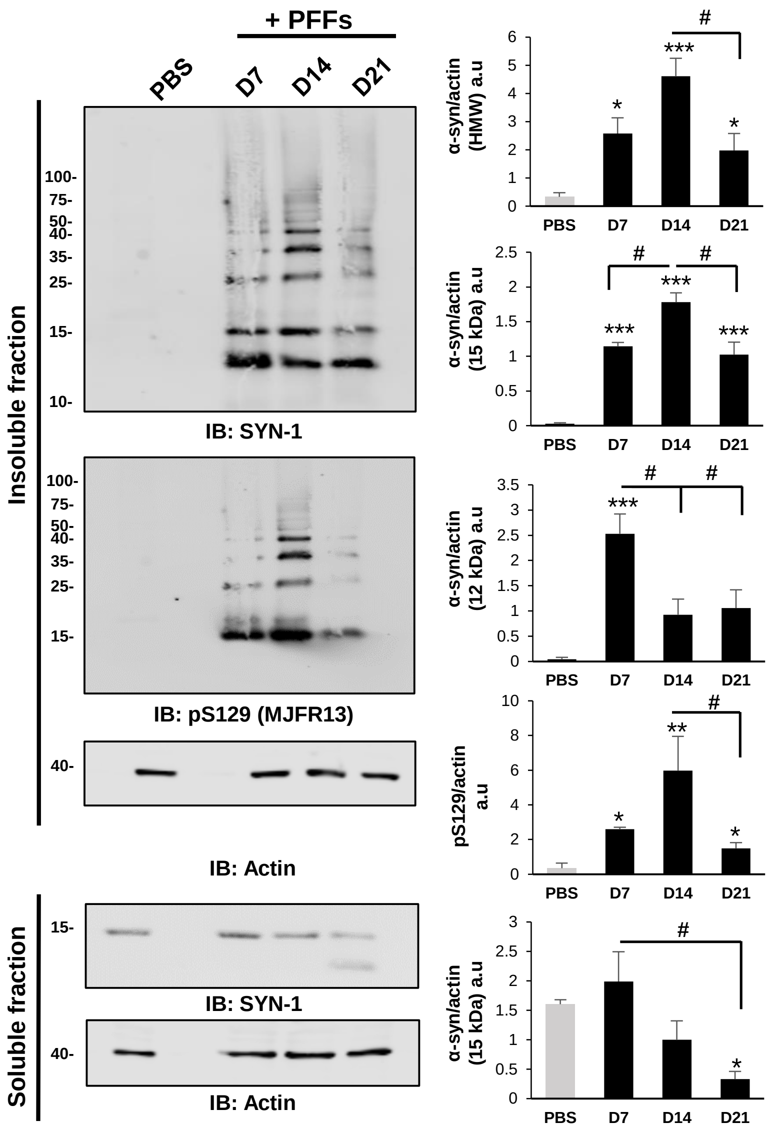

**Figure S8. Related to Figure 4**

**A**

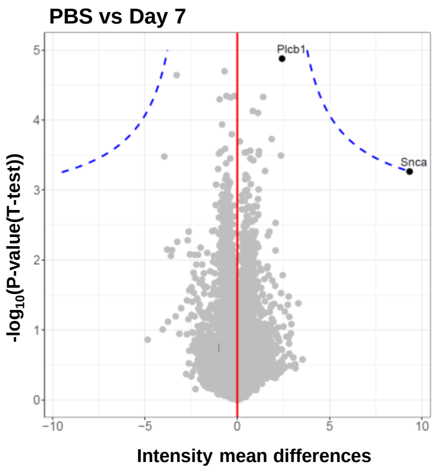


**B**

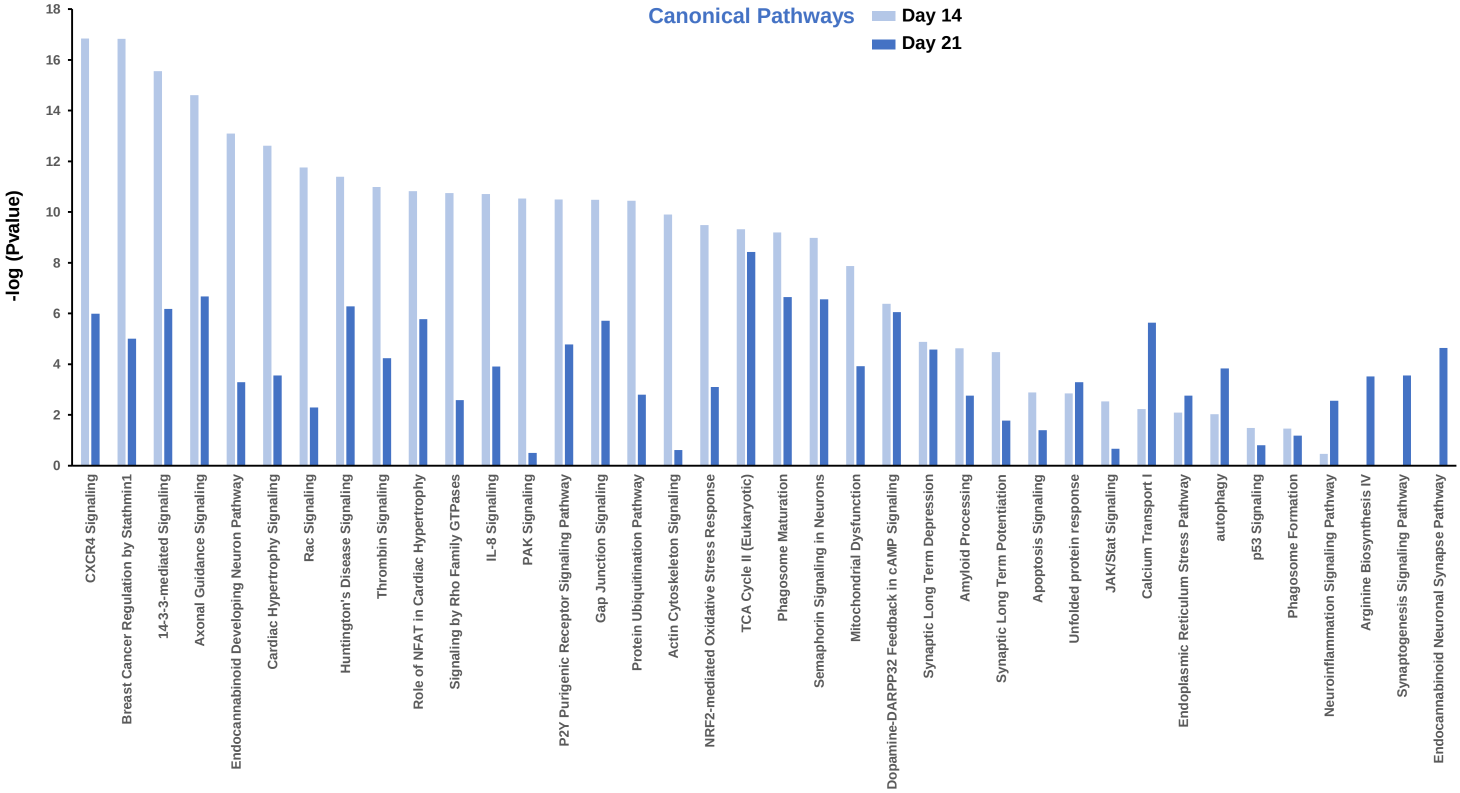


**C**

| Proteins name (Hendersen et al., 2017) | Insoluble Fraction (D14) | Insoluble Fraction (D21) |
|---|---|---|
| Collagen alpha-1(XII) chain | + | + |
| MAP/microtubule affinity-regulating kinase 1 | - | - |
| p21-activated kinase | - | - |
| α-synuclein | + | + |
| TBC1 domain family, member 10 | + | + |
| β-synuclein | - | - |
| Sequestosome-1 | + | + |
| Ubiquitin C | + | |
| Phospholipase C, β1 | + | + |
| HECT and RLD domain containing E3 ubiquitin protein ligase family member 1 | - | - |
| Hectd1 HECT domain containing 1 | + | + |
| Transformation-related protein 53 binding protein 1 | + | + |

**Figure S9. Related to Figure 5**

**A**

| Differentially-expressed genes (FDR cutoff of adjusted p value <0.01) | | | |
|---|---|---|---|
| Gene count | Total genes | Up-regulated | Down-Regulated |
| PBS vs D7 | 75 | 27 | 48 |
| PBS vs D14 | 435 | 106 | 223 |
| PBS vs D21 | 1017 | 455 | 562 |
| D14 vs D21 | 194 | 83 | 111 |

**B**

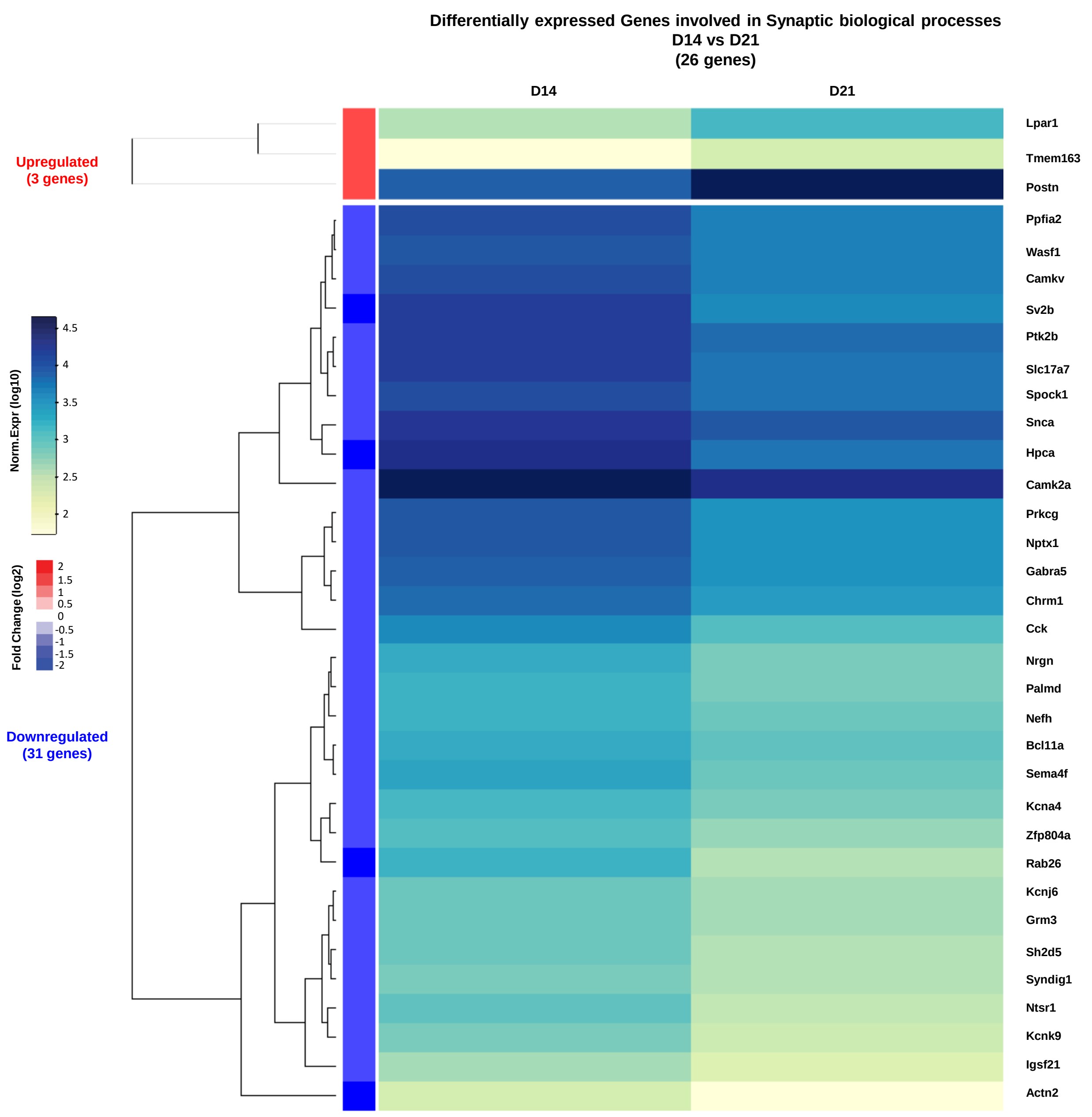

**Figure S10. Related to Figure 8**

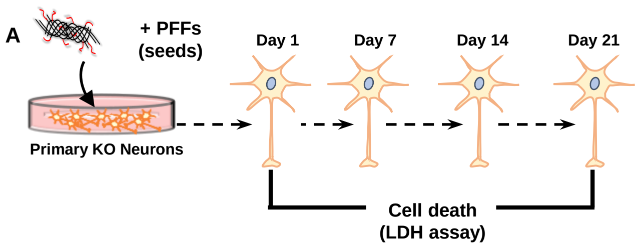


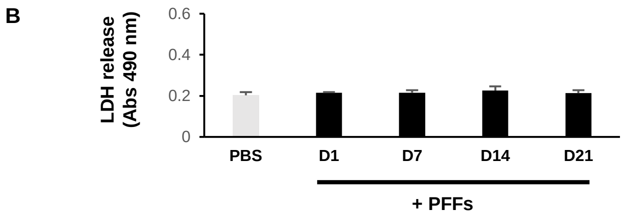

#### Figure S11. Related to Figure 4 and the discussion

| Proteins name | Insoluble Fraction (D14) | Insoluble Fraction (D21) | References |
|---|---|---|---|
| acyl-CoA synthetase long-chain family member 4 isoform 1 | + | + | Xia, Q., et al. (2008) |
| aggresome related proteins | + | + | Wakabayashi, K., et al. (2013) |
| a2-macroglobulin | + | + | Wakabayashi, K., et al. (2013) |
| amphiphysin isoform 1 | - | + | Xia, Q., et al. (2008) |
| a-synuclein | + | + | Wakabayashi, K., et al. (2013), Xia, Q., et al. (2008); Leverenz, J. B., et al. (2007) |
| Amyloid precursor protein | + | + | Wakabayashi, K., et al. (2013) |
| atlastin | + | + | Xia, Q., et al. (2008) |
| band 4.1-like protein | + | - | Leverenz, J. B., et al. (2007) |
| C terminus of Hsp70-interacting protein | + | - | Wakabayashi, K., et al. (2013) |
| Calcium/calmodulin-dependent protein kinase II | - | + | Wakabayashi, K., et al. (2013) |
| calnexin | + | + | Xia, Q., et al. (2008) |
| calreticulin precursor | - | + | Xia, Q., et al. (2008) |
| cathepsin D preproprotein | - | + | Xia, Q., et al. (2008) |
| coatomer protein complex | + | + | Xia, Q., et al. (2008) |
| coatomer protein complex, subunit beta | + | + | Xia, Q., et al. (2008) |
| coatomer protein complex, subunit zeta 1 | + | + | Xia, Q., et al. (2008) |
| cofilin 1 | + | - | Xia, Q., et al. (2008) |
| coronin, actin binding protein, 1A | + | - | Xia, Q., et al. (2008) |
| crystallin mu | + | + | Xia, Q., et al. (2008) |
| Cyclin-dependent kinase 5 | + | + | Wakabayashi, K., et al. (2013) |
| cysteine-rich protein 2 | + | + | Xia, Q., et al. (2008) |
| cytosolic nonspecific dipeptidase (EC 3.4.13.18) | - | + | Xia, Q., et al. (2008) |
| Cytochrome c | + | + | Wakabayashi, K., et al. (2013) |
| dihydropyrimidinase-like 2 | + | + | Xia, Q., et al. (2008) and Leverenz, J. B., et al. (2007) |
| Di-Ras2 | + | + | Xia, Q., et al. (2008) |
| doublecortin and CaM kinase-like 1 | + | + | Xia, Q., et al. (2008) |
| dynamin 3 | - | + | Leverenz, J. B., et al. (2007) |
| dynein, cytoplasmic, heavy polypeptide 1 | + | + | Xia, Q., et al. (2008) and Leverenz, J. B., et al. (2007) |
| dynein, cytoplasmic, intermediate polypeptide 1 | + | + | Xia, Q., et al. (2008) |
| dynein, cytoplasmic, light intermediate polypeptide 2 | + | + | Xia, Q., et al. (2008) |
| electron-transferring-flavoprotein dehydrogenase | + | + | Xia, Q., et al. (2008) |
| elongation factor 1 α2 | + | - | Leverenz, J. B., et al. (2007) |
| eukaryotic translation elongation factor 2 | + | - | Xia, Q., et al. (2008) |
| exportin 1 | + | - | Xia, Q., et al. (2008) |
| fascin 1 | + | + | Xia, Q., et al. (2008) and Leverenz, J. B., et al. (2007) |
| fatty acid synthase | + | + | Xia, Q., et al. (2008) |
| G-protein-coupled receptor kinase 5 | - | + | Wakabayashi, K., et al. (2013) |
| GATE-16/GABARAPL1 | + | - | Wakabayashi, K., et al. (2013) |
| glutamate dehydrogenase 1 | + | + | Xia, Q., et al. (2008) |
| glutamate-cysteine ligase regulatory protein | - | + | Xia, Q., et al. (2008) |
| glutathione S-transferase | + | + | Xia, Q., et al. (2008) |
| GSK3 b (glycogen synthase kinase 3b) | + | + | Wakabayashi, K., et al. (2013) |
| Heat-shock proteins 27 | - | - | Wakabayashi, K., et al. (2013) |
| Heat-shock proteins 40 | - | - | Wakabayashi, K., et al. (2013) |
| Heat-shock proteins 60 | + | + | Wakabayashi, K., et al. (2013) |
| Heat-shock proteins 70 | + | - | Wakabayashi, K., et al. (2013) |
| Heat-shock proteins 90 | + | + | Wakabayashi, K., et al. (2013) and Xia, Q., et al. (2008) |
| Heat-shock proteins 110 | - | - | Wakabayashi, K., et al. (2013) |
| Heme oxygenase | - | + | Wakabayashi, K., et al. (2013) |
| HDAC 6 | + | + | Wakabayashi, K., et al. (2013) |
| importin 7 | + | - | Xia, Q., et al. (2008) |
| kinesin family member 3A | + | + | Xia, Q., et al. (2008) |
| LIM and SH3 protein 1 | + | - | Xia, Q., et al. (2008) |
| Microtubule-associated protein 1 | + | - | Wakabayashi, K., et al. (2013) and Xia, Q., et al. (2008) |
| Microtubule-associated protein 1B | + | - | Wakabayashi, K., et al. (2013) and Xia, Q., et al. (2008) |
| Microtubule-associated protein 1A | + | - | Wakabayashi, K., et al. (2013) and Xia, Q., et al. (2008) |
| mitofusin 2 | + | - | Xia, Q., et al. (2008) |
| mitogen-activated protein kinase 1 | + | + | Xia, Q., et al. (2008) |
| NEDD8 | + | - | Wakabayashi, K., et al. (2013) |
| neuronal membrane glycoprotein | - | + | Leverenz, J. B., et al. (2007) |
| nuclear receptor binding protein | - | + | Xia, Q., et al. (2008) |
| OTU domain, ubiquitin aldehyde binding 1 | + | - | Xia, Q., et al. (2008) |
| P450 (cytochrome) oxidoreductase | + | - | Xia, Q., et al. (2008) |
| peroxiredoxin 5, mitochondrial precursor | + | - | Leverenz, J. B., et al. (2007) |
| phosphatidylinositol-4-phosphate 5-kinase type II beta isoform a | + | - | Xia, Q., et al. (2008) |
| phosphoglycerate mutase 1 (brain) | + | - | Xia, Q., et al. (2008) |
| Phospholipase C b1 | + | + | Wakabayashi, K., et al. (2013) |
| platelet-activating factor acetylhydrolase, isoform Ib, alpha subunit (45kD) | + | + | Xia, Q., et al. (2008) |
| p62/sequestosome 1 | + | + | Wakabayashi, K., et al. (2013) |
| profilin 1 | + | - | Xia, Q., et al. (2008) |
| progesterone receptor membrane component 1 | + | - | Xia, Q., et al. (2008) |
| Proteasome | + | + | Wakabayashi, K., et al. (2013) |
| Proteasome activators (PA700, PA28) | + | + | Wakabayashi, K., et al. (2013) |
| protein disulfide isomerase-related protein | + | + | Xia, Q., et al. (2008) |
| protein kinase C, beta 1 | + | + | Xia, Q., et al. (2008) |
| protein kinase C, gamma | + | + | Xia, Q., et al. (2008) |
| protein phosphatase 1, catalytic subunit, gamma isoform | + | - | Xia, Q., et al. (2008) |
| regulator of nonsense transcripts 1 | - | + | Xia, Q., et al. (2008) |
| reticulon 3 isoform a | + | + | Xia, Q., et al. (2008) |
| rho-associated protein kinase 2 | + | - | Leverenz, J. B., et al. (2007) |
| ROC1 (E3 ubiquitin-protein ligase RBX1) | + | - | Wakabayashi, K., et al. (2013) |
| SEC23-related protein A | + | - | Xia, Q., et al. (2008) |
| similar to aminopeptidase puromycin sensitive | - | + | Xia, Q., et al. (2008) |
| SLC25A3 | + | + | Leverenz, J. B., et al. (2007) |
| Sodium/potassium-transporting ATPase α1 chain | - | + | Leverenz, J. B., et al. (2007) |
| Sodium/potassium-transporting ATPase β1 chain | - | + | Leverenz, J. B., et al. (2007) |
| solute carrier family 30 (zinc transporter), member 9 | + | + | Xia, Q., et al. (2008) |
| STIP1 homology and U-Box containing protein 1 | + | - | Xia, Q., et al. (2008) |
| succinate dehydrogenase complex, subunit B, iron sulfur (Ip) | + | - | Xia, Q., et al. (2008) |
| Superoxide dismutase 2 (Mn superoxide dismutase) | - | + | Xia, Q., et al. (2008) |
| synaptic vesicle glycoprotein 2 | + | + | Xia, Q., et al. (2008) |
| synaptotagmin | - | + | Xia, Q., et al. (2008) |
| triosephosphate isomerase 1 | + | - | Xia, Q., et al. (2008) |
| Tubulin α1 chain | + | + | Wakabayashi, K., et al. (2013) and Leverenz, J. B., et al. (2007) |
| TUBB β Tubulin 1 | + | + | Leverenz, J. B., et al. (2007) |
| Tubulin β | + | + | Wakabayashi, K., et al. (2013) and Leverenz, J. B., et al. (2007) and xia |
| Tubulin β2 chain | - | + | Leverenz, J. B., et al. (2007) |
| Tubulin β4 chain | + | + | Leverenz, J. B., et al. (2007) |
| Tubulin β5 | + | - | Leverenz, J. B., et al. (2007) |
| Tubulin β5 chain | + | - | Leverenz, J. B., et al. (2007) |
| βTubulin | + | + | Wakabayashi, K., et al. (2013) and Leverenz, J. B., et al. (2007) |
| γ Tubulin | + | + | Wakabayashi, K., et al. (2013) and Leverenz, J. B., et al. (2007) |
| Tubulin polymerization promoting protein/p25 | - | - | Wakabayashi, K., et al. (2013) |
| Ubiquitin | + | + | Wakabayashi, K., et al. (2013) and Xia, Q., et al. (2008) |
| Ubiquitin conjugating enzyme UbcH7 (E2) | + | - | Wakabayashi, K., et al. (2013) |
| Ubiquitin C-terminal hydrolase | + | + | Wakabayashi, K., et al. (2013) |
| vacuolar protein sorting 33B | + | + | Xia, Q., et al. (2008) |
| vacuolar protein sorting 35 | + | + | Xia, Q., et al. (2008) |
| vesicle trafficking protein sec22b | + | - | Xia, Q., et al. (2008) |
| voltage dependent anion channel 1 | - | + | Leverenz, J. B., et al. (2007) |

#### Figure S12. Related to Figure 4 and the discussion

| Protein degradation machineries | Proteins name | Insoluble Fraction (D14) | Insoluble Fraction (D21) |
|---|---|---|---|
| **Ubiquitin Proteasome Pathway** | Anaphase-promoting complex subunit 5 | + | - |
| | Cell division cycle protein 23 homolog | + | - |
| | Cullin-2 | + | - |
| | Cullin-3 | - | + |
| | Cullin-5 | - | + |
| | Cullin-associated NEDD8-dissociated protein 1 | + | + |
| | E2/E3 hybrid ubiquitin-protein ligase UBE2O | - | + |
| | E3 ubiquitin-protein ligase HECTD1 | + | + |
| | E3 ubiquitin-protein ligase MARCH5 | + | - |
| | E3 ubiquitin-protein ligase NEDD4 | + | + |
| | E3 ubiquitin-protein ligase RBX1 | + | + |
| | E3 ubiquitin-protein ligase RNF14 | + | - |
| | E3 ubiquitin-protein ligase RNF126 | - | + |
| | E3 ubiquitin-protein ligase TRIM32 | + | - |
| | NEDD8-activating enzyme E1 catalytic subunit | + | - |
| | NEDD8-activating enzyme E1 regulatory subunit | - | + |
| | NEDD8-conjugating enzyme Ubc12 | + | - |
| | Proteasome-associated protein ECM29 homolog | + | + |
| | Proteasome complex activator | + | + |
| | SUMO-activating enzyme subunit 1 | + | + |
| | SUMO-activating enzyme subunit 2 | + | + |
| | SUMO-conjugating enzyme UBC9 | + | - |
| | Transcription elongation factor B polypeptide 1 | + | + |
| | Transcription elongation factor B polypeptide 2 | + | + |
| | Ubiquitin C-terminal hydrolase isozyme L1 UCHL1 | + | - |
| | Ubiquitin C-terminal hydrolase 5 USP5 | + | - |
| | Ubiquitin C-terminal hydrolase FAF-X USP9x | + | + |
| | Ubiquitin C-terminal hydrolase 14 USP14 | + | - |
| | Ubiquitin Cl-terminal hydrolase 15 USP15 | + | - |
| | Ubiquitin C-terminal hydrolase 47 USP47 | + | - |
| | Ubiquitin conjugating enzyme UbcH7 | + | - |
| | Ubiquitin-conjugating enzyme E2 D2 | + | + |
| | Ubiquitin-conjugating enzyme E2 D3 | + | - |
| | Ubiquitin-conjugating enzyme E2 E2 | - | + |
| | Ubiquitin-conjugating enzyme E2 L3 | + | - |
| | Ubiquitin-conjugating enzyme E2 O | + | - |
| | Ubiquitin-conjugating enzyme E2Q-like protein 1 | - | + |
| | Ubiquitin-like modifier-activating enzyme 1 | + | + |
| | Ubiquitin-like modifier-activating enzyme 1 | - | + |
| | Ubiquitin-like modifier-activating enzyme 6 | + | - |
| | Ubiquitin-protein ligase E3B | + | - |
| **Autophagy** | Autophagy-related protein 9 | + | - |
| | Cathepsin D | - | + |
| | Cathepsin F | - | + |
| | Lysosome-associated membrane glycoprotein 1 | - | + |
| | Microtubule-associated proteins 1A/1B light chain 3A – LC3 A | + | - |
| | Next to BRCA1 gene 1 protein | - | + |
| | Sequestosome-1 | + | + |
| | Serine/threonine-protein kinase mTOR | - | + |
| | Ubiquitin-like modifier-activating enzyme ATG7 | + | - |
| | Vacuolar protein sorting-associated protein 16 homolog | - | + |
| | Vacuolar protein sorting-associated protein 33A | + | - |
| | Vacuolar protein sorting-associated protein 33B | + | + |
| **Aggresome related proteins** | Histone deacetylase 6 | + | + |
| | Dynein, cytoplasmic, heavy chain | + | + |
| | Dynein, cytoplasmic, intermediate chain | + | + |
| | Microtubule-associated protein 1B | + | - |
| | Microtubule-associated proteins 1A/1B light chain 3A | + | - |
| | Regulator of microtubule dynamics protein 1 | - | + |

| Protein degradation machineries | Proteins name | Insoluble Fraction (D14) | Insoluble Fraction (D21) |
|---|---|---|---|
| **Unfolded Protein Response** | 78 kDa glucose-regulated protein | + | + |
| | Calnexin | + | + |
| | Calreticulin | - | + |
| | Endoplasmin | + | + |
| | Heat shock 70 kDa protein 4 | + | - |
| | Mitogen-activated protein kinase 8 | + | + |
| | Protein disulfide-isomerase | - | + |
| | Protein disulfide-isomerase A6 | + | - |
| | Protein sel-1 homolog 1 | + | - |
| | Transitional endoplasmic reticulum ATPase | + | + |
| **Chaperones** | 78 kDa glucose-regulated protein Hsp5a | + | + |
| | DnaJ homolog subfamily C member 5 Dnajc5 | + | - |
| | DnaJ (Hsp40) homolog | - | + |
| | Endoplasmin Hsp90b1 | + | + |
| | Heat shock 70 kDa protein 12A Hspa12a | + | + |
| | Heat shock 70 kDa protein 13 Hspa13 | + | + |
| | Heat shock 70 kDa protein 4 Hspa4 | + | - |
| | Heat shock protein 60 kDa, mitochondrial Hspd1 | + | + |
| | Heat shock protein HSP 90-alpha Hsp90aa1 | + | - |
| | Heat shock protein HSP 90-beta Hsp90ab1 | + | + |
| | Histone chaperone ASF1A | - | + |
| | Mitochondrial chaperone BCS1 | - | + |
| | Proteasome assembly chaperone 1 | + | - |
| | Tubulin-specific chaperone E | + | + |
| **Sumoylation Pathway** | C-terminal-binding protein 1 | - | + |
| | Dual specificity mitogen-activated protein kinase kinase 4 | + | + |
| | Mitochondrial Rho GTPase 1 | + | - |
| | Mitogen-activated protein kinase 10 | + | - |
| | Mitogen-activated protein kinase 8 | + | + |
| | Rab GDP dissociation inhibitor alpha | - | + |
| | Rab GDP dissociation inhibitor beta | + | + |
| | Rab GDP dissociation inhibitor beta | + | - |
| | Rho-related GTP-binding protein RhoB | + | - |
| | SUMO-activating enzyme subunit 1 | + | + |
| | SUMO-activating enzyme subunit 2 | + | + |
| | SUMO-conjugating enzyme UBC9 | + | - |
| **Endolysosomal Pathway** | Argininosuccinate synthase | - | + |
| | Calpain-2 catalytic subunit | + | + |
| | Cathepsin D | - | + |
| | Cathepsin F | - | + |
| | CD63 antigen | - | + |
| | Chitinase domain-containing protein 1 | - | + |
| | Lysosomal acid phosphatase | - | + |
| | Lysosome membrane protein 2 | - | + |
| | Lysosome-associated membrane glycoprotein 1 | - | + |
| | Membrane protein MLC1 | - | + |
| | Next to BRCA1 gene 1 protein | - | + |
| | Ragulator complex protein LAMTOR2 | + | - |
| | Ragulator complex protein LAMTOR3 | + | - |
| | Ras-related GTP-binding protein A | - | + |
| | Ras-related protein Rab-7a | + | - |
| | Sequestosome-1 | + | + |
| | Serine/threonine-protein kinase mTOR | - | + |
| | Vacuolar fusion protein CCZ1 homolog | - | + |
| | Vacuolar protein sorting-associated protein 16 homolog | - | + |
| | Vacuolar protein sorting-associated protein 33B | + | + |
| | Vacuolar protein sorting-associated protein 35 | + | + |

**Figure S13. Related to Discussion**

#### Day 7

| | | |
|---|---|---|
| **Subcellular localization** | Neurites | +++ |
| | Cell bodies | + |
| **LB markers** | p62 (autophagy) and ub (proteasome) | √ |
| **α-syn PTMs** | pS129 | √ |
| | C-ter truncation | x |
| **Ultrastructure** | Fibrils length | Long filaments (length: 600 nm to 2μm) |
| | Lateral association and packing | x |
| | Organelles interaction | x |
| **Transcriptomic** | Differentially-expressed genes | 75 genes |
| **LB Proteome** | Enrichment of the insoluble fraction | Only 2 proteins |
| **Biological Processes** | Mitochondria dysfunctions | x |
| | Synaptic impairment | x |
| | Cell death | x |

#### Day 14

| | | |
|---|---|---|
| **Subcellular localization** | Neurites | ++ |
| | Cell bodies | ++ |
| **LB markers** | p62 (autophagy) and ub (proteasome) | √ |
| **α-syn PTMs** | pS129 | √ |
| | C-ter truncation | √ |
| **Ultrastructure** | Fibrils length | Shorter filaments (~450 nm) |
| | Lateral association and packing | √ |
| | Organelles interaction | √ |
| **Transcriptomic** | Differentially-expressed genes | 435 genes |
| **LB Proteome** | Enrichment of the insoluble fraction | 633 proteins |
| **Biological Processes** | Mitochondria dysfunctions | x |
| | Synaptic impairment | √ (early event – reduction pre-synaptic area) |
| | Cell death | √ (early event – caspase 3 activation) |

#### Day 21

| | | |
|---|---|---|
| **Subcellular localization** | Neurites | ++ |
| | Cell bodies | +++ |
| **LB markers** | p62 (autophagy), ub (proteasome) and lipids | √ |
| **α-syn PTMs** | pS129 | √ |
| | C-ter truncation | √ |
| **Ultrastructure** | Fibrils length | Short filaments (~300 nm) |
| | Lateral association and packing | √ |
| | Organelles interaction and sequestration | √ |
| **Transcriptomic** | Differentially-expressed genes | 1017 genes |
| **LB Proteome** | Enrichment of the insoluble fraction | 568 proteins |
| **Biological Processes** | Mitochondria dysfunctions ↑<br>-Respiration ↓<br>-Mitochondria membrane potential ↓<br>-Mitochondrial protein levels ↓ | √ |
| | Synaptic impairment ↑<br>-Neurites ↓<br>-Pre-synaptic area ↓<br>-Synaptic plasticity ↓<br>-Pre and Post synaptic protein levels ↓ | √ |
| | Cell death (late events – loss of plasma membrane integrity) | √ |

**Figure S14. Related to Figure 6 and Discussion**

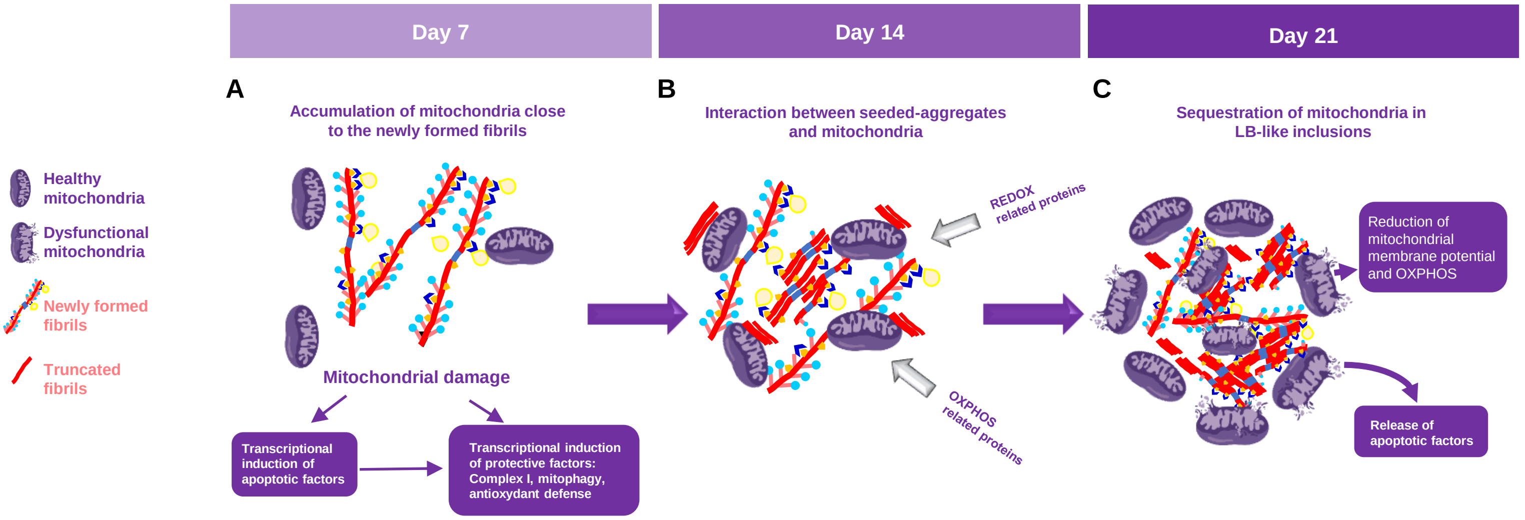


| Stage | | D7: Cellular responses related to mitochondrial damage during the process of fibrils formation | D14: Interaction of mitochondrial and antioxidant defense components with α-syn seeded aggregates | D21: Mitochondrial sequestration into LB-like inclusions induces mitochondrial dysfunctions and cell death |
|---|---|---|---|---|
| Evidence for aggregate-mitochondria interaction | ICC | Moderate colocalization of pS129 positive aggregates with TOM20, TIM23 and VDAC1 | Strong colocalization of pS129 positive aggregates with TOM20, TIM23 and VDAC1 | Strong colocalization of pS129 positive aggregates with TOM20, TIM23 and VDAC1 |
| | EM | Mitochondria accumulate in proximity to cytoplasmic, fibrillar aggregates (not in the neuritic ones) | Mitochondria interact with fibrillar aggregates | Mitochondria get incorporated into maturing LB-like inclusions |
| | proteomics | No mitochondrial proteins detected in insoluble fraction of PFF-treated neurons | Many mitochondrial components, such as OXPHOS proteins and MFN2 are enriched in the insoluble fraction of PFF-treated neurons | Reduction of mitochondrial components found in the insoluble fraction of PFF-treated neurons as compared to D14 |
| Observations | | a. Transcriptional upregulation (until D21) of antioxidant defense systems, mitochondrial quality control, OXPHOS<br>b. Reduced mitochondrial ROS production.<br>c. Maintenance of OXPHOS performance and mitochondrial membrane potential | a. Sequestration of OXPHOS and antioxidant defense components<br>b. Mitochondrial ROS back to baseline level<br>c. Despite compact aggregate-organelle associations OXPHOS performance and mitochondrial membrane potential remain high | a. Mitochondrial membrane potential reduced<br>b. Heavily impaired OXHOS capacities<br>c. Reduced levels of CI-NDUFB8, OPA1 and MFN2 |
| Interpretation | | a. Aggregates start to coordinate mitochondria in neuronal soma<br>b. Interaction is unstable<br>c. Neuronal damage signalling induces protective pathways counteracting mitochondrial dysfunction | a. Oxidation and other associated PTMs stabilize aggregates, immobilizing interacting mitochondria<br>b. Enhanced fibril growth on membraneous substrate provided by mitochondrial envelop | a. Dynamics of mitochondria-enclosing aggregates mechanically disrupt membranes → reduced mitochondrial membrane potential and release of apoptotic factors<br>b. Accumulating defective mitochondria surpass neuronal clearance capacities |
